# Eggplant’s foliar chlorogenic acid provides resistance against the tropical armyworm

**DOI:** 10.1101/2023.02.10.527964

**Authors:** Manish Kumar, K.P. Umesh, Prashasti P. Pandey, D. M. Firake, Sagar S Pandit

## Abstract

Lepidopteran pests are the major crop devastators. Farmers have to resort to heavy synthetic pesticide application for their control. It increases the pesticide residue contamination on produce and causes health hazards. Synthetic pesticides also endanger beneficial insects and pollute the environment. Therefore, the use of safe and eco-friendly botanicals as biopesticides is rapidly increasing. Despite their high demand, only a few botanicals are commercially available. Consequently, biopesticide discovery research boomed in the last decade.

*Spodoptera litura* Fabricius (armyworm) is a polyphagous multi-insecticide-resistant lepidopteran pest. It is a serious concern for several commercially important crops. In this study, we employed a chemical ecology approach to discover a biopesticide against it. As a biopesticide source, we explored secondary metabolite-rich *Solanum melongena* L. (eggplant), one of the armyworm’s hosts. We found that the armyworm larvae show differential occurrence on seven eggplant varieties; the Himalayan eggplant variety RC-RL-22 (RL22) showed no armyworm infestation. When reared in a no-choice condition on RL22, larval mortality was two-fold higher, and mass was three-fold lower than the varieties showing high infestation. We tested whether RL22’s secondary metabolite(s) were associated with this hampered larval performance. Using LC-ESI-QTOF-based non-targeted metabolomics of eggplant varieties, we identified candidate metabolites. 5-*O*-caffeoylquinic acid (chlorogenic acid; CGA) showed a strong negative correlation (r= -0.88; *p*= 0.008) with the larval performance. CGA-spiked (average physiological concentration) artificial diet (CGA-AD)-fed larvae showed a three-fold mass reduction and two-fold mortality increase than the control artificial diet (AD)-fed larvae; pupation and eclosion also significantly reduced (1.3-fold and 1.4-fold, respectively) in the CGA-ingested larvae. We used a reverse genetics approach to assess the *in planta* insecticidal potential of CGA. When RL22’s CGA biosynthesis gene hydroxycinnamoyl-CoA quinate transferase (*Sm*HQT) was silenced using virus-induced gene silencing (VIGS), CGA levels decreased by three-fold. This CGA depletion rendered RL22 two-fold armyworm-susceptible than controls. Foliar CGA application restored RL22’s armyworm resistance.

Overall, this study showed that CGA exhibits larvicidal properties against the armyworm. It is also safe for beneficial organisms. CGA is a well-known dietary supplement and an antioxidant for humans. Thus, it is safe for human consumption. Together, high CGA-containing varieties can be used to reduce the armyworm infestation risk. CGA is a promising biopesticide candidate for the field trial phase against the lepidopteran pests, especially armyworm. If successful, it can be integrated into the pest control measures.

## Introduction

Plant and insect herbivores’ interactions are complex and often chemically-based ^1^. Since insect herbivores heavily depend on plants for food and shelter, they pose a major threat to plants ^2-5^. Continuous feeding by the insect herbivores causes severe damage to plant productivity and fitness^6^. The co-evolution of plants and insects has led to the development of various defense mechanisms in the plant to deter herbivores and reduce their attack^7^. To reduce and sometimes to restrict the herbivores, plants have evolved physical barriers like cell wall thickening, thorns, prickles, and wax deposition, mainly which have limited deterring effects on insects. Plants have also evolved to produce secondary defense metabolites such as phytotoxins, which can deter even all those herbivores which can escape the physical barriers developed by plants^8-10^. Alkaloids, phenolics, terpenoids, and glycosylated phytoanticipins are the common plant defense metabolite classes^11,12^,

In the agriculture, insect pests cause heavy crop losses. These pests cause global annual agricultural loses over $100 billion^13^. To control pests, farmers often resort to the heavy use of synthetic pesticides (one or more in combination). The injudicious use of synthetic pesticides is hazardous to beneficial insects like pollinators and biocontrol agents. They are also hazardous to humans. Worldwide, ∼385 million unintentional severe pesticide poisoning cases including around 11,000 fatalities are reported^14^. Approximately 0.4 million tons of insecticides produced annually increase pesticide residue contamination and cause health hazards, which endangers beneficial insects and pollutes the environment^15-17^. Owing to the hazards of synthetic pesticides, the demand for biopesticides, the pest management agents based on living organisms or natural products, is increasing. Biopesticides promise pest control with much lesser hazards than synthetic pesticides. Therefore, research on biopesticides to reduce the synthetic pesticide load is being promoted. Plants being a rich source of secondary metabolites, plant-based biopesticide discovery became a prime focus area. Nicotine, one of the earliest known botanicals, was widely used as an insecticide^18^. Some commercially successful botanical insecticides are Tetranortriterpenoid azadirachtin from *Azadirachta indica* A. Juss (Neem)^19^, a polyhydroxylic diterpene ryanodine from *Ryania speciosa* Vahl (Ryania)^20^, oleoresin pyrethrum from *Tanacetum cinerariaefolium* Sch.Bip. (Dalmatian chrysanthemum)^21^, and isoflavone rotenone from *Derris, Lonchocarpus*, and *Tephrosia* species^22^. Several other secondary metabolites are likely to possess insecticidal properties; however, they have not been identified and commercialized.

Phenolics play important roles in protecting plants from ultraviolet radiation, herbivores, and pathogens^23^. Phenolic ingestion exerts oxidative stress in insects with alkaline gut pH^24^. 5-*O*-caffeoylquinic acid (chlorogenic acid; CGA)^25^ is a major phenolic found in the leaves of eggplant, which has been reported as a defense molecule against herbivores^26^ causes oxidative bursts in the insects^27^ and has antibacterial and antifungal activities^28,29^. Three alternate CGA biosynthesis pathways have been reported in the plants^28,30^. The most accepted pathway and common in Solanaceae plants is the shikimate pathway in which hydroxycinnamoyl-CoA quinate transferase (HQT) conjugates quinic acid (QNA) to caffeoyl-CoA to form CGA^30,31^. However, there is no detailed information available in the literature about the role of HQT in the biosynthesis of CGA and other derivative compounds in eggplant^28,32^.

Here, we explored the secondary metabolite-rich eggplant as a biopesticide source by employing a chemical ecology approach against *Spodoptera litura* Fabricius (Lepidoptera: Noctuidae), which is a gregarious, polyphagous multi-insecticide-resistant lepidopteran pest^33^. Based on various hosts and high fecundity rate, this pest is commonly known as a tropical armyworm, taro caterpillar, tobacco cutworm, cotton leafworm, and cluster caterpillar. It is a serious concern for several commercially important crops, including tomato, soybean, groundnut, cotton, tobacco, tomato, and eggplant. Its short life cycle, high fecundity on several hosts, larval host switching ability, and adults’ migratory habit contribute to its success as a polyphagous pest^12,34^. According to the global arthropod pesticide resistance database, armyworm larva has evolved resistance against almost all commercial pesticides and also against the Bt-toxin^35,36^. It has become one of the most pesticide-applied pests and, thus, a major cause of health and environmental hazards. Although the polyphagous armyworm frequently infests eggplant, the infestation is rarely severe. It often prefers other hosts, such as castor, cotton, soybean, groundnut, and cauliflower, over the eggplant, indicating that eggplant is an inauspicious host^37,38^. Moreover, several eggplant metabolites belonging to chemical classes like terpenoids, alkaloids, and phenolics are reported to show promising insecticidal activities^39^. This information renders eggplant an appropriate candidate for discovering a biopesticide against armyworm.

## Materials and Methods

### Field and plants

Seven different varieties of *Solanum melongena* (eggplant) were planted in the field in a randomized complete block design (RCBD) in a plot of 30×30 meters, which is divided into four equal plots of 15×15 meters. The eggplant varieties used for field plantation were: Himalayan eggplant variety (RL-22), Ankur Kavach (KV), JK Agro Hybrid-green long (JK), Omaxe-CVK MK (CVK), Ankur Vijay (VJ), L-Riccia-Hirvi Kateri (HK), KGN’s F1 Pinstripe (KP). Each plant was planted at a distance of 0.8 m from the other. Larvae’s natural occurrence data were collected from this field (n= 10 plants). Leaf tissues were collected from this field for metabolomics analyses. All harvested leaf tissues were flash-frozen in liquid nitrogen and were stored at -80°C until further processing.

### Armyworm and natural enemies

*Spodoptera litura* (armyworm) larvae were collected from the agricultural fields in and around Pune, India. Larvae were reared on the fresh *Ricinus communis* L. (castor) leaves to maintain the source culture. Separate cultures were maintained on the eggplant leaves and artificial diet (common bean powder and chickpea flour-based diet)^12,40^ for the experiments (Table 2). Eggs and early instar larvae were transferred to an artificial diet spiked with metabolites for the metabolite activity testing experiments. Adults were fed a 10% sucrose solution. Ad libitum feeding was ensured in all the cultures. All the cultures were maintained under controlled conditions (relative humidity: 65%, photoperiod: 16 h, temperature: 25± 1°C).

The eggplant field was surveyed to find the armyworm’s natural enemies, as described by Mitra *et al*.^34^. Based on abundance and ease of collection, maintenance, and bioassays, a total of eight predators (spiders, ants, and praying mantis) were selected for further experiments. They were fed (ad libitum) the first and second instar armyworm larvae (to ease their maintenance) and were maintained in the above-controlled conditions.

### Armyworm’s performance assays on eggplant leaves

To study the larval performance of eggplant leaves, eggs were hatched on the leaves of different eggplant varieties. Larval mass was measured every fifth day (n= 30). Larvae were supplied with fresh leaves, and ad libitum feeding was ensured until the pupation. Larval mortality was also recorded on every fifth day.

### Armyworm’s performance assays on the candidate metabolite-spiked artificial diets

To analyze the effects of candidate metabolites on neonate mortality, 100 neonates were placed on an artificial diet spiked with candidate metabolite CGA (five replicates, each containing 20 neonates). After 24h and 48h, mortality was recorded. Similarly, mortality was recorded in the neonates fed with artificial diets spiked with different concentrations of the candidate metabolite. Neonates were fed on artificial diets spiked with the following concentrations of the candidate metabolite, 100 μg/g of diet, 200 μg/g of diet, 400 μg/g of diet, 800 μg/g of diet, and 1 mg/g of diet.

To measure the larval performance on artificial diets spiked with different eggplant metabolites, neonates were fed on artificial diets spiked with candidate metabolites. Metabolites were freshly mixed with the artificial diet before every experiment. Larval mass was measured on every fifth day (n= 30). Armyworm eggs were inoculated on artificial diets spiked separately with CGA (650 µg/g; n= 50 for each treatment) to study the effects of eggplant-specific metabolites. Neonate mortality was recorded after 24h and 48h. Larval mass was also measured on the above diets every fifth day.

### Extraction of metabolites for UPLC/ESI/QTOF-based analysis

To extract metabolites from eggplant leaves, 1 ml of 70% methanol (spiked with 400 ng/ml formononetin as an internal standard for both positive and negative ionization-based data acquisition mode) was added to the 200 mg of homogenized leaf tissue. The mixture was vortexed, and the tissue debris was removed by centrifugation (13000 rpm, 10 min). The supernatant was incubated at -80°C overnight to precipitate high molecular weight lipids, which were removed by centrifugation (13000 rpm, 20 min, 4°C). Frass and hemolymph samples were extracted similarly, using 70% methanol in a ratio of 100 mg/ml and 10 µl*/*100ul, respectively. Metabolites from the other larval tissues, like gut and fat bodies, were extracted using 1 ml of 70% methanol (spiked with IS) per 100 mg of tissue.

### UPLC/ESI/QTOF-based data acquisition and non-targeted metabolomics

All the extracts were analyzed using the SCIEX-X500R-QTOF (UPLC/ESI-QTOF-MS). Samples were separated on a Phenomenex Gemini® C18 column (50×4.6 mm, 5μm, 118Å), using MilliQ water with 0.1% formic acid and methanol with 0.1% formic acid as mobile phase. A gradient of 0 min 5% B, 1 min 5% B, 10 min 95% B, 11 min 95% B, 12 min 5% B, and 15 min 5% B, with a flow rate of 0.5 ml/min, was used. Compounds were analyzed in both negative and positive ion modes (capillary voltage-4500 V, capillary exit-130 V, dry gas temperature-200°C).

For both qualitative and quantitative data analyses, SCIEX-OS software was used. To quantify the metabolite using peak area calculation, the MQ4 algorithm was used, and relative concentrations were calculated with reference to the internal standard (IS) formononetin (400 ng/ml added into extraction buffer). Parameters: minimum peak width-3 points, minimum peak heights-100, signal/noise integration threshold-3, XIC width-0.02 Da, Gaussian smooth width-1 point, Noise percentage-95%, baseline subtract window-0.5 min, peak splitting-2 points, RT half window-30 sec were used to select and calculate peak area of parent ion mass of each metabolite. Peak areas were further converted into concentrations (in units of a gram) using standard curves of peak area for those metabolites for which pure standards were procured. For metabolites whose pure standards were not available, the peak area of IS was used as a reference to calculate their relative concentrations across the samples. All the concentrations were calculated into µg/g or ng/g fresh mass (FM) of respective samples.

A non-targeted metabolomics-based approach was used to identify metabolites in eggplant leaves. MSDIAL (http://prime.psc.riken.jp/compms/msdial/main.html) was used for the compound annotation^41^. The algorithm of MSDIAL allows *in-silico* compound search and identification of unknown metabolites with reference to the MSP spectral kit containing EI-MS, MS/MS, and CCS values from public mass spectral databases (such as MassBank, ReSpect, MetaboBASE, Fiehn/Vaniya natural product library, GNPS, etc.). Raw data files (.wiff2) were directly imported as profile data. File deconvolution parameters were set as MS1 tolerance of 0.01 Da, MS2 tolerance of 0.025 Da, peak detection of 1000 amplitudes (minimum peak height), and mass slice width of 0.1 Da, respectively ^41,42^. Spectral data (MS/MS) from public MSP spectral kit available online on the RIKEN CompMS server (http://prime.psc.riken.jp/compms/msdial/main.html#MSP) were used for metabolites identification as [M-H]^-^ and [M+H]^+^ adducts separately. Identification parameters were set as MS1 tolerance of 0.01 Da, MS2 tolerance of 0.05 Da, and 80% identification score cut-off. Alignment parameters were set as a retention time tolerance of 0.05 min and an MS1 tolerance of 0.01 Da. A minimum of two references MS2 fragment matches were set as a cut-off to confirm the identity of the suggested metabolite^41,42^. Retention time, formula, and metabolite names were exported and used for the peak area calculation using SCIEX-OS 2.0 software, followed by the quantitation with the help of the internal standard (formononetin).

### Cloning of VIGS fragments into tobacco rattle virus-based *p*TRV2 vector

*Sm*HQT CDS sequence was retrieved from the eggplant genome database (http://eggplant.kazusa.or.jp/) based on the BLAST and multiple sequence alignment of reported HQTs from other plants with the candidate sequences^43^; A phylogenetic tree was constructed using the neighbor-joining (Dayhoff matrix) algorithm (1000 bootstrap replicates) with the MEGA7 software. To clone the *Sm*HQT gene fragment into the *p*TRV2 VIGS vector, primers (Table S1) were designed to amplify the 351 bp region (towards the 3’ end) of the *Sm*HQT mRNA sequence, which was cloned into the *p*TRV2 vector (kindly received from Professor Sir David Baulcombe, University of Cambridge). Cloning was confirmed by sequencing the recombinant vector, hereafter referred to as pTRV2-*Sm*HQT.

For the VIGS, eggplant seeds were germinated in the sterilized field soil in the 4× 4× 11 (l× b× h) inch pots. Plants were maintained in the climate chamber (relative humidity: 70-80%, photoperiod: 16 h light, temperature: 22± 1°C). The 4-5 leaf-stage plants were used for the VIGS procedure, as described by Ghosh *et al*.^44^.

Primary cultures of *A. tumefaciens* containing *p*TRV1, *p*TRV2, *p*TRV2-*Sm*HQT, pTRV2-*Sl*CE4, and pTRV2-*Sl*CE4 were grown at 27°C for 16 hours in the YEP medium containing three antibiotics (25 μg/mL rifampicin and 50 μg/mL kanamycin). Secondary cultures were generated by inoculating 500uL of the respective primary cultures in 100mL YEP medium containing antibiotics, 10 mM MES, and 20 μM acetosyringone. These were grown at 27°C with 200 rpm shaking until their OD_600_ reached 2.0. Cells were harvested by centrifugation and were resuspended in the infiltration buffer (10 mM MgCl_2_, 10 mM MES [2-(4-Morpholino)-Ethane Sulfonic Acid], 200 μM acetosyringone (3,5-Dimethoxy-4-hydroxy-acetophenone), pH 5.6). These cultures were infiltrated in the eggplants as per the procedure described by Ghosh *et al*.^44^ to generate *hqt*-silenced and control eggplant lines.

### Nutritional indices

Nutritional indices explain food consumption, utilization and excretion budgets, and the state of digestion physiology, which helps estimate the diet’s suitability. Nutritional indices of the armyworm larvae feeding on eggplant (control, *hqt*-silenced, and CGA-complemented) leaves and artificial diets (spiked with different candidate metabolites) were calculated as described by Waldbauer^45^ and as standardized for armyworm larvae by Mitra *et al*.^34^ and Umesh *et al*.^12^. Ingestion and excretion were budgeted using the excretion efficiency determination assays^12^. Larvae freshly molted into the fourth instar were first starved for 4 h and then fed on a measured amount of artificial diet spiked with CGA (650 µg/g). After 24 hours of feeding, larvae were starved again for 4 hours. The mass of larvae and frass was measured before and after every starvation. Frass samples were collected and stored at -80°C for UPLC/ESI-QTOF-MS analysis.

### Dual-choice and no-choice assays to analyze natural enemies’ survivorship and performance

All the predators (spiders, ants, and mantid) used in the choice assays were starved for 48 h before the experiment. For the dual-choice assays conducted to analyze predators’ preferences, CGA-ingested and AD-fed larvae were provided to each predator individual as choices. For the no-choice assays performed to analyze predators’ performances and survivorships, CGA-ingested and AD-fed larvae were provided separately to each predator individual. In the preference determination assays (1 h duration), complete ingestion of the prey (armyworm larva) by the predator was scored as ‘1’, and no ingestion and prey abandoning were scored as ‘0’ (n= 3, each with 5 predator individuals). In the performance analysis assays, the number of larvae ingested by each predator individual in 24 h was recorded (n= 3, each with 5 predator individuals). Predator mortality was recorded in the survivorship assays (duration= 48 h; n= 3, each with 5 predator individuals).

### Statistical analysis

Field experiment data from the randomized blocks were analyzed both collectively and separately. Correlation analysis was performed to understand whether armyworms’ differential larval occurrence and performance are influenced by chemical composition and their differential abundance among different eggplant varieties. Directions and strengths of correlations of armyworm’s larval occurrence with larval mass and larval mortality in larval performance assays were calculated using Spearman’s Rho (*r*_*s*_) tests. Two-tailed student’s test (*P*≤ 0.05) was used to determine the significance of obtained correlation results. Principal component analysis (PCA) was performed based on Bray-Curtis dissimilarities using Past 3.26 to determine the eggplant type with key chemical components contributing to how they are differentially preferred by armyworm larvae when given a choice in the field^46^. The homogeneity of the quantitative data was tested using Levene’s, and the normality was tested using the Jarque-Bera test. The homogenous and normal data were analyzed using one-way ANOVA, and the statistical significance was determined using Tukey’s *post hoc* test (*P*≤ 0.05). No-choice assay data were analyzed using a student’s t-test (one-tailed, *P*≤ 0.05).

## Results

### Armyworm’s eggplant variety preferences

The armyworm larvae showed differential occurrence and feeding on the seven eggplant varieties; the highest occurrence was observed on VJ (13.7± 2.98) and CVK (12.8± 3.33), followed by JG (7.1± 3.33), KV (6.1± 3.33), HK (4.0± 1.32), and KP (2.7± 1.18) (Figure 1B). RL22 had the lowest (near zero) larval occurrence. Selected eggplants varieties were: Himalayan eggplant variety (RL22), Ankur Kavach (KV), JK Agro Hybrid-green long (JK), Omaxe-CVK MK (CVK), Ankur Vijay (VJ), L-Riccia Hirvi Kateri (HK), and KGN’s F1 Pinstripe (KP). Larval performance (measured in terms of larval mass) showed a similar trend. Larval mass was the highest on VJ (505± 20), followed by CVK (456± 20), JG (306± 18), KV (288± 18.2), HK (232± 16.4), and KP (217± 13.7) (Figure 1C). Larvae reared on RL22 showed the lowest mass (200± 10.7). Similarly, the neonate mortality was the highest on RL22 (92.22± 2.94), followed by HK (85.55± 2.23) and KP (82.23± 4.00), whereas lowest on KV (54.45± 4.01), JG (53.34± 5.10), CVK (42.23± 2.94), and VJ (34.45± 2.94) (Figure 1D). Larval mortality in the later instar stages also showed a similar pattern (Figure 1D).

**Figure 1.**
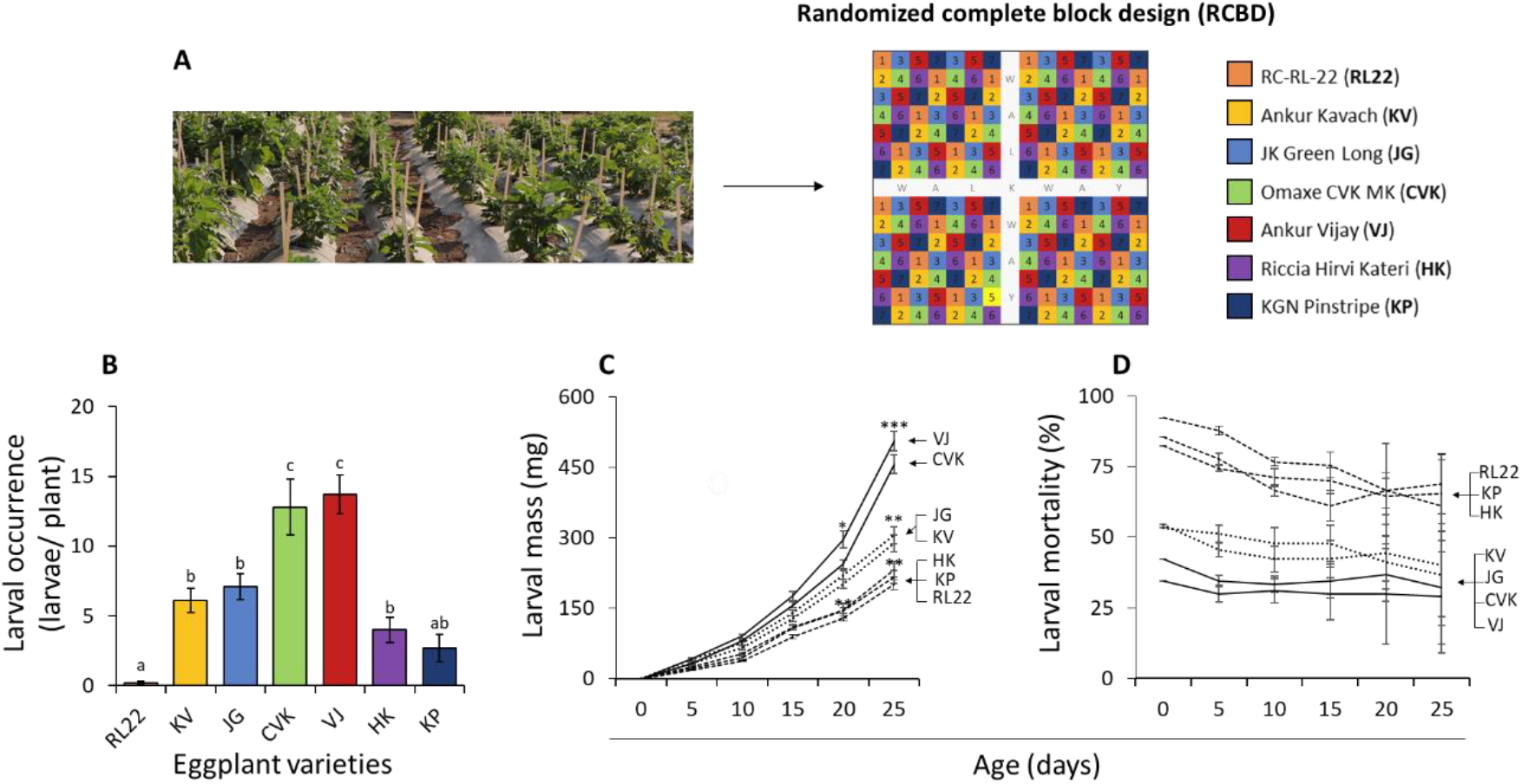
Armyworm larval occurrence and performance. (A) A schematic of the experimental field’s randomized complete block design (RCBD) with seven eggplant varieties (RL22, KV, JG, CVK, VJ, HK, and KP). (B) Natural occurrence of the armyworm larvae on seven eggplant varieties (one-way ANOVA, F_6, 63_= 19.1, *P*≤ 0.0001, n= 10 plants). (C) Mass of armyworm larvae feeding on the leaves of the seven eggplants varieties [One-way ANOVA: 5^th^ day (F_6, 161_= 23.45, *P*≤ 0.0001), 10^th^ day (F_6, 161_= 86.45, *P*≤ 0.0001), 15^th^ day (F_6, 161_= 72.98, *P*≤ 0.0001), 20^th^ day (F_6, 161_= 205.67, *P*≤ 0.0001), 25^th^ day (F_6, 161_= 261.8, *P*≤ 0.0001)). (D) Larval mortality on eggplant varieties [one-way ANOVA, neonates (F_4, 53_= 18.54, *P*≤ 0.0001), 5^th^ day (F_4, 53_= 78.25, *P*≤ 0.0001), 10^th^ day (F_4, 53_= 95.8, *P*≤ 0.0001), 15^th^ day (F_4, 53_= 46.2, *P*≤ 0.0001), 20^th^ day (F_4, 53_= 27.45, *P*≤ 0.0001), 25^th^ day (F_4, 53_= 15.21, *P*≤ 0.0001)]. Significant differences (*P*≤ 0.05) were determined using Tukey’s *post hoc* test.

### Candidate compound discovery

To test whether the armyworm’s differential occurrence and performance were associated with the eggplant secondary metabolite(s), we analyzed the eggplant leaf metabolome using a UPLC-ESI-QTOF-based non-targeted metabolomics analysis. A total of 332 metabolites were identified (Table S1 & S2). Metabolites identified using MSDIAL software were used to perform multivariate analyses to group the eggplant varieties based on their chemical compositions; the Principal Component Analysis (PCA) showed the separate clustering of RL22 from the other varieties (Figure 2A). CGA contributed the most (0.89 loading on PC1) to this separation.

**Figure 2.**
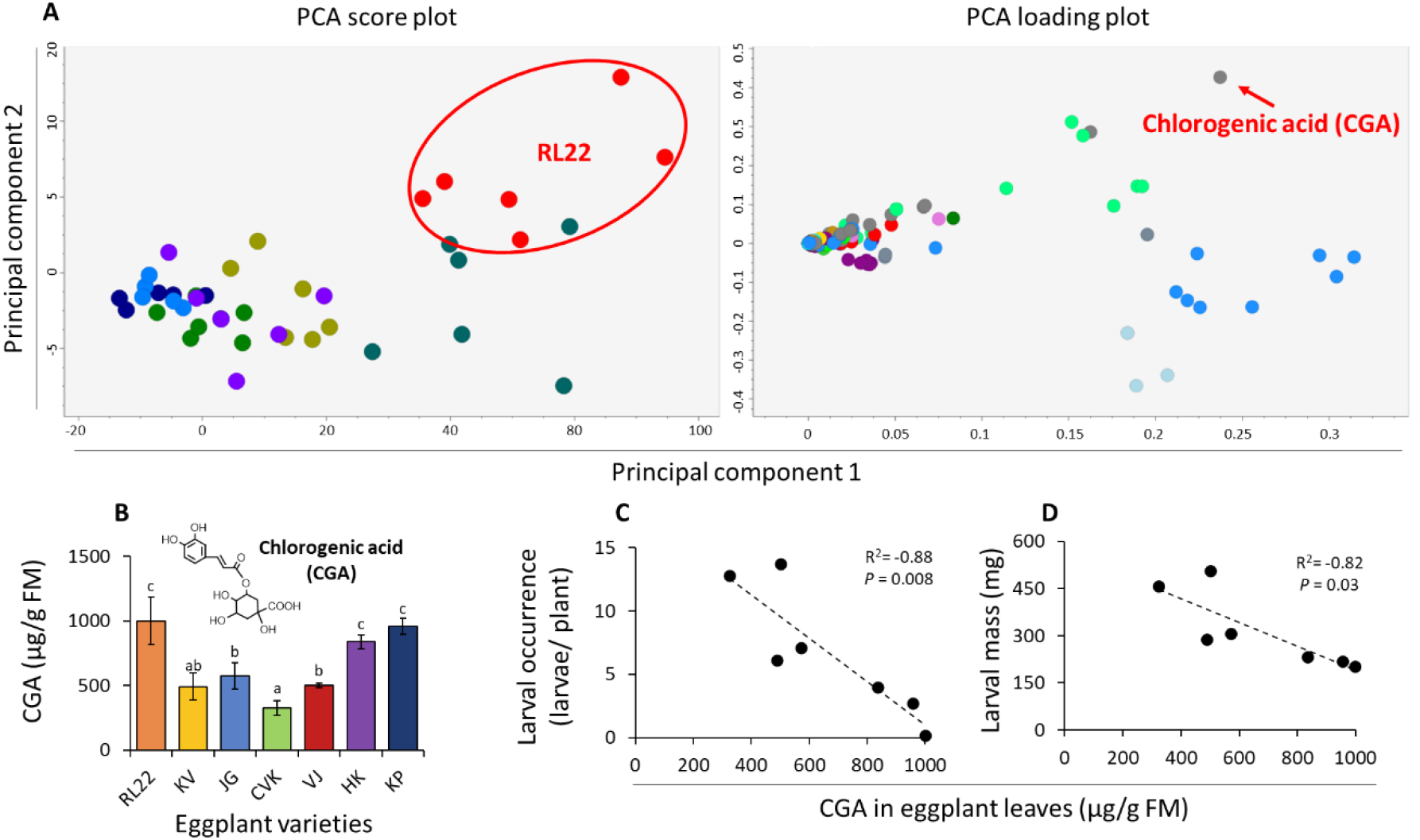
Eggplant CGA content negatively correlates with the armyworm larval occurrence and performance. (A) Principal Component Analysis (PCA) shows the separately clustered RL22 based on its CGA contribution (0.89 loading on PCA1). (B) Foliar CGA content of the seven eggplant varieties (one-way ANOVA, F_6, 35_= 8.91, *P*≤ 0.0001). Spearman’s (*r*_s_) correlation between eggplant varieties’ CGA concentrations and (C) larval occurrence, and (D) larval mass. For one-way ANOVA, statistical significance (*P*≤ 0.05) was determined using Tukey’s *post hoc* test. For Spearman’s Rho (rs), the student’s two-tailed t-test (*P*≤ 0.05) was performed to determine the correlation significance.

RL22 showed the highest CGA concentration (999.57± 185.07), followed by KP (957.26± 106.96) and HK (836.86± 130.67), whereas lowest in JG (572.85± 54.49), VJ (502± 62.37), KV (489.58± 104.24), and CVK (324.74± 56.05) (Figure 2B). Eggplant varieties’ CGA contents showed a strong negative concentration with larval occurrence (*R*^*2*^= -0.88, *P*= 0.008) (Figure 2C) and larval mass (*R*^*2*^= -0.82, *P*= 0.03) (Figure 2D).

### CGA’s effect on the armyworm

To ascertain that CGA was associated with the larvae’s differential occurrence and performance, larvae were fed CGA via an artificial diet (AD). The cumulative larval and pupal duration of the CGA-fed larvae was 1.5-fold more than the control AD-fed larvae (Figure 3B, C). Similar effects of CGA ingestion were seen on the adult life span, egg number, and egg hatching, as they showed a 2.04-fold increase, 2.75-fold decrease, and 1.48-fold decrease, respectively than the AD-fed controls (Figure 3D-F). Likewise, the mass of the CGA-ingested larvae was 2-fold lower than the controls. CGA ingestion also increased larval mortality by >3-fold compared to the controls (Figure G, H).

**Figure 3.**
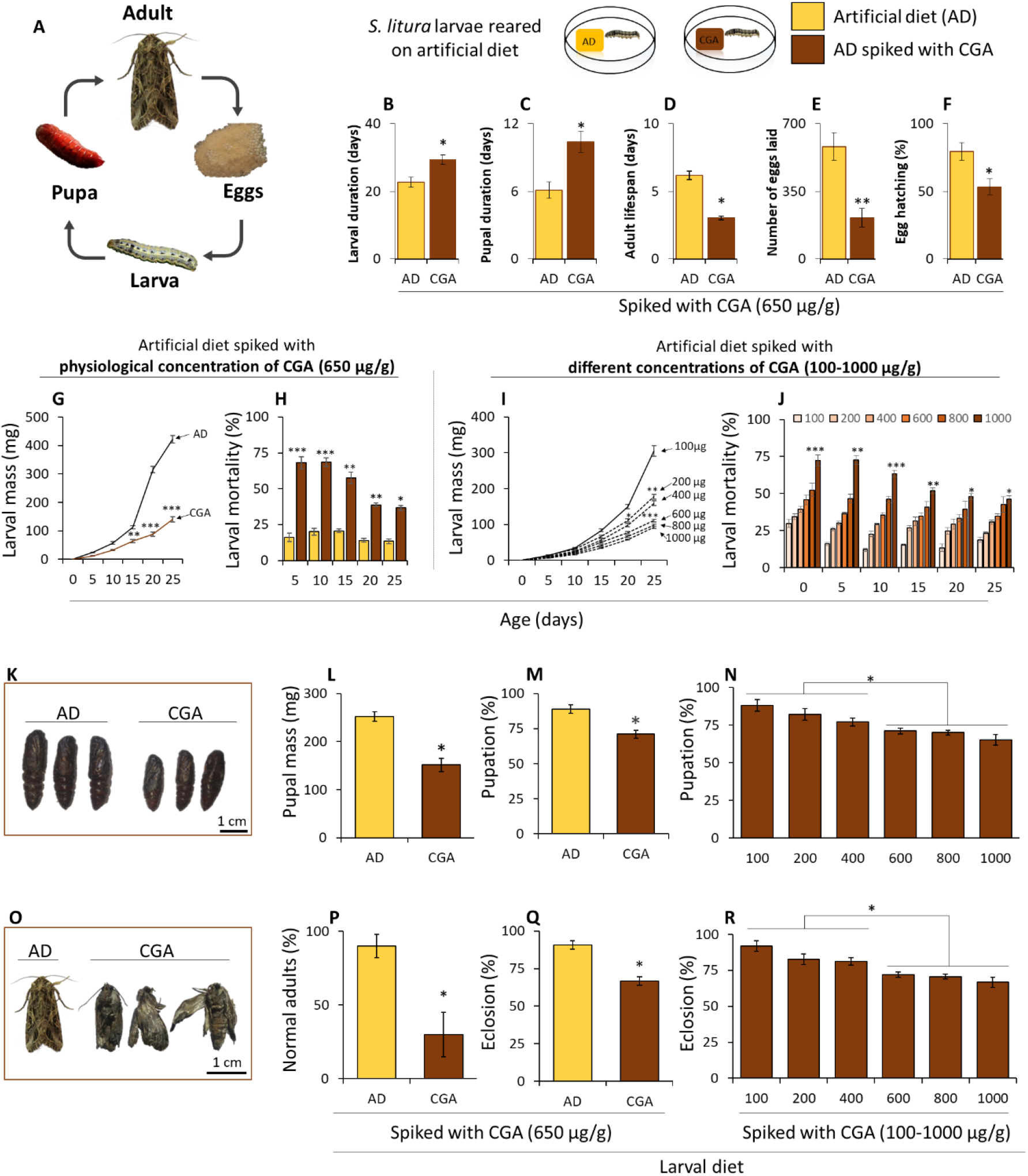
Eggplant’s chlorogenic acid negatively affects armyworm’s performance when fed via an artificial diet. (A) Armyworm life cycle. (B) Larval duration, (C) pupal duration, (D) adult lifespan, (E) egg laying, and (F) egg hatching when larvae were reared on the CGA-spiked artificial diet (student’s two-tailed t-test, *\*≡P<* 0.05, \**\*≡P<* 0.001, n= 30 larvae). (G) Larval mass and (H) mortality when reared on the CGA-spiked artificial diet (student’s two-tailed t-test, *\*≡P<* 0.05, \**\*≡P<* 0.001, \*\**\*≡P<* 0.0001, n= 30 larvae). (I) Larval mass [one-way ANOVA, neonates (all masses were considered “ 0”), 5^th^ day (F_5, 174_= 37.27, *P*≤ 0.0001), 10^th^ day (F_5, 11_= 18.32, *P*≤ 0.0001), 15^th^ day (F_5, 11_= 40.16, *P*≤ 0.0001), 20^th^ day (F_5, 11_= 78.63, *P*≤ 0.0001), 25^th^ day (F_5, 11_= 115.3, *P*≤ 0.0001)] and (J) larval mortality [one-way ANOVA, neonates (F_5, 12_= 12.22, *P*≤ 0.0001), 5^th^ day (F_5, 11_= 95.8, *P*≤ 0.0001), 10^th^ day (F_5, 11_= 95.8, *P*≤ 0.0001), 15^th^ day (F_5, 11_= 26.2, *P*≤ 0.0001), 20^th^ day (F_5, 11_= 13.56, *P*≤ 0.0001), 25^th^ day (F_5, 11_= 15.21, *P*≤ 0.0001)] when larvae were fed on different CGA concentration-spiked diet. (K) Pupae of the larvae reared on CGA-spiked and control diets. (L) pupal mass and pupation (%), when the larvae were reared on CGA-spiked (physiological concentration) and control diets (student’s two-tailed t-test, *\*≡P<* 0.05, \**\*≡P<* 0.001, \*\**\*≡P<* 0.0001, n= 30 larvae). (N) pupation (%), when larvae were reared on a diet spiked with different CGA concentrations (one-way ANOVA, F_5, 24_= 8.18, *P*= 0.0001). (O) Normal and deformed adults of the larvae were reared on control and CGA-spiked diets, respectively. (P) healthy adults (%) and (Q) eclosion (%), when armyworm larvae were reared on a CGA-spiked diet) (student’s two-tailed t-test, *\*≡P<* 0.05, \**\*≡P<* 0.001, \*\**\*≡P<* 0.0001, n= 30 adults). (R) eclosion (%), when larvae were reared on diets spiked with different CGA concentrations (one-way ANOVA, F_5, 24_= 6.92, *P*= 0.0004). For one-way ANOVA, statistical significance (*P*≤ 0.05) was determined using Tukey’s *post hoc* test.

We also ascertained the CGA effect by analyzing the larvae feeding on the diets of different CGA concentrations. We observed that the larval mass decreased and larval mortality increased with the CGA concentration increase (Figure 3I, J). CGA ingestion caused a 1.6-fold and 1.25-fold decrease in the pupal mass (Figure 3K, L) and pupation rate (Figure 3M), respectively, to the controls. Pupation (%) was also reduced with the diet’s increasing CGA concentrations (Figure 3N). Adults of the CGA-fed larvae showed deformities (Figure 3O); the number of normal adults was 2.4-fold lower in the CGA-fed treatments than the controls (Figure 3P). CGA-fed insects’ eclosion (%) was also 1.36-fold lower than the controls (Figure 3Q). It showed a consistent decline with the diet’s increasing CGA concentration (Figure 3R).

### Reverse genetics approach to assess the insecticidal potential of CGA *in planta*

We used a reverse genetics analysis to ascertain whether the armyworm’s low occurrence and performance on RL22 were associated with CGA. CGA-deplete RL22 plants were generated by silencing the CGA biosynthesis gene HQT. First, the *Sm*HQT gene sequence was mined from the eggplant genome database using a sequence similarity analysis (Figure 4A). Only one HQT was found in the eggplant. *Sm*HCT was the closest eggplant gene to *Sm*HQT (54.37% sequence similarity). *Sm*HQT showed >95% sequence similarity with the reported HQT gene from the other *Solanum* spp. and was placed in the same clade with potato, tomato, and tobacco HQTs. *Sm*HQT was silenced using the VIGS (Figure 4B). *Sm*HQT transcript levels were 5.65-fold lower in *Sm*HQT-silenced plants than in the wild type (WT) and empty vector-infiltrated (EV) control plants (Figure 4C). An off-target effect was observed on the *Sm*HCT transcript levels (Figure 4D). *Sm*PDS silencing in eggplant was used as a control to standardize the VIGS procedure and as a visual indicator of VIGS spread and gene silencing effect.

**Figure 4.**
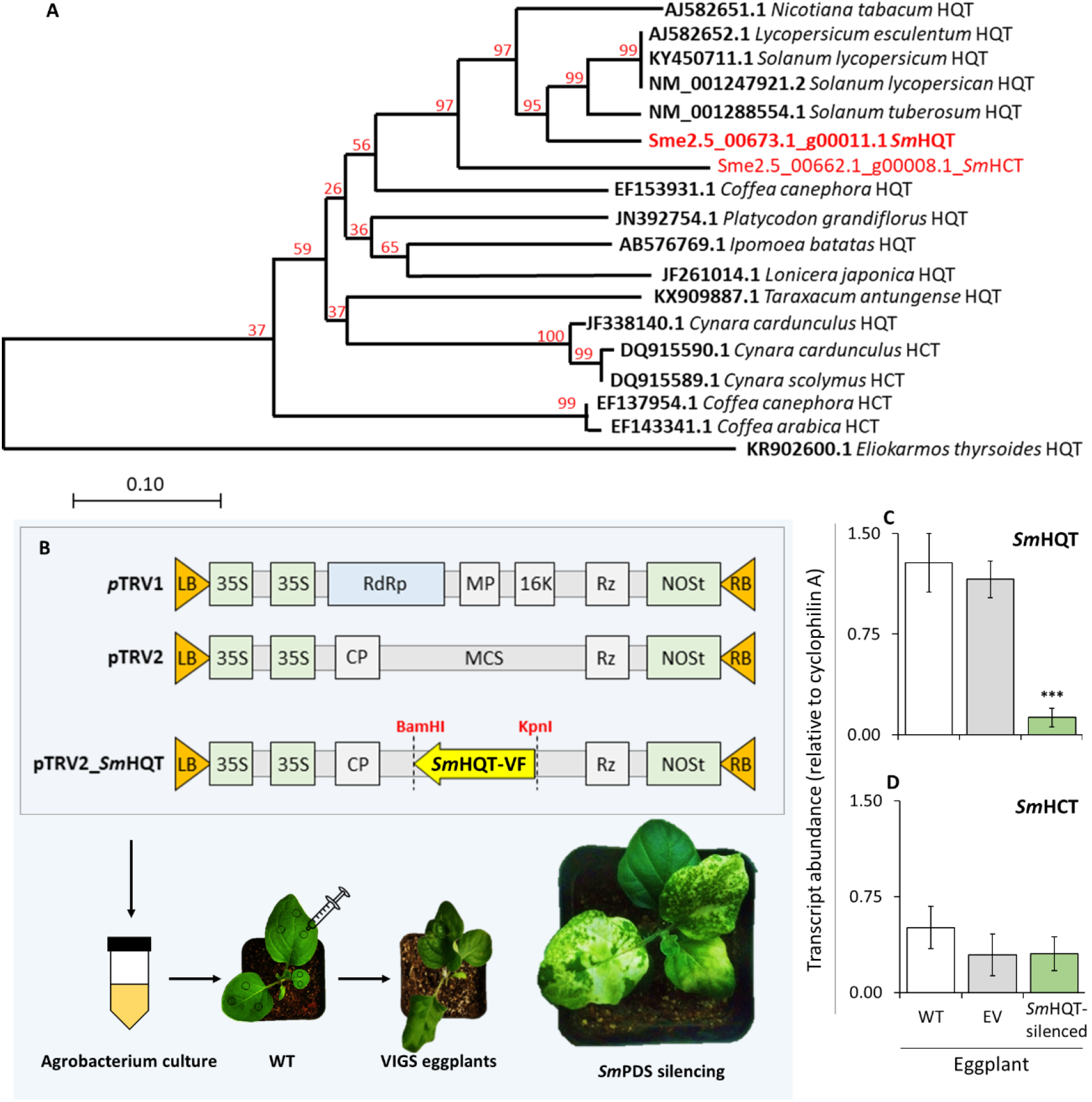
*Sm*HQT gene silencing causes reduced CGA biosynthesis in eggplant. (A) Phylogenetic tree constructed using the neighbor-joining (Dayhoff matrix) algorithm. Bootstrap values are given for each branch (1000 replicates). The tree was built using all the reported *Sm*HQT genes (from NCBI) of Solanaceae and other plant families. Candidate *Sm*HQT and *Sm*HCT genes are highlighted. **(B)** A schematic of *p*TRV vector constructs used for the *Sm*HQT VIGS. *Sm*PDS-silencing was used as a VIGS positive control. (C) *Sm*HQT transcript abundance (relative to Cyclophilin A) in the leaves of wild type (WT), empty vector-infiltrated (EV), and *p*TRV2*-Sm*HQT construct-infiltrated (*Sm*HQT-silencing) eggplants (student’s two-tailed t-test, *\*≡P<* 0.05, \**\*≡P<* 0.001, \*\**\*≡P<* 0.0001, n= 10 plants). (D) *Sm*HCT transcript abundance (relative to Cyclophilin A) in the leaves of wild type (WT), empty vector-infiltrated (EV), and *p*TRV2*-Sm*HQT construct-infiltrated (*Sm*HQT-silencing) eggplants (student’s two-tailed t-test, *P*> 0.05; n= 10).

A phylogenetic tree constructed using all the reported HQT genes from the Solanaceae plant family suggested that *Sm*HQT-silenced plants showed 4.9-fold lower CGA concentrations than the WT and EV controls (Figure S1).

### Effect of *Sm*HQT silencing on eggplant’s other phenolics

Whether the *Sm*HQT silencing affected the other phenolics’ whose biosynthesis is associated with the CGA biosynthesis pathway was analyzed. These phenolics did not show concentration differences compared to the controls (Figure S1).

### Effect of SmHQT silencing on the larval performance

To analyze the effect of *Sm*HQT-silencing on the armyworm larvae, untreated (control), water-complemented (control), and CGA-complemented WT, EV, and *Sm*HQT-silenced plants were used (Figure 4). Further, the larval performance on these treatments was analyzed with the help of nutritional indices (Figure 5). Consumption index (CI), approximate digestibility (AD), and the relative growth rate (GR) of the larvae increased in *Sm*HQT-silenced than the controls (Figure 5D-F). CGA complementation reduced these indices. The efficiency of conversion of both ingested (ECI) and digested food (ECD) also increased in the larvae feeding on the *Sm*HQT-silenced plants as compared to the controls (Figure 5G, H).

**Figure 5.**
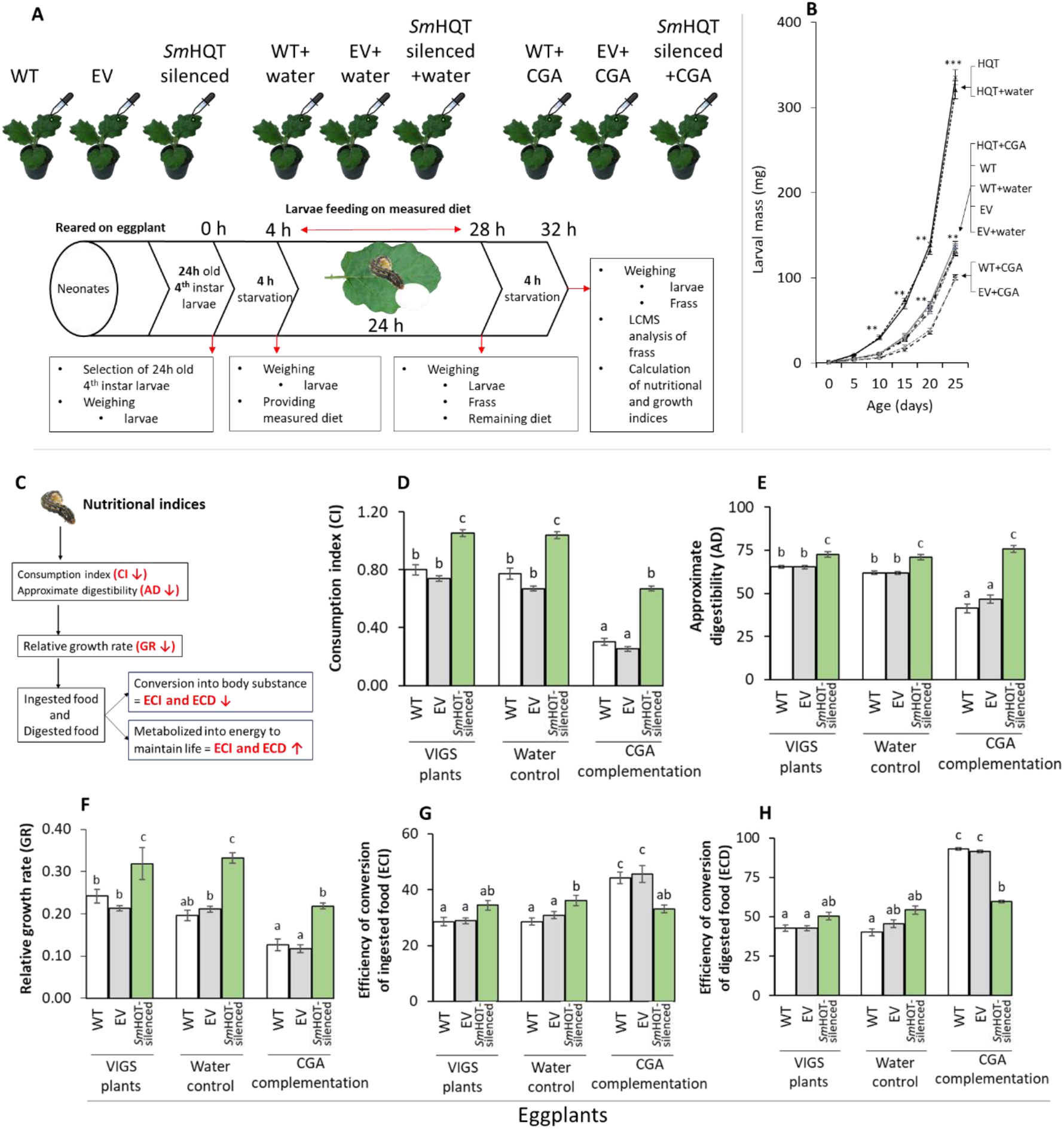
Nutritional indices study shows an adverse effect of CGA on armyworm larvae and negatively affects its performance. (A) A schematic of the ingestion-excretion budgeting assays and the eggplant lines used in the experiment. (B) Larval mass on the different lines [one-way ANOVA, 5^th^ day (F_8, 261_= 32.82, *P*≤ 0.0001), 10^th^ day (F_8, 261_= 95.4, *P*≤ 0.0001), 15^th^ day (F_8, 261_= 74.43, *P*≤ 0.0001), 20^th^ day (F_8, 261_= 215.4, *P*≤ 0.0001), 25^th^ day (F_8, 261_= 281.7, *P*≤ 0.0001)). (C) Different nutritional indices and their relations. (D) Consumption index [(CI), F_8, 216_= 122.6, *P*≤ 0.0001], (E) approximate digestibility [(AD), F_8, 216_= 48.64, *P*≤ 0.0001], (F) relative growth rate [(GR), F_8, 216_= 42.11, *P*≤ 0.0001], (G) efficiency of conversion of ingested food [(ECI), F_8, 216_= 119.5, *P*≤ 0.0001], (H) efficiency of conversion of digested food [(ECD), F_8, 216_= 13.47, *P*≤ 0.0001] of the larvae fed on WT, EV, and *Sm*HQT-silenced eggplants, with water-control and CGA complementation. Statistical significance (*P*≤ 0.05) was determined using Tukey’s *post hoc* test.

Mass (mean± SE) of the larvae feeding *Sm*HQT-silenced (335.46± 8.77) and *Sm*HQT-silenced+water (320.56± 10.80) eggplant lines was the highest (Figure 5B). The mass of the larvae feeding WT+CGA (100.76± 2.71) and EV+CGA (98.73± 2.36) plants was the lowest. Larvae feeding on the WT, EV, WT+water, and EV+water treatments gained similar mass. CGA complementation to the *Sm*HQT-silenced eggplants restored the WT phenotype (Figure 5B).

## Discussion

Plant extracts and pure phytochemicals have been used as biopesticides for a long time^47^. Since phytochemicals are natural compounds, they are considered to be safer for farm workers, agricultural produce consumers, and also for the environment, than the synthetic pesticides; therefore, they are often preferred over the synthetic pesticides^48,49^. Especially the organic farming, in which the synthetic compounds are not used, heavily relies on such botanicals^49,50^. Pests’ non-host or less preferred host species are most likely to contain deterrent, antifeedant, or antidigestive plant metabolites; therefore, such species are often used to obtain the biopesticides^51-56^. Several factors underlie the discovery and establishment of such compounds as insecticides. Fundamental and the most important factor is that such compounds are often present in plants as constituents of complex mixtures of chemically similar compounds, which render their identification, purification, characterization, and commercialization challenging^50,53^. In this work, which originated from a vital field observation on the ecology of the multi-insecticide resistant polyphagous pest armyworm, we integrated the pest’s behavioral ecology, metabolomics, and reverse-genetics approaches to identify the CGA as a potential biopesticide. We found that CGA affects larval physiology, growth, and development that is associated with armyworm’s high mortality and low occurrence on high CGA-containing plants. Armyworm larvae need to use a two-step detoxification process to counter-adapt CGA, which incurs physiological costs. Lastly, we found that when used as a pesticide against the armyworm, CGA does not harm the armyworm’s natural enemies. CGA is a dietary supplement and an antioxidant for humans. Thus, it is safe for human consumption. Together, high CGA-containing varieties can be used to reduce the armyworm infestation risk, and CGA can be used as a biopesticide.

Eggplant is one of the highest insecticide-applied vegetables since all of its developmental stages are susceptible to attacks by various insect pests^57-63^. Armyworm, mainly a folivore, cause severe damage to the early vegetative stages of the eggplant crop. Its differential occurrence on seven co-growing eggplant varieties hinted at the varying suitability of these varieties to it. This hypothesis was further strengthened by the larvae’s differential larval performance on these varieties. CGA is a defense metabolite found across many plant groups and is the most abundant phenolic in eggplant^26,64,65^. In some insect species, CGA interacts with the dietary proteins and interferes with their food digestion and absorption in the gut ^24,26^. It is likely that due to this activity of CGA, an adverse effect was observed on the armyworm’s larval mass when fed on an eggplant leaf. This result is congruent to that of Stevenson *et al*., who showed CGA’s inhibitory effect on armyworm larval development^33^; authors found CGA as an important factor in groundnut’s (*Arachis paraguariensis* hodat & Hassl.) resistance to the armyworm. Duffey and Isman also showed CGA’s negative effects on corn earworm (*Heliothis zea* Boddie)^66^. Leiss *et al*. also identified CGA as a resistance factor of chrysanthemum [*Dendranthema grandiflora* (Ramat.) Kitam.] against the thrips by comparing the metabolomic profile of thrips-resistant (high CGA) and thrips-susceptible (low CGA) plants^67^; they validated CGA’s negative effects using the AD-based bioassays^67^. Kundu and Vadassery reviewed that CGA has been proven to be an efficient defense molecule against a broad range of insect herbivores^26^. Although CGA has been reported as an efficient larvicide, its effect on an insect’s overall life cycle is unexplored. This study provides insights into the ill effects of CGA on other developmental stages like pupa and adult.

RNAi-based reverse genetics methods have caused a major impact on plant secondary metabolites research^68,69^. For example, Bi *et al*. could reveal the effects of phenolics, including CGA, on tobacco hornworm (*Manduca sexta* Linnaeus) and tobacco budworm (*Heliothis virescensi* Fabricius)^70^; they found that the phenolic content reduction in the tobacco plants rendered them susceptible to these herbivores. We also found that upon silencing *Sm*HQT, eggplant was rendered susceptible to the armyworm. Larval performance improved on the *Sm*HQT-silenced plants, and the CGA complementation could restore the resistance on these plants. These results indicated that CGA is eggplant RL22’s resistance factor. This conclusion is corroborated by finding from a study by Bejai *et al*. (2012) when the Egyptian cotton leafworm (*Spodoptera littoralis* Boisduval) showed lower mass gain with an increase in glucosinolate concentration when they were fed on WT and glucosinolate-overexpressed Arabidopsis^71^.

Analysis of nutritional and growth indices facilitated the understanding of CGA’s effect on larval performance. When the fourth instar armyworm larvae were fed on eggplant leaves, the growth rate (GR) was significantly reduced with increased concentration in CGA-complemented WT, EV, and *Sm*HQT-silenced eggplant compared to controls. This corresponds to a similar reduction in consumption index (CI) and approximate digestibility (AD). This decrease in CI is likely due to the extract’s antifeedant nature, which accounts for the decrease in GR. Similar antifeedant effects were observed by Wheeler and Isman; when they increased the diet concentrations of cape mahogany (*Trichilia americana* Sessé & Mociño) plant extracts, they found reductions in ECI, ECD, GR, and CI of armyworm larva ^72^.

Another study by Xie *et al*. (1994) showed a similar effect of hirtin and Indian heynea (*Trichilia connaroides* Wight & Arn.) extract when ingested by pearly underwing moth (*Peridroma saucia* Hübner) and armyworm^73^. Neem (*Azadirachta indica* A. Juss) seed kernel extract and Chinese chaste tree (*Vitex negundo* L.) leaf extract also showed similar effects on the nutritional indices of the rice leafroller (*Cnaphalocrocis medinalis* Guenée)^74^. ECI is an overall measure of an insect’s ability to utilize the food that it ingests for growth. A drop in ECI indicates that more food is being metabolized for energy (used for detoxification) and less is being converted to the body substance (i.e., growth). In our study also, both ECI and ECD decreased upon CGA ingestion. Since WT and EV already had high concentrations and CGA complementation increased the toxin concentration in the diet, which could be lethal to larva, that’s why armyworm had to switch the conversion of ingested and digested food from body substance to energy production for detoxification-related metabolism. Change in these indices depending on the CGA concentration in the diet of armyworm supports the potential of CGA as a candidate biopesticide.

To summarize, this study showed that CGA exhibits larvicidal properties against the armyworm. It is also safe for beneficial organisms. Thus, CGA is a promising biopesticide candidate for the field trial phase against Lepidopteran pests, especially armyworm. CGA alone or in combination with other insecticides to target the pests could be integrated into future pest control measures in integrated pest management (IPM).

## Supporting information

Kumar et al 2023_SI.pdf

## Acknowledgments

Authors thank Ministry of Human Resource Development, India, for financial support and funding the AB Sciex X500R UPLC-ESI-QTOF system. MK thanks the Council of Scientific and Industrial Research India, for Ph.D. fellowship and U.K.P thanks Department of Science and Technology, India for Inspire fellowship. authors thank IISER Pune for providing the on-campus field site, Mr. G. Pingalkar for field support, and Mr. G. Pawar for help in field experiments.

## Author Contributions

MK and SP conceived and designed the experiments. All authors performed the experiments and collected the data. MK conducted the statistical analyses. All authors interpreted and discussed the results. MK and SP wrote the manuscript with inputs from all the authors; SP acquired funds, administered the project, and supervised the research.

## The following supporting information is available for this article

**Figure S1** | SmHQT gene silencing in eggplant leaf did not affect the flux of other phenolics’ biosynthesis and concentrations

**Table S1** | Primers used in the cloning and transcript quantitation experiments

**Table S2** | Metabolites identified from the seven different eggplant varieties using the non-targeted metabolomics.

**Table S3** | List of identified and annotated metabolites in eggplant leaves and their correlations with larval occurrence, mass, and mortality.

