## Supplementary material for "Eggplant’s foliar chlorogenic acid provides resistance against the tropical armyworm": Kumar et al 2023_SI.pdf

**The following supporting information is available for this article:**

**Figure S1 |** *Sm*HQT gene silencing in eggplant leaf did not affect the flux of other phenolics' biosynthesis and concentrations

**Table S1 |** Primers used in the cloning and transcript quantitation experiments

**Table S2 |** Metabolites identified from the seven different eggplant varieties using the non-targeted metabolomics.

**Table S3 |** List of identified and annotated metabolites in eggplant leaves and their correlations with larval occurrence, mass, and mortality.

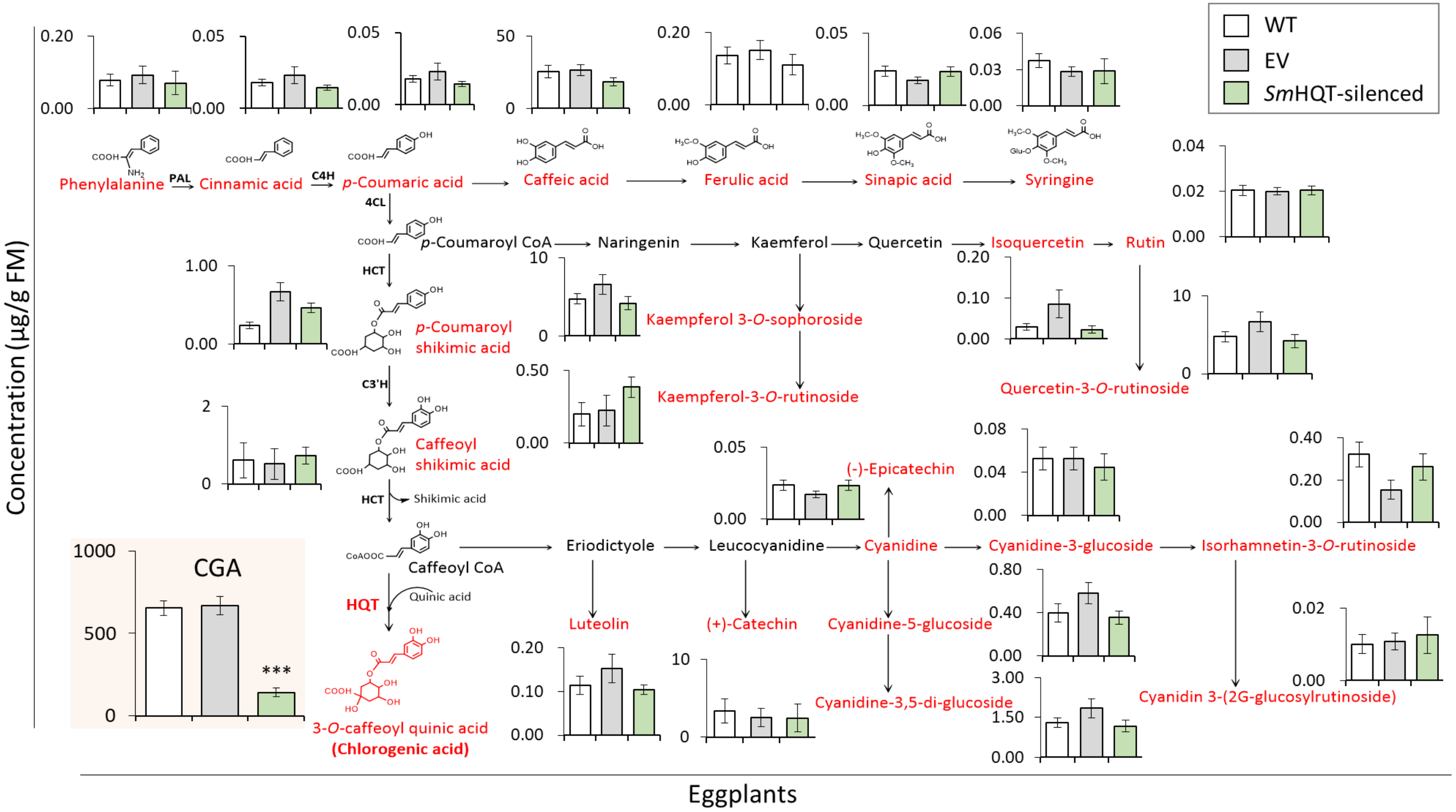


**Figure S1 | *Sm*HQT gene silencing in eggplant leaf did not affect the flux of other phenolics' biosynthesis and concentrations.** Concentrations of all the phenolics in the control and SmHQT-silenced eggplant leaf. Statistical significance was determined by the student's two-tailed t-test ($P \leq 0.05$, n=10).

**Table S1 |** Primers used in the cloning and transcript quantitation experiments.

| **Sequence ID** | **Sequence name** | **Gene code** | **Forward and reverse primers (5'-3')** |
|---|---|---|---|
| **Primers for VIGS fragment amplification and cloning** | | | |
| Sme2.5_00673.1_g00011.1 | *Solanum melongena* hydroxycinnamoyl coenzyme A-quinate transferase mRNA | *Sm*HQT | CCTTCTTTTGAGCATGTCGAG |
| | | | GAGCTTCAATCAGAACCGTTG |
| **RT-qPCR primers of *Sm*HQT, *Sm*HCT** | | | |
| Sme2.5_00673.1_g00011.1 | *Solanum melongena* hydroxycinnamoyl coenzyme A-quinate transferase mRNA, complete cds | *Sm*HQT | GATATCTCAACCTTCCCACTCG |
| | | | CAGATAACGTGTGGAACACTCC |
| Sme2.5_04555.1_g00001.1 | *Solanum melongena* hydroxycinnamoyl-CoA shikimate/quinate hydroxycinnamoyl transferase mRNA | *Sm*HCT | ATTGTAAGGGGCAAGGTGTG |
| | | | TAATCAACGGCGGGAATAAG |
| **RT-qPCR primers for the eggplant reference gene** | | | |
| Sme2.5_00563.1_g00006.1 | *Solanum melongena* Cyclophilin A (CyP), mRNA | Cyclo a | CAAAACCGCTGAGAACTTCCG |
| | | | CTTGACACATGAACCCTGGGA |

**Table S2 |** Metabolites identified from the seven different eggplant varieties using the non-targeted metabolomics.

| Class | Compound Name | Molecular formula | Concentrations [relative to formononetin (IS)] among different eggplant varieties [ng $g^{-1}$ FM (mean± SE)] | | | | | | |
|---|---|---|---|---|---|---|---|---|---|
| | | | 0 | KV | JG | CVK | VJ | HK | KP |
| Aldehyde and ketones | 1-(4-methoxyphenyl) ethanone | C9H10O2 | 32.38± 10.07 | 36.04± 11.33 | 19.36± 7.92 | 10± 1.88 | 7.86± 2.01 | 26.85± 4.89 | 22.46± 7.72 |
| | Dihydrojasmone | C11H18O | 49.75± 7.11 | 37.02± 4.82 | 34.23± 10.91 | 19.8± 3.49 | 29.79± 2.48 | 25.87± 2.41 | 22± 3.57 |
| | Phenylacetaldehyde | C8H8O | 369.9± 27.1 | 565.35± 51.72 | 254.84± 38.51 | 218.38± 21.64 | 262.96± 11.61 | 286.79± 46.89 | 302.45± 37.21 |
| | 3-Formylindole | C9H7NO | 14.76± 4.55 | 22.02± 9.62 | 7.89± 1.86 | 10.12± 2.9 | 4.56± 1.65 | 12.45± 2.01 | 8.52± 4.13 |
| | 4-Hydroxybenzaldehyde | C7H6O2 | 40.72± 8.28 | 60.76± 20.56 | 21.6± 5.54 | 24.16± 4.62 | 8.67± 3.46 | 31.61± 7.58 | 13.39± 4.03 |
| | Protocatechuic aldehyde | C7H6O3 | 311.34± 74.63 | 194.57± 38.33 | 95.49± 14.24 | 102.49± 16.14 | 84.47± 11.58 | 82.44± 14.45 | 53.91± 22.93 |
| Alkaloids and derivatives | Brucine | C23H26N2O4 | 122.95± 10.5 | 9.65± 0.51 | 23.12± 6.64 | 11.06± 1.98 | 14.27± 1.73 | 19.18± 5.99 | 10.38± 3.06 |
| | Calycanthine | C22H26N4 | 973.55± 223.01 | 1064.99± 303.62 | 767.27± 165.29 | 1122.97± 486.76 | 689.27± 225.21 | 1098.26± 428.83 | 766.9± 374.88 |
| | Cinchonine | C19H22N2O | 11.69± 1.98 | 2.6± 0.27 | 2.46± 0.65 | 1.84± 0.3 | 0.9± 0.16 | 1.75± 0.64 | 0.99± 0.1 |
| | Hirsuteine | C22H26N2O3 | 2.19± 0.95 | 3.85± 0.4 | 1.51± 0.2 | 1.39± 0.43 | 0.88± 0.17 | 1.03± 0.27 | 0.94± 0.23 |
| | HYDROQUINIDINE | C20H26N2O2 | 110.83± 41.02 | 17.84± 2.37 | 12.12± 2.24 | 8.61± 3.02 | 3.19± 0.99 | 8.31± 2.37 | 9.14± 2.68 |

| | | | | | | | | | |
|---|---|---|---|---|---|---|---|---|---|
| | Imperialine | C27H43NO3 | 11.48± 6.04 | 2.13± 0.42 | 4.44± 1.51 | 5.32± 1.87 | 2.9± 1.94 | 3.66± 1.7 | 0.82± 0.31 |
| | Khasianine | C39H63NO11 | 6.93± 6.88 | 0.46± 0.16 | 0.8± 0.33 | 0.16± 0.08 | 0.21± 0.05 | 0.88± 0.29 | 0.18± 0.04 |
| | quinidine | C20H24N2O2 | 2.23± 0.42 | 2.24± 0.31 | 1.73± 0.28 | 1.29± 0.4 | 1.75± 0.3 | 1.53± 0.23 | 1.02± 0.48 |
| | Ranaconitine | C32H44N2O9 | 48.63± 16.88 | 65.27± 21.93 | 44.41± 11.85 | 78.79± 27.58 | 35.97± 9.85 | 75.5± 33.65 | 58.49± 34.98 |
| | Solamargine | C45H73NO15 | 16.33± 16.08 | 1.56± 0.93 | 2.89± 1.15 | 0.04± 0.04 | 3.25± 2.9 | 2.22± 0.77 | 1.21± 0.58 |
| | Solanidine base -2H + 1O, O-Hex-dHex-dHex | C45H71NO15 | 4± 2.55 | 0.61± 0.15 | 2.51± 0.73 | 0.18± 0.08 | 1.67± 1.08 | 1.57± 0.42 | 0.4± 0.26 |
| | Solasodine | C27H43NO2 | 19.81± 19.59 | 0.18± 0.09 | 0.12± 0.08 | 0.45± 0.31 | 0.39± 0.18 | 0.61± 0.35 | 0.2± 0.12 |
| | Solasonine | C45H73NO16 | 6.64± 5.02 | 1.03± 0.38 | 2.07± 0.64 | 2.08± 1.02 | 1.49± 1.31 | 0.92± 0.27 | 0.57± 0.24 |
| | Splendoline | C21H26N2O4 | 91.81± 15.34 | 175.09± 34.46 | 92.45± 13.08 | 289.95± 50.99 | 145.34± 33.69 | 104± 20.05 | 207.73± 32.14 |
| | Strictosamide | C26H30N2O8 | 20.11± 12.72 | 18.76± 10.04 | 2.74± 2.12 | 1.45± 1.45 | 2.07± 0.54 | 1.28± 0.92 | 1.6± 0.88 |
| | Subsessiline | C43H48N4O6 | 11.21± 3.03 | 30.9± 16.67 | 8.32± 1.68 | 14.92± 5.04 | 8.71± 2.76 | 17.76± 5.64 | 8.01± 3.47 |
| | Trigonelline | C7H7NO2 | 2149.19± 79.53 | 2170.92± 165.73 | 2105.82± 216.77 | 2099.89± 149.3 | 1816.79± 97.2 | 1964.57± 155.11 | 2025.3± 92.94 |
| | Vinpocetine | C22H26N2O2 | 12.73± 1.16 | 6.76± 1.85 | 13.82± 3.24 | 7.74± 1.24 | 5.71± 1.28 | 13.94± 3.49 | 10.11± 2.25 |
| | Xanthine | C5H4N4O2 | 23.96± 13.49 | 4.25± 1.62 | 12.15± 5.32 | 4.53± 1.18 | 1.69± 1.69 | 13.97± 5.03 | 7.13± 3.57 |

| | | | | | | | | | |
|---|---|---|---|---|---|---|---|---|---|
| Amines | Spermidine | C7H19N3 | 17.15± 2.58 | 36.61± 11.17 | 11.51± 2.23 | 31.11± 4.09 | 24.47± 6.17 | 15.78± 3.48 | 15.24± 4.71 |
| | Spermine | C10H26N4 | 31.68± 7.83 | 3.76± 1.62 | 0.11± 0.04 | 7.32± 7.25 | 1.62± 0.93 | 1.24± 0.69 | 1.51± 0.45 |
| | Tebuconazole | C16H22ClN3O | 4.19± 0.94 | 5.65± 0.98 | 5.26± 1.21 | 3.54± 1.04 | 2.08± 0.78 | 3.91± 0.92 | 3.57± 0.74 |
| | Tyramine | C8H11NO | 85.18± 8.05 | 132.9± 16.78 | 48.97± 6.68 | 42.07± 5.74 | 55.37± 4.36 | 57.53± 11.22 | 63.07± 7.78 |
| Amino acids and derivatives | 1-Aminocyclopropane-1-carboxylate | C4H7NO2 | 117.63± 13.31 | 93.46± 7.44 | 85.7± 2.62 | 77.49± 2.54 | 87.36± 5.92 | 83.55± 5.19 | 82.03± 7.67 |
| | 2-(4-aminotetrahydro-2H-pyran-4-yl) acetic acid | C7H13NO3 | 99.55± 31.89 | 34.57± 11.45 | 77.5± 15.37 | 44.02± 11.25 | 25.68± 4.54 | 71.42± 16.56 | 51.53± 12.52 |
| | Aphyllic Acid | C15H26N2O2 | 17.49± 8.81 | 13.71± 2.11 | 4.16± 1.26 | 3.15± 1.1 | 2.28± 1.02 | 3.88± 1.37 | 2.04± 0.93 |
| | Arginine | C6H14N4O2 | 17.71± 4.58 | 128.69± 67.07 | 21.29± 6.78 | 57.83± 28.73 | 19.23± 11.63 | 26.57± 4.21 | 14.13± 2.59 |
| | Asparagine | C4H8N2O3 | 125.94± 34.14 | 265.99± 211.45 | 89.12± 23.42 | 61.55± 17.72 | 41.9± 18.13 | 18.91± 3.45 | 40.91± 15.67 |
| | Aspartic acid | C4H7NO4 | 836.75± 42.51 | 653.35± 96.33 | 371.5± 37.67 | 348.26± 30.91 | 185.47± 28.93 | 219.74± 22.71 | 252.12± 33.71 |
| | Betaine | C5H11NO2 | 52.49± 6.01 | 59.5± 10.34 | 80.83± 33.62 | 21.13± 2.61 | 48.97± 14.31 | 35.69± 2.67 | 28.02± 3.8 |
| | Citrulline | C6H13N3O3 | 49.18± 7.61 | 33.97± 5.63 | 60.36± 8.09 | 29.69± 5.16 | 45.75± 9.02 | 50.24± 8.42 | 26.6± 5.66 |
| | CocamidopropylBetaine | C19H38N2O3 | 24.25± 2.4 | 25.61± 3.6 | 22.51± 2.77 | 19.65± 2.1 | 21.07± 2.44 | 17.35± 2.52 | 17.6± 2.59 |
| | Diphenylamine | C12H11N | 11.43± 1.45 | 21.04± 10.88 | 9.84± 1.61 | 8.96± 1.44 | 9.26± 1.43 | 7.8± 1.58 | 8.13± 1.72 |

| | | | | | | | | | |
|---|---|---|---|---|---|---|---|---|---|
| | DL-3-Aminoisobutyric acid | C4H9NO2 | 176.53± 16.91 | 157.72± 8.28 | 105.74± 7.43 | 87.56± 2.87 | 75.35± 5.21 | 88.44± 5.07 | 76.92± 6.49 |
| | feruloyltyramine | C18H19NO4 | 61.61± 7.3 | 156.05± 90.11 | 36.15± 21.54 | 25.79± 6.09 | 20.34± 5.15 | 16± 3.06 | 14.47± 4.23 |
| | gamma-Glutamyltyrosine | C14H18N2O6 | 15.45± 6.06 | 39.23± 15.83 | 28.38± 4.39 | 27.87± 4.29 | 17.46± 7.04 | 31.06± 5.72 | 13.1± 2.78 |
| | Glutamic acid | C5H9NO4 | 1266.33± 125.77 | 1042.1± 107.51 | 942.6± 32.87 | 889.76± 48.64 | 927.08± 63.98 | 942.28± 45.23 | 898.83± 96.51 |
| | Glutamine | C5H10N2O3 | 595.15± 62.96 | 687.95± 78.31 | 533.56± 98.34 | 513.51± 70.3 | 605.93± 143.21 | 394.09± 32.9 | 459.68± 48.46 |
| | Glutathione (oxidized) | C20H32N6O12S2 | 166.51± 16.19 | 79.26± 12.66 | 106.32± 23.95 | 73.79± 8.74 | 145.04± 22.14 | 76.38± 15.11 | 61.88± 13.77 |
| | Histamine | C5H9N3 | 210.48± 90.25 | 599.77± 113.47 | 565.62± 115.34 | 268.48± 68.27 | 467.74± 98.9 | 487.57± 42.07 | 302.09± 74.59 |
| | Isoleucine | C6H13NO2 | 434.2± 49.87 | 736.47± 186.94 | 401.41± 93.41 | 467.13± 62.38 | 399.89± 128.81 | 324.71± 52.34 | 288.65± 35.53 |
| | L-5-Oxoproline | C5H7NO3 | 216.86± 20.22 | 186.41± 9.59 | 119.29± 7.46 | 97.37± 2.72 | 76.85± 5.96 | 93.15± 4.89 | 82.62± 6.9 |
| | L-beta-Homoisoleucine | C7H15NO2 | 25.52± 11.25 | 17.96± 4.13 | 12.29± 2.71 | 16.83± 5.75 | 7.28± 2.34 | 18.46± 3.99 | 16.46± 4.58 |
| | L-glutamic acid-L-glutamine | C10H17N3O6 | 47.7± 7.2 | 53.16± 15.79 | 36.12± 6.65 | 44.13± 8 | 44.03± 14.28 | 35.96± 6.36 | 24.75± 5.4 |
| | L-Glutathione (oxidized form) | C20H32N6O12S2 | 207.06± 36.97 | 87.48± 17.92 | 30.36± 9.64 | 17.54± 3.58 | 13.51± 1.98 | 8.47± 1.14 | 8.57± 1.61 |
| | L-Phenylalanine | C9H11NO2 | 1709.99± 134.62 | 1930.33± 222.21 | 1541.42± 112.26 | 1643.49± 113.42 | 1528.65± 187.38 | 1738.82± 375.51 | 1187.51± 166.48 |
| | N, N-Dimethylarginine | C8H18N4O2 | 17.29± 1.51 | 25.55± 6.96 | 16.33± 2.18 | 27.59± 5.72 | 16.3± 5.01 | 19.71± 2.5 | 16.44± 2.49 |

| | | | | | | | | | |
|---|---|---|---|---|---|---|---|---|---|
| | N-Acetylarginine | C8H16N4O3 | 31.34± 2.55 | 64.35± 5.22 | 47.53± 8.69 | 38.08± 4.73 | 42.58± 4.31 | 43.27± 4.63 | 39.69± 3.59 |
| | N-Acetylhistidine | C8H11N3O3 | 33.95± 6.49 | 23.91± 6.09 | 24.41± 6.49 | 17.16± 3.11 | 17.99± 5.77 | 15.07± 2.79 | 12.23± 1.44 |
| | N-ACETYL-L-LEUCINE | C8H15NO3 | 49.61± 11.15 | 139.5± 69.15 | 70.5± 16.05 | 76.99± 11.13 | 23.9± 3.17 | 37.86± 4.32 | 27.18± 5.04 |
| | N-acetyl phenylalanine | C11H13NO3 | 94.88± 7.5 | 130± 37.07 | 82.66± 16.58 | 63.76± 8.74 | 25.18± 5.41 | 50.72± 8.11 | 39.69± 6.85 |
| | N-acetyltryptophan | C13H14N2O3 | 6.62± 0.47 | 12.29± 2.61 | 6.6± 1.76 | 4.93± 1.16 | 1.4± 0.45 | 3.21± 0.46 | 3.41± 0.45 |
| | N-acetyltyramine | C10H13NO2 | 54.3± 19.58 | 199.18± 74.46 | 99.85± 24.68 | 35.02± 14.1 | 59.34± 23.19 | 127.23± 33.91 | 99.04± 48.24 |
| | N-Fructosyl tyrosine | C15H21NO8 | 126.71± 22.41 | 123.26± 33.25 | 102.43± 17.14 | 67.19± 17.51 | 129.61± 16.23 | 58.88± 8.3 | 66.55± 12.17 |
| | O-Phosphothreonine | C4H10NO6P | 32.41± 7.88 | 9.63± 3.59 | 7.05± 1.36 | 5.84± 1.53 | 5.6± 1.23 | 2.51± 0.54 | 4.88± 1.81 |
| | Pantothenic acid | C9H17NO5 | 147.31± 37.09 | 206.66± 33.82 | 286.2± 38.84 | 146.24± 40.49 | 190.72± 22.2 | 208.96± 21.73 | 159.69± 15.43 |
| | Phenylalanine | C9H11NO2 | 2442.97± 317.14 | 4178.65± 489.25 | 2711.9± 347.82 | 2312.98± 286.09 | 1817.96± 340.9 | 2804± 573.91 | 1738.3± 391.33 |
| | Proline | C5H9NO2 | 1379.62± 97.51 | 1158.42± 204.74 | 1045.79± 92 | 996.39± 81.18 | 944.39± 84.23 | 1081.66± 43.43 | 827.4± 62.85 |
| | Pyroglutamic acid | C5H7NO3 | 61.71± 11.48 | 33.36± 2.86 | 39.07± 7 | 50.44± 4.78 | 30.23± 5.08 | 33.09± 6.5 | 28.99± 3.52 |
| | Quercetin 3-O-malonylglucoside | C24H22O15 | 6.55± 1.15 | 14.49± 3.03 | 22.73± 4.96 | 17.34± 1.96 | 13.63± 2.6 | 20.53± 2.9 | 17.65± 3.32 |
| | Serine | C3H7NO3 | 140.76± 30.32 | 111± 25.25 | 81.82± 5.63 | 64.2± 5.74 | 47.38± 8.44 | 58.25± 8.15 | 51.14± 4.98 |

| | | | | | | | | | |
|---|---|---|---|---|---|---|---|---|---|
| | Threonine | C4H9NO3 | 60.27± 10.34 | 26.2± 5.28 | 20.49± 1.04 | 13.63± 1.55 | 11.53± 1.56 | 10.52± 1.43 | 14.7± 1.44 |
| | Triethanolamine | C6H15NO3 | 23.55± 3.29 | 32.88± 6.7 | 23.71± 2.57 | 27.25± 6.48 | 19.87± 5.74 | 28.68± 6.58 | 29.19± 5.94 |
| | Tyrosine | C9H11NO3 | 247.22± 57.81 | 313.59± 69.38 | 212± 36.91 | 290.18± 39.66 | 127.13± 38.91 | 296.33± 87.66 | 189.63± 67.8 |
| | Valine | C5H11NO2 | 1.04± 0.16 | 0.97± 0.16 | 0.82± 0.14 | 0.82± 0.28 | 0.43± 0.1 | 0.8± 0.16 | 0.69± 0.22 |
| Anthocyans | Cyanidin-3,5-di-O-glucoside | C27H31O16 | 0.46± 0.22 | 0.49± 0.17 | 0.02± 0.02 | 0.12± 0.05 | 0.01± 0.01 | 0± 0 | 0± 0 |
| | Cyanidin-3-glucoside | C21H21O11 | 2.37± 0.35 | 2.12± 0.38 | 0.79± 0.21 | 0.76± 0.14 | 0.81± 0.32 | 0.49± 0.12 | 0.94± 0.31 |
| | Cyanidin-3-O-sambubioside | C26H29O15 | 0.24± 0.1 | 0.28± 0.17 | 0± 0 | 0.31± 0.09 | 0± 0 | 0± 0 | 0.06± 0.04 |
| | Malvidin-3-O-glucoside | C23H25O12 | 0.31± 0.09 | 0.15± 0.05 | 0.1± 0.06 | 0.08± 0.05 | 0± 0.05 | 0.09± 0.06 | 0.12± 0.05 |
| | Peonidin-3-O-glucoside | C22H23O11 | 6.45± 0.75 | 3.78± 0.63 | 1.2± 0.27 | 1.49± 0.29 | 0.48± 0.06 | 0.58± 0.08 | 0.42± 0.06 |
| Carbohydrates and derivatives | 1-Cinnamoylpyrrolidine | C13H15NO | 15.05± 1.51 | 27.18± 2.82 | 13.97± 3.69 | 11.04± 4 | 17.03± 8.23 | 14.9± 1.26 | 29.38± 6.56 |
| | 2-alpha-Mannobiose | C12H22O11 | 22.19± 5.66 | 7.17± 1.19 | 10.56± 7.92 | 1.98± 0.66 | 1.02± 0.16 | 2.58± 0.62 | 2± 0.53 |
| | 2'-Deoxyguanosine-5'-diphosphate | C10H15N5O10P2 | 66.81± 31.95 | 78.75± 16.54 | 80.92± 32.11 | 83.97± 53.81 | 20.27± 8.19 | 73.95± 49.73 | 76.93± 35.73 |
| | 5'-S-Methylthioadenosine | C11H15N5O3S | 456.66± 39.46 | 328.71± 26.66 | 303.45± 23.02 | 335.12± 38.49 | 304.37± 19.95 | 259.77± 16.44 | 273.5± 33.5 |
| | Adenine | C5H5N5 | 189.38± 5.27 | 140.95± 19.9 | 95.45± 10.19 | 92.29± 11.08 | 68.11± 9.6 | 79.61± 9.96 | 59.51± 10.83 |

| | | | | | | | | | |
|---|---|---|---|---|---|---|---|---|---|
| | Adenosine | C10H13N5O4 | 78.41± 19.05 | 92.44± 18.95 | 76.04± 9.62 | 105.53± 17.86 | 94.03± 26.94 | 57.97± 17.51 | 52.46± 4.41 |
| | Adenosine 3':5'-cyclic monophosphate | C10H12N5O6P | 62.83± 23.21 | 65.25± 14.43 | 38.14± 16.51 | 34.52± 11.87 | 22.54± 5.54 | 18.09± 8.73 | 32.29± 7.69 |
| | Adenosine 3'-monophosphate | C10H14N5O7P | 387.49± 102.76 | 257.25± 41.89 | 213.22± 75.07 | 135.38± 47.19 | 343.14± 108.13 | 101.29± 25.97 | 180.86± 82.96 |
| | Adenosine 5'-diphosphate | C10H15N5O10P2 | 12.6± 5.13 | 20.14± 3.97 | 21.92± 10.15 | 14.67± 8.72 | 7.74± 2.74 | 16.28± 10.83 | 21.82± 11.33 |
| | ADENOSINE 5'-DIPHOSPHATE-2 | C10H15N5O10P2 | 71.9± 34.53 | 69.93± 22.32 | 87.45± 35.11 | 28.42± 11.42 | 15.66± 10.95 | 79.67± 53.88 | 41.34± 35.61 |
| | Adenosine_Diphosphate | C10H15N5O10P2 | 5.54± 2.14 | 4.83± 1.15 | 10.53± 4.92 | 7.57± 3.57 | 5.78± 2.22 | 7.19± 3.86 | 9.31± 3.17 |
| | ADP | C10H15N5O10P2 | 47.8± 34.74 | 47.08± 13.51 | 49.76± 32.65 | 79.85± 56.91 | 28.71± 9.66 | 20.58± 16.04 | 77.74± 36.92 |
| | AMP | C10H14N5O7P | 449.92± 97.08 | 391.96± 72.98 | 329.39± 69.79 | 162.52± 32.63 | 329.81± 58.77 | 175.73± 37.68 | 156.87± 62.87 |
| | beta-D-glucopyranosiduronic acid | C21H26N2O4 | 61.72± 19.52 | 174.4± 34.31 | 3.89± 0.62 | 289.33± 50.86 | 144.7± 33.81 | 52.12± 33 | 207.59± 32.12 |
| | beta-Guanidinopropionic acid | C4H9N3O2 | 46.2± 9.42 | 83.45± 15.33 | 55.42± 16.3 | 60.68± 22.47 | 30.56± 5.2 | 70.31± 12.94 | 39.95± 11.55 |
| | beta-Nicotinamide adenine dinucleotide | C21H27N7O14P2 | 2.96± 2.8 | 0± 0 | 0.72± 0.72 | 2.04± 1.93 | 0.68± 0.6 | 3.98± 3.59 | 7.63± 4.77 |
| | D-Arabinose-5-phosphate disodium salt | C5H11O8P | 79.73± 5.66 | 50.71± 4.11 | 27.68± 7.37 | 23.98± 5.66 | 20.63± 2.97 | 11.41± 3.56 | 20.36± 3.33 |
| | Glucose-6-phosphate | C6H13O9P | 121.12± 43.09 | 79.15± 15.94 | 95.01± 20.65 | 97.43± 18.17 | 72.76± 22.85 | 59.77± 17.55 | 83.69± 18.4 |
| | Guanine | C5H5N5O | 324.07± 55.49 | 237.85± 32.31 | 265.43± 21.96 | 295.09± 45.73 | 234.79± 33.76 | 239.52± 50.95 | 199.28± 45.88 |

| | | | | | | | | | |
|---|---|---|---|---|---|---|---|---|---|
| | Guanosine | C10H13N5O5 | 540.45± 82.98 | 380.41± 56.72 | 395.7± 33.14 | 453.5± 71.89 | 340.96± 52.32 | 389.3± 70.91 | 311.11± 66.6 |
| | Guanosine 5'-diphosphate-D-mannose | C16H25N5O16P2 | 18.28± 10.71 | 2.26± 1.13 | 12.22± 7.49 | 12.58± 9.52 | 1.12± 1.8 | 17.73± 12.8 | 17.89± 7.96 |
| | Hypoxanthine | C5H4N4O | 39.66± 7.59 | 23.42± 4.13 | 41.86± 7.34 | 39.77± 5.88 | 24.25± 8.16 | 33.54± 8.36 | 28.29± 2.92 |
| | Inosine | C10H12N4O5 | 26.97± 6.41 | 19.61± 3.99 | 31.3± 5.48 | 26.69± 3.21 | 17.16± 6.13 | 25.41± 6.04 | 22.39± 2.84 |
| | isomaltulose | C12H22O11 | 261.17± 75.48 | 125.2± 55.1 | 191.74± 41.67 | 118.44± 31.42 | 105.65± 41.67 | 127.57± 29.88 | 126.8± 33.43 |
| | Mannose 1-phosphate | C6H13O9P | 1612.25± 225.08 | 951.34± 136.95 | 444.68± 159.14 | 386.95± 90.24 | 362.17± 49 | 167.08± 61.72 | 321.99± 107.61 |
| | Melezitose | C18H32O16 | 179.22± 32.39 | 97.19± 12.32 | 117.24± 21.64 | 113.73± 11.25 | 100.82± 13.03 | 97.02± 11.56 | 104.45± 13.1 |
| | N2-Methylguanosine | C11H15N5O5 | 9.72± 3.07 | 9.13± 3.37 | 12.69± 3.12 | 8.96± 3.21 | 6.73± 1.42 | 10.61± 2.7 | 8.79± 1.88 |
| | roseoside | C19H30O8 | 302.69± 48.6 | 467.98± 59.12 | 281.17± 35.31 | 432.64± 63.5 | 365.41± 80.54 | 301.89± 52.68 | 486.71± 101.39 |
| | Sorbitol | C6H14O6 | 23.74± 8.06 | 30.71± 7.96 | 30.58± 10.34 | 18.01± 3.04 | 10.13± 3.79 | 15.31± 3.07 | 11.87± 1.15 |
| | Swertiamarin | C16H22O10 | 0.12± 0.05 | 0.08± 0.05 | 0± 0 | 0± 0 | 0± 0 | 0± 0 | 0± 0 |
| | trans-piceid | C20H22O8 | 6.38± 0.97 | 4.61± 0.78 | 1.94± 0.33 | 2.58± 0.72 | 1.09± 0.22 | 1.55± 0.42 | 0.47± 0.08 |
| | Trehalose | C12H22O11 | 2768± 310.99 | 1825.52± 152.87 | 853.06± 39.41 | 643.8± 45.93 | 341.42± 39.05 | 510.08± 30.57 | 511.66± 80.82 |
| | UDP-D-glucose | C15H24N2O17P2 | 632.29± 276.37 | 97.7± 29.46 | 257.26± 156.04 | 135.03± 58.15 | 6.33± 22.53 | 170.1± 99.8 | 56.37± 17.47 |

| | | | | | | | | | |
|---|---|---|---|---|---|---|---|---|---|
| | Uracil | C4H4N2O2 | 107.61± 21.64 | 69.49± 7.73 | 68.46± 8.93 | 80.44± 11.16 | 63.44± 11.87 | 54.78± 10.51 | 56.94± 12.28 |
| | Uridine | C9H12N2O6 | 43.93± 3.11 | 38.6± 5.57 | 18.15± 0.88 | 13.35± 2.41 | 9.61± 0.86 | 8.35± 0.62 | 8.14± 1.56 |
| | Uridine 5'-diphospho-D-glucose | C15H24N2O17P2 | 321.86± 141.35 | 97.69± 29.45 | 257.25± 156.05 | 135.02± 58.15 | 6.32± 22.53 | 170.08± 99.8 | 56.36± 17.46 |
| | Xanthosine | C10H12N4O6 | 105.38± 19.55 | 82.14± 9.55 | 89.61± 13.04 | 61.89± 8.35 | 33.05± 6.76 | 69.39± 11.77 | 73.18± 27.51 |
| Carboxylic acids and derivatives | 2,8-Quinolinediol | C9H7NO2 | 7.36± 4.7 | 2.51± 2.51 | 4.56± 2.93 | 2.56± 1.65 | 2.71± 2.63 | 4.63± 2.96 | 4.23± 2.75 |
| | 2-acetoxy-4-pentadecylbenzoic acid | C24H38O4 | 84.64± 9.88 | 86.09± 11.54 | 77.55± 10.14 | 60.95± 12.88 | 65.79± 14.67 | 61.53± 12.95 | 67.11± 8.42 |
| | 2-Hydroxy-4-methylpentanoic acid | C6H12O3 | 1.1± 0.64 | 8.22± 3.49 | 0.86± 0.27 | 0.63± 0.09 | 0.37± 0.08 | 0.55± 0.21 | 0.45± 0.18 |
| | 2-Isopropylmalic acid | C7H12O5 | 492.09± 125.83 | 882.43± 302.91 | 252.29± 27.73 | 262.98± 40.61 | 209.37± 38.94 | 164.08± 53.65 | 117.81± 15.88 |
| | 2-Oxobutyric acid | C4H6O3 | 9.64± 0.9 | 29.36± 5.37 | 12.06± 2.1 | 10.78± 2.64 | 8.23± 2.87 | 14.31± 1.97 | 11.01± 1.5 |
| | 3-(4-HYDROXY-3,5-DIMETHOXYPHENYL)-2-PROPENOIC ACID | C11H12O5 | 8.81± 1.23 | 22.38± 2.72 | 36.33± 4.92 | 27.89± 3.84 | 17.51± 2.58 | 39.27± 7.76 | 20.11± 3.14 |
| | 3-(4-HYDROXYPHENYL)PROP-2-ENOIC ACID | C9H8O3 | 100.14± 12.7 | 92.92± 18.79 | 84.13± 11.96 | 58.32± 13.71 | 104.09± 8.56 | 61.96± 12.64 | 63.25± 9.43 |
| | 3-(Benzoyloxy)-2-hydroxypropyl beta-D-glucopyranosiduronic acid | C16H20O10 | 82.89± 13.78 | 54.9± 6.75 | 73.21± 16.48 | 62.5± 6.73 | 39.73± 2.47 | 75.21± 13.03 | 63.79± 11.8 |
| | 3-Indolepropionic acid | C11H11NO2 | 27.79± 8.71 | 34.02± 7.08 | 21.88± 6.22 | 21.57± 7.3 | 10.12± 1.93 | 31.69± 5.96 | 19.09± 7.42 |

| | | | | | | | | | |
|---|---|---|---|---|---|---|---|---|---|
| | 3-phenyl lactic acid | C9H10O3 | 0.1± 0.06 | 66.39± 19.13 | 0.16± 0.05 | 0.08± 0.04 | 0.03± 0.02 | 0.02± 0.02 | 0.12± 0.04 |
| | 4-Hydroxyquinoline-2-carboxylic acid | C10H7NO3 | 238.6± 22.58 | 166.36± 18.4 | 128.12± 18.2 | 70.38± 10.21 | 76.67± 7.17 | 83.41± 8.64 | 91.58± 5.9 |
| | Benzoic acid | C7H6O2 | 4.41± 0.91 | 3.34± 1.39 | 1.92± 0.7 | 0.76± 0.34 | 2.05± 0.66 | 2.62± 0.84 | 3.58± 1.35 |
| | cis-Aconitate | C6H6O6 | 869.21± 62.44 | 1146.29± 82.57 | 796.68± 63.39 | 602.84± 65.95 | 483.63± 40.61 | 485.35± 39.97 | 435.49± 29.83 |
| | Citric acid | C6H8O7 | 9041.11± 1396.46 | 9522.81± 2199.61 | 7065.11± 1016.65 | 7780.48± 1081.04 | 6118.67± 991.11 | 6226.08± 795.8 | 5909.42± 406.64 |
| | Diethyl phthalate | C12H14O4 | 62.23± 8.72 | 66.07± 12.76 | 57.41± 10.29 | 50.78± 9.36 | 54.77± 7.86 | 44.85± 8.44 | 46.88± 8.58 |
| | Dioctyl Phthalate | C24H38O4 | 86.42± 9.89 | 87.59± 11.93 | 81.13± 11.09 | 76.46± 9.8 | 69.43± 15.56 | 64.35± 13.48 | 62.67± 13.41 |
| | Ethylenediaminetetraacetic acid EDTA | C10H16N2O8 | 4.96± 1.32 | 6.26± 1.42 | 5.84± 1.75 | 4.98± 1.53 | 6.22± 1.53 | 4.13± 1.04 | 3.71± 1.02 |
| | Fumaric acid | C4H4O4 | 4035.82± 108.71 | 4542.32± 446.76 | 3050.23± 251.71 | 2401.29± 290.16 | 1434.91± 236.44 | 2095.87± 111.44 | 1837.03± 119.27 |
| | Kynurenic acid | C10H7NO3 | 238.67± 22.61 | 166.38± 18.41 | 128.25± 18.15 | 70.32± 10.2 | 76.57± 7.17 | 83.32± 8.63 | 91.47± 5.92 |
| | L-Glutamic acid | C5H9NO4 | 1929.19± 204.91 | 1605.3± 88.32 | 921.76± 71.05 | 698.85± 24.8 | 534.34± 41.29 | 670.25± 32.62 | 586.94± 47.04 |
| | L-kynurenine | C10H12N2O3 | 34.37± 4.29 | 47.16± 3.1 | 45.33± 6.81 | 42.24± 4.23 | 35.36± 6.01 | 43.51± 4.82 | 36.43± 6.15 |
| | L-Saccharopine | C11H20N2O6 | 17.89± 1.87 | 22.13± 6.16 | 13.81± 1.31 | 13.57± 1.32 | 10.19± 1.11 | 49.12± 33.78 | 14.2± 1.46 |
| | Methyl nicotinic acid | C7H7NO2 | 101.3± 10.97 | 72.88± 15.96 | 54.65± 4.8 | 102.31± 16.58 | 59.57± 4.04 | 50.81± 5.56 | 49.6± 4.84 |

| | | | | | | | | | |
|---|---|---|---|---|---|---|---|---|---|
| | MUCIC ACID | C6H10O8 | 1047.47± 159.75 | 759.22± 147.54 | 337.58± 76.01 | 237.21± 49.66 | 179.83± 23.03 | 192.62± 19.75 | 179.09± 20.18 |
| | Nicotinic acid | C6H5NO2 | 11.87± 3.72 | 9.38± 1.57 | 7.62± 1.7 | 13.95± 3.63 | 6.46± 0.65 | 7.84± 1.74 | 6.32± 2.27 |
| | Phthalic anhydride | C8H4O3 | 479.53± 54.15 | 391.46± 38.13 | 387.17± 29.99 | 348.67± 36.41 | 333.82± 27.8 | 321.79± 40.03 | 357.34± 45.59 |
| | Quercetin-4'-glucoside | C21H20O12 | 105.28± 16.16 | 280.78± 48.36 | 268.14± 40.39 | 227.73± 39.76 | 238.69± 49.87 | 210.29± 17.31 | 243± 46.62 |
| | Salicylic acid | C7H6O3 | 299.59± 71.67 | 236± 23.36 | 101.87± 9.35 | 106.5± 13.72 | 90.52± 7.94 | 91.48± 8.41 | 87.96± 15.2 |
| | Trifloxystrobin | C20H19F3N2O4 | 5.58± 1.6 | 6.57± 1.39 | 5.39± 1.1 | 4.54± 1.08 | 5.33± 1.11 | 4.52± 1.25 | 4.96± 1.34 |
| | Tryptophan | C11H12N2O2 | 4895.94± 640.23 | 7477.03± 1392.95 | 3974.2± 975.25 | 4248.6± 1375.89 | 1902.42± 341.64 | 4418.96± 630.39 | 2394.45± 506.25 |
| | Vanillic acid | C8H8O4 | 413.83± 33.74 | 292.68± 13.08 | 257.91± 31.82 | 150.04± 11.46 | 94.81± 19.24 | 168.41± 17.12 | 140.61± 12.4 |
| | Xanthurenic Acid | C10H7NO4 | 33.47± 10.56 | 21.6± 5.64 | 36.4± 12.67 | 17.45± 5.54 | 21.47± 8.42 | 30.22± 9.18 | 63.31± 35.34 |
| Cholines | Acetylcholine | C7H16NO2 | 73.25± 5.78 | 99.84± 26.14 | 61.53± 11.78 | 66.61± 17.99 | 70.06± 22.93 | 36.48± 8.06 | 50.46± 10.57 |
| Fatty acid and derivatives | 9,12,15-octadecatrienoic acid | C18H30O2 | 683.35± 186.94 | 711.88± 97.02 | 786.03± 147.35 | 478.07± 115.92 | 837.51± 85.39 | 595.64± 88.08 | 450.83± 145.83 |
| | Acaranoic acid | C17H30O4 | 0.2± 0.09 | 0.06± 0.03 | 0± 0 | 0± 0 | 0± 0 | 0± 0 | 0± 0 |
| | Avocadyne Acetate | C19H34O4 | 48.29± 10.88 | 59.52± 15.51 | 41.43± 8.98 | 37.93± 9.5 | 38.89± 8.73 | 30.84± 6.32 | 27.16± 9.01 |
| | Azelaic acid | C9H16O4 | 68.53± 9.52 | 48.11± 7.18 | 32.37± 1.45 | 18.69± 2.8 | 19.97± 2.99 | 22.62± 2.32 | 14.13± 1.33 |

| | | | | | | | | | |
|---|---|---|---|---|---|---|---|---|---|
| | beta-D-Glucopyranoside | C21H36O10 | 7.5± 1.12 | 1.6± 0.72 | 0.9± 0.34 | 1.84± 0.43 | 1.01± 0.32 | 1.6± 0.21 | 0.86± 0.14 |
| | Heptadecanoic acid | C17H34O2 | 14.65± 0.96 | 11.08± 0.66 | 5.24± 0.4 | 3.38± 0.14 | 2.56± 0.19 | 2.65± 0.15 | 2.31± 0.2 |
| | hexadecanedioic acid | C16H30O4 | 4.12± 0.64 | 2.83± 0.37 | 2.77± 0.4 | 2.48± 0.37 | 1.83± 0.41 | 2.1± 0.33 | 2.63± 0.49 |
| | Linoelaidic acid | C18H32O2 | 44.11± 3.13 | 31.36± 3.14 | 13.86± 2.58 | 6.93± 1.57 | 4.76± 1.26 | 5.56± 0.99 | 4.85± 0.71 |
| | Linoleic acid | C18H32O2 | 1007.19± 157.12 | 618.26± 62.99 | 280.49± 30.96 | 158.58± 27.42 | 135.43± 11.92 | 152.35± 15.14 | 118.93± 22.11 |
| | linolenic acid | C18H30O2 | 683.41± 186.95 | 712.02± 96.92 | 786.43± 147.39 | 477.9± 115.78 | 837.99± 85.49 | 595.77± 88.05 | 450.85± 145.83 |
| | Mesaconic acid | C5H6O4 | 65.14± 8.15 | 69.79± 17.2 | 44.28± 4.29 | 32.71± 4.21 | 24.72± 3.21 | 32.63± 3.67 | 28.56± 6.01 |
| | methyl palmitate | C17H34O2 | 38.46± 10.05 | 20.58± 1.29 | 38.3± 16.36 | 32.57± 6.25 | 22.27± 5.25 | 28.41± 10.29 | 26.53± 6.37 |
| | Palmitic acid (NMR) | C16H32O2 | 260.53± 27.93 | 204.61± 18.91 | 87.62± 10.94 | 46.03± 3.37 | 41.09± 5 | 40.61± 2.36 | 34.84± 4.16 |
| | Palmitoleic acid | C16H30O2 | 102.42± 5.32 | 96.87± 5.99 | 43.77± 5.29 | 27.91± 1.97 | 14.06± 2.3 | 22.52± 1.6 | 19.92± 2.01 |
| | Stearic acid | C18H36O2 | 48.92± 3.26 | 40.01± 1.95 | 20.21± 1.51 | 14.33± 0.91 | 12.51± 0.81 | 13.47± 0.62 | 11.74± 0.79 |
| | Trans-Vaccenic acid | C18H34O2 | 117.16± 13.65 | 82.24± 6.14 | 36.81± 3.29 | 20.48± 1.98 | 17.99± 2.3 | 17.48± 1.21 | 19.35± 3.29 |
| | y-Linolenic acid | C18H30O2 | 3121.51± 536.94 | 1983.01± 294.54 | 772.68± 83.96 | 420.11± 75.26 | 332± 40.47 | 445.21± 49.66 | 319.77± 42.38 |
| Flavins | (-)-RIBOFLAVIN | C17H20N4O6 | 85.7± 10.57 | 91.5± 3.67 | 82.76± 6.48 | 99.72± 17.28 | 84.14± 6.73 | 74.05± 14.58 | 51.79± 10.45 |

| | | | | | | | | | |
|---|---|---|---|---|---|---|---|---|---|
| Furanochromones | 5-O-methylvisammioside | C22H28O10 | 39.59± 13.44 | 6.28± 3.49 | 9.7± 4.08 | 16.11± 5.92 | 4.65± 2.86 | 10.36± 6.19 | 11.22± 5.03 |
| Indoles | Indole-3-carbinol | C9H9NO | 0.36± 0.12 | 0.37± 0.22 | 0.43± 0.23 | 0.33± 0.1 | 2.8± 1.71 | 0.79± 0.6 | 0.45± 0.11 |
| Indolizidines | Corynoxine | C22H28N2O4 | 1.65± 0.18 | 39.38± 20.57 | 1.41± 0.45 | 2.12± 0.31 | 3.79± 1.74 | 1.28± 0.3 | 10.17± 2.35 |
| Lignans and derivatives | Arctigenin | C21H24O6 | 2.84± 0.3 | 1.56± 0.19 | 0.74± 0.1 | 0.82± 0.05 | 0.35± 0.07 | 0.58± 0.04 | 0.43± 0.02 |
| | Eleutheroside E | C34H46O18 | 3.45± 0.74 | 4.25± 1.45 | 3.16± 1.64 | 2.76± 0.83 | 0.25± 0.5 | 1.54± 0.41 | 0.87± 0.38 |
| | Secoisolariciresinol | C20H26O6 | 0.88± 0.08 | 1.02± 0.17 | 0.59± 0.14 | 0.17± 0.06 | 0.06± 0.07 | 0.33± 0.07 | 0.09± 0.06 |
| Lipids | Dehydrophytosphingosine | C18H37NO3 | 50.25± 27.15 | 103.97± 7.89 | 73.03± 30.51 | 77.79± 26.89 | 67.39± 18.08 | 99.43± 32.45 | 98.33± 48.67 |
| | Phosphocholine | C5H15NO4P | 9.68± 1.32 | 5.43± 0.56 | 6.85± 1.09 | 6.23± 1.92 | 3.82± 0.44 | 6.97± 2.05 | 6.27± 1.36 |
| | quercetin-3-O-glc-1-3-rham-1-6-glucoside | C33H40O21 | 195.76± 30.7 | 627.59± 175.15 | 166.36± 58.26 | 179.78± 54.23 | 406.68± 110.48 | 57.54± 15.23 | 29.87± 6.44 |
| | sn-Glycero-3-phosphocholine | C8H21NO6P | 77.68± 15.09 | 34.97± 1.65 | 43.71± 5.99 | 52.63± 17.41 | 32.43± 3.97 | 36.22± 3.29 | 29.81± 5.67 |
| Macrolides and derivatives | curvularin | C16H20O5 | 0.11± 0.04 | 0.12± 0.05 | 0± 0 | 0± 0 | 0± 0 | 0± 0 | 0± 0 |
| | dihydroalbocycline | C18H30O4 | 41.45± 18.7 | 44.48± 5.94 | 29.63± 6.12 | 44.51± 14.71 | 27.88± 10.77 | 37.77± 9.37 | 20.69± 7.57 |
| | Gardnutine | C20H22N2O2 | 13.72± 2.33 | 16.86± 2.36 | 8.71± 1.26 | 9.01± 1.94 | 4.62± 1.67 | 8.92± 1.16 | 7.34± 1.65 |
| Oxanes | Cochlioquinone A | C30H44O8 | 808.76± 144.62 | 511.59± 96.67 | 172.02± 19.27 | 100.11± 8.8 | 65.23± 12.54 | 80.17± 8.69 | 75.78± 6.95 |

| | | | | | | | | | |
|---|---|---|---|---|---|---|---|---|---|
| | 1-O-beta-D-Glucopyranosyl sinapate | C17H22O10 | 41.11± 8.62 | 24.48± 2.51 | 12.67± 3.56 | 17.1± 6.43 | 3.91± 1.38 | 13.67± 5.58 | 10.6± 4.96 |
| | 2-Glucosyloxy-4-methoxy cinnamic acid | C16H20O9 | 2.16± 0.59 | 5.04± 1.2 | 7.08± 2.45 | 5.85± 1.66 | 4.93± 0.92 | 5.53± 1.44 | 7.66± 1.13 |
| | 3,4-di-O-caffeoylquinic acid | C25H24O12 | 56.79± 5.8 | 24.81± 3.28 | 14.22± 2.19 | 12.29± 2.93 | 9.63± 1.88 | 6.54± 0.84 | 5.36± 0.87 |
| | 3-Deoxycaryoptinol | C24H34O7 | 10.45± 0.76 | 10.43± 1.23 | 6.62± 0.86 | 3.63± 0.49 | 1.23± 0.74 | 5.3± 0.48 | 2.99± 0.26 |
| | 3-Hydroxy-4-methoxy cinnamic acid (isoferulic acid) | C10H10O4 | 245.93± 9.84 | 103.18± 24.68 | 162.63± 23.33 | 85.02± 18.75 | 94.8± 18.36 | 105.98± 11 | 79.86± 14.05 |
| | 3-Hydroxycinnamic acid | C9H8O3 | 195.57± 20.77 | 171.44± 32.45 | 72.73± 10.25 | 47.24± 11.53 | 60.61± 7.16 | 36.41± 6.37 | 29.97± 5.42 |
| | 3-O-Feruloylquinic acid | C17H20O9 | 2512.86± 349.14 | 928.49± 117.92 | 488.57± 41.29 | 451.57± 75.66 | 218.5± 26.69 | 249.13± 10.4 | 272.49± 28.37 |
| | 4',5,7-Trihydroxy-6,8-diprenylisoflavone | C25H26O5 | 3.44± 0.3 | 2.17± 0.16 | 3.27± 0.95 | 2.47± 0.37 | 2± 0.45 | 2.51± 0.39 | 2.28± 0.32 |
| | 4-Caffeoylquinic acid | C16H18O9 | 10385.67 ± 1661.77 | 6025.36± 665.96 | 2059.18± 180.12 | 1938.37± 633.09 | 1809.8± 274.2 | 1127.04± 102.78 | 893.26± 70.86 |
| | 6,7-Dihydroxycoumarin | C9H6O4 | 6.05± 1.87 | 72.65± 21.96 | 15.8± 8.67 | 39.48± 16.84 | 10.6± 4.53 | 12.79± 3.63 | 13.39± 7.38 |
| | Apigenin-7-O-glucoside | C21H20O10 | 4.93± 1.13 | 2.03± 0.75 | 1.79± 0.87 | 1.24± 0.42 | 0.23± 0.39 | 1.01± 0.36 | 0.95± 0.1 |
| | Benzoic acid + 1O, 2MeO, O-Hex | C15H20O10 | 407.25± 46.79 | 288.45± 29.13 | 179.35± 19.04 | 157.91± 29.08 | 70.64± 11.87 | 78.21± 15.64 | 33.93± 4.97 |
| | Benzoic acid + 2O, O-Hex | C13H16O9 | 12954.86 ± 786.81 | 8593.74± 1238.53 | 4512.09± 1112.99 | 4717.8± 837.97 | 1022.15± 377.68 | 3459.78± 774.17 | 4059.15± 1048.35 |
| | Bergenin | C14H16O9 | 1.55± 0.46 | 0.82± 0.26 | 0.51± 0.19 | 0.64± 0.13 | 0.09± 0.06 | 0.18± 0.06 | 0.26± 0.05 |

| | | | | | | | | | |
|---|---|---|---|---|---|---|---|---|---|
| | Biochanin-7-O-glucoside | C22H22O10 | 3.06± 0.62 | 2.15± 0.9 | 0.66± 0.16 | 0.67± 0.12 | 0.14± 0.08 | 0.26± 0.08 | 0.23± 0.03 |
| | Caffeic acid | C9H8O4 | 57.63± 7.93 | 41.23± 6.91 | 62.24± 8.75 | 36.96± 9.37 | 55.55± 3.57 | 41.62± 9.34 | 38.73± 7.96 |
| | Caffeic acid hexoside | C15H18O9 | 1229.03± 127.81 | 876.17± 171.9 | 483.45± 82.16 | 269.63± 79.5 | 269.65± 31.39 | 205.81± 55.84 | 142.54± 34.44 |
| | Caffeoyl putrescin | C13H18N2O3 | 329.71± 279.63 | 260.04± 124.83 | 71.36± 34.49 | 16.78± 7.83 | 21.17± 9.24 | 32.33± 8.69 | 106.05± 63.44 |
| | Caffeoyl quinic acid | C16H18O9 | 13572.19 ± 2201.83 | 6119.25± 1569.16 | 4965.55± 678.44 | 1342.58± 277.73 | 1693.69± 254.42 | 1815.71± 245.62 | 1377.97± 281.76 |
| | Caffeoylcholine | C14H20NO4 | 14.03± 3.32 | 37.4± 4.27 | 22.77± 5.86 | 26.67± 4.47 | 16.37± 4.08 | 16.38± 4.66 | 16.02± 1.79 |
| | CHLOROGENIC ACID | C16H18O9 | 6588.81± 752.25 | 4419.92± 716.66 | 1787.25± 132.8 | 2123.61± 561.31 | 1773.93± 285.93 | 1009.96± 131.17 | 792.14± 85.85 |
| | Citrusin | C16H22O7 | 0.92± 0.12 | 1.1± 0.34 | 1.07± 0.27 | 1.36± 0.66 | 1.62± 0.13 | 1.04± 0.36 | 1.55± 0.23 |
| | Coumarin + 1O + 1MeO, O-Hex-Hex | C22H28O14 | 1328.45± 181.47 | 404.6± 26.67 | 179.24± 3.86 | 134.06± 21.45 | 73.63± 9.14 | 76.33± 11.32 | 55.72± 9.89 |
| | Coumaroyl Hexoside | C15H18O8 | 1696.3± 183.72 | 1386.74± 278.16 | 463.42± 73.01 | 312.41± 82.73 | 359.8± 50.89 | 195.98± 42.36 | 157.37± 32.38 |
| | Coumaroyl quinic acid | C16H18O8 | 650.74± 92.73 | 727.19± 74.55 | 330.6± 46.66 | 216.08± 34.46 | 179.92± 21.16 | 159.31± 31.26 | 149.12± 31.08 |
| | Coumaroyl tyramine | C17H17NO3 | 53.24± 6.42 | 464.86± 311.24 | 189.39± 146.29 | 161.86± 85.68 | 71.11± 6.91 | 44.53± 10.71 | 48.22± 8.71 |
| | Cupressuflavone | C30H18O10 | 0.55± 0.17 | 0.41± 0.09 | 0.98± 0.57 | 0.43± 0.26 | 0.26± 0.15 | 0.11± 0.08 | 1.23± 0.45 |
| | Cyanidin | C15H11O6 | 0.7± 0.13 | 0.23± 0.06 | 0.13± 0.07 | 0.12± 0.04 | 0.04± 0.04 | 0.05± 0.03 | 0.06± 0.04 |

| | | | | | | | | | |
|---|---|---|---|---|---|---|---|---|---|
| | Cyanidin 3-(2G-glucosylrutinoside) | C33H41O20 | 166.61± 32.15 | 79.03± 20.9 | 19.59± 6.05 | 23.09± 3.81 | 64.22± 16.83 | 5.67± 1.58 | 5.47± 0.81 |
| | Cyanidin-3-O-(2"-O-beta-xylopyranosyl-beta-glucopyranoside) | C26H29O15 | 5.44± 1.02 | 2.46± 0.35 | 2.07± 0.24 | 2.49± 0.67 | 2.31± 0.12 | 1.74± 0.47 | 3.05± 0.39 |
| | Cyclohexanecarboxylic acid | C16H18O8 | 141.34± 37.4 | 96.84± 30.15 | 155.23± 21.77 | 113.73± 19.7 | 131.6± 25.58 | 108.59± 42.6 | 128.01± 27.07 |
| | Cynarin | C25H24O12 | 56.88± 5.74 | 24.84± 3.28 | 14.18± 2.18 | 12.3± 2.94 | 9.63± 1.88 | 6.54± 0.84 | 5.36± 0.87 |
| | D-(-)-Quinic acid | C7H12O6 | 8051.7± 1670.78 | 6326.07± 1329.37 | 2366.46± 706.78 | 2823.36± 706.05 | 2843.34± 580.91 | 1192.65± 317.77 | 1935.99± 335.41 |
| | Daphnetin | C9H6O4 | 40.98± 4.28 | 56.16± 3.94 | 22.89± 2.38 | 12.96± 1.8 | 15.48± 1.14 | 14.97± 1.98 | 10.37± 0.87 |
| | Di-dihydro caffeoyl spermidine | C25H35N3O6 | 43.38± 14.15 | 23.54± 13.75 | 85.94± 44.51 | 13.72± 2.74 | 24.63± 12.35 | 61.72± 35.26 | 67.51± 22.41 |
| | Epigallocatechin | C15H14O7 | 358.19± 153.18 | 201.31± 65.9 | 283.56± 51.22 | 70.88± 47.48 | 294.07± 86.75 | 203.02± 67.22 | 154.29± 68.43 |
| | epsilon-Viniferin | C28H22O6 | 17.35± 4.93 | 91.85± 13.65 | 69.43± 13.01 | 107.5± 16.76 | 63.3± 18.45 | 62.92± 15.96 | 73.84± 10.5 |
| | Eriodictyol-7-O-neohesperidoside | C27H32O15 | 224.94± 31.01 | 136.22± 19.38 | 39.53± 5.58 | 45.38± 5.85 | 20.12± 1.4 | 23.92± 4.88 | 18.5± 3.12 |
| | Eriodictyol-7-O-rutinoside | C27H32O15 | 95.65± 21.54 | 56.16± 11.99 | 63.84± 7.23 | 53.66± 9.15 | 57.45± 6.39 | 50.69± 7.82 | 52.3± 6.27 |
| | Esculin | C15H16O9 | 30± 5.91 | 33.88± 3.21 | 33.8± 3.07 | 14.85± 1.91 | 27.28± 2.24 | 26.23± 1.38 | 26.65± 4.58 |
| | Eupatilin | C18H16O7 | 31.16± 2.7 | 20.02± 3.3 | 10.73± 1.51 | 6.76± 1.75 | 5.82± 0.55 | 4.72± 1.02 | 3.53± 0.73 |
| | Ferulic acid | C10H10O4 | 116.96± 17.49 | 41± 11.83 | 64.2± 20.14 | 69.97± 23.51 | 22.66± 2.97 | 92.36± 25.83 | 64.04± 19.09 |

| | | | | | | | | | |
|---|---|---|---|---|---|---|---|---|---|
| | Feruloyl tyramine | C18H19NO4 | 261.09± 33.97 | 662.49± 314.46 | 318.69± 197.88 | 203.85± 52.66 | 192.34± 52.32 | 159.98± 38.04 | 123.88± 14.65 |
| | Flavone base + 4O, O-MalonylHex | C24H22O14 | 74.27± 15.55 | 20.59± 3.52 | 15.5± 2.1 | 16.1± 1.42 | 6.06± 2.48 | 7.36± 1.91 | 14.1± 1.82 |
| | Flavonol base + 4O, O-Hex-dHex, O-Hex | C33H40O21 | 73.2± 15.88 | 220.22± 33.61 | 38.43± 15.48 | 30.85± 6.48 | 53.2± 15.59 | 8.87± 1.55 | 2.99± 0.93 |
| | Glabrol | C25H28O4 | 35.66± 4.14 | 35.17± 2.78 | 33.05± 1.56 | 28.83± 2.75 | 26.65± 2.58 | 26.11± 1.27 | 21.87± 1.72 |
| | Hyperoside | C21H20O12 | 68.25± 11.97 | 201.99± 30.01 | 97.38± 10.26 | 90.5± 16.19 | 62.81± 6.24 | 64.42± 7.98 | 67.25± 12.79 |
| | irigenin | C18H16O8 | 26.22± 3.68 | 32.81± 6.61 | 40.19± 12.76 | 27.03± 5.42 | 40.37± 6.78 | 23.18± 8.25 | 32.18± 8.13 |
| | Isorhamnetin | C16H12O7 | 155.47± 14.81 | 183.29± 35.73 | 282.78± 27.6 | 290.26± 40.46 | 189.31± 24.53 | 240.75± 40.48 | 260.09± 17.77 |
| | Isorhamnetin 3-galactoside | C22H22O12 | 12.25± 1.65 | 18.88± 2.2 | 9.72± 1.78 | 7.72± 1.14 | 3.55± 0.69 | 5.68± 0.83 | 6.9± 1.58 |
| | isorhamnetin-3-glucoside-4'-glucoside (Isorhamnetin 3,4'-diglucoside) | C28H32O17 | 49.19± 5.4 | 69.21± 9.68 | 58.35± 11.96 | 41.3± 5.54 | 17.51± 5.9 | 40.43± 2.37 | 21.82± 1.41 |
| | isorhamnetin-3-O-glucoside | C22H22O12 | 308.22± 59.25 | 298.45± 46.06 | 225.2± 25.93 | 194.7± 32.11 | 96.63± 26.39 | 123.84± 24.61 | 103.09± 7.54 |
| | Isorhamnetin-3-O-rutinoside | C28H32O16 | 21.53± 3.75 | 13.8± 2.98 | 13.34± 3.14 | 6.24± 0.83 | 4.23± 0.92 | 4.21± 0.92 | 4.03± 1.34 |
| | Kaempferol | C15H10O6 | 495.39± 70.75 | 258.92± 36.67 | 297.06± 94.1 | 291.75± 41.91 | 271.78± 29.89 | 166.19± 36.28 | 294.06± 26.92 |
| | Kaempferol 3-O-gentiobioside | C27H30O16 | 8.69± 3.86 | 0.78± 0.3 | 0.36± 0.15 | 0.61± 0.26 | 0.25± 0.07 | 0.21± 0.07 | 0.22± 0.07 |
| | Kaempferol 3-O-sophoroside | C27H30O16 | 669.43± 55.06 | 439.98± 46.34 | 173.61± 15.59 | 155.6± 28.48 | 95.4± 14.81 | 78.14± 17.87 | 114.27± 12.28 |

| | | | | | | | | | |
|---|---|---|---|---|---|---|---|---|---|
| | Kaempferol-3-Glucoside-3"-Rhamnoside | C27H30O15 | 256.9± 33.64 | 146.88± 51.05 | 137.64± 52.35 | 80.54± 9.88 | 114.55± 19.46 | 60.64± 19.17 | 57.86± 9.21 |
| | Kaempferol-3-O-glucoside | C21H20O11 | 34.98± 3.97 | 5.31± 0.74 | 2.92± 1.43 | 3.91± 0.49 | 1.96± 0.41 | 0.99± 0.26 | 4.07± 0.78 |
| | kaempferol-3-O-rutinoside | C27H30O15 | 0± 0 | 0± 0 | 0± 0 | 0± 0 | 0± 0 | 0± 0 | 0± 0 |
| | Kaempferol-4'-glucoside | C21H20O11 | 1288.39± 192.23 | 480.81± 78.6 | 554.58± 130.76 | 707.76± 90.19 | 518.22± 83.28 | 327.61± 63.9 | 764.07± 101.9 |
| | Kaempferol-7-neohesperidoside | C27H30O15 | 81.86± 12.44 | 37.62± 9.25 | 16.41± 7.52 | 8.14± 1.32 | 11.04± 1.88 | 5.23± 1.49 | 5.32± 1.3 |
| | Kukoamine B | C28H42N4O6 | 11.24± 3.68 | 9.47± 2.31 | 7.39± 2.59 | 6.55± 1.93 | 5.98± 1.62 | 9.82± 2.68 | 4.47± 1.81 |
| | Luteolin-3',7-di-O-glucoside | C27H30O16 | 11.24± 3.29 | 6.24± 4.66 | 12.12± 7.81 | 5.08± 3.25 | 1.46± 0.79 | 4.46± 1.81 | 1.42± 0.46 |
| | Luteolin-7-O-glucoside | C21H20O11 | 1.18± 0.43 | 0.29± 0.17 | 0.42± 0.29 | 0.29± 0.12 | 0.02± 0.05 | 0.12± 0.04 | 0.17± 0.07 |
| | methoxy-myricetin-O-hexosyl-deoxyhexoside | C28H32O17 | 276.23± 41.73 | 291.7± 48.45 | 522.16± 112.56 | 447.46± 39.82 | 324.35± 52.88 | 465.2± 65.96 | 385.46± 39.14 |
| | methyl chlorogenate | C17H20O9 | 938.54± 185.23 | 356.18± 98.27 | 343.46± 45.96 | 325.01± 70.01 | 266.78± 26.88 | 293.83± 36.44 | 420.31± 90.65 |
| | Methylophiopogonanone A | C19H18O6 | 321.53± 63.06 | 203.65± 22.94 | 200.48± 27.3 | 182.33± 13.02 | 185.72± 16.03 | 164.91± 13.86 | 175.33± 20.09 |
| | N-Caffeoylputrescine | C13H18N2O3 | 4447.58± 1696.11 | 7374.79± 1834.97 | 4007.92± 1124.97 | 2860.76± 861.33 | 5953.71± 2216.23 | 4448.75± 1170.36 | 7311.05± 1969.47 |
| | Panasenoside | C27H30O16 | 74.32± 8.98 | 39.89± 7.89 | 33.12± 4.78 | 77.08± 14.66 | 31.04± 2.02 | 23.81± 5.81 | 42.85± 4.54 |
| | Prunin | C21H22O10 | 12.29± 2.34 | 6.34± 1.86 | 5.04± 1.12 | 8.71± 1.94 | 5.65± 0.6 | 4.01± 0.36 | 5.26± 0.74 |

| | | | | | | | | | |
|---|---|---|---|---|---|---|---|---|---|
| | Quercetin | C15H10O7 | 100.13± 14.88 | 203.02± 40.12 | 232.87± 45.57 | 203.17± 33.35 | 201.42± 35.87 | 171.07± 13.06 | 186.59± 19.51 |
| | Quercetin 3-gentiobioside | C27H30O17 | 708.06± 139.31 | 1292.34± 262.26 | 1813.57± 422.49 | 1552.13± 251.78 | 1455.86± 327.61 | 1313.58± 145.06 | 1429.27± 160.66 |
| | Quercetin-3,4'-O-di-beta-glucopyranoside | C27H30O17 | 328.95± 72.17 | 408.73± 40.08 | 181.39± 20.1 | 214.07± 35.94 | 126.96± 14.5 | 129.36± 12.36 | 100.97± 4.6 |
| | RUTOSIDE (rutin) | C27H30O16 | 162.44± 135.83 | 141.47± 90.36 | 24.03± 4.94 | 20.55± 5.58 | 12.05± 2.65 | 7.05± 1.07 | 29.89± 19.61 |
| | Scoparone | C11H10O4 | 331.79± 196.51 | 33.7± 14.49 | 166.19± 80.21 | 68.12± 46.28 | 36.85± 15.79 | 333.41± 205.59 | 194.78± 89.6 |
| | Scopoletin | C10H8O4 | 45.79± 12.32 | 26.67± 12.12 | 39.53± 14.87 | 25.03± 5.78 | 13.73± 4.82 | 42.45± 9.95 | 47.86± 6.94 |
| | Tiliroside | C30H26O13 | 12.05± 1.79 | 6.13± 1.48 | 2.59± 1.02 | 1.28± 0.2 | 1.91± 0.22 | 0.75± 0.19 | 0.78± 0.18 |
| | trans-5-O-Caffeoylquinic acid | C16H18O9 | 0± 0 | 0± 0 | 0± 0 | 0± 0 | 0± 0 | 0± 0 | 7.86± 6.42 |
| | trans-Caffeic acid | C9H8O4 | 106.07± 11.96 | 70.8± 12.76 | 49.15± 7.57 | 24.94± 6.17 | 29.42± 2.63 | 23.78± 4.64 | 18.85± 3.2 |
| | 8-Gingerol | C19H30O4 | 240.7± 74.35 | 307.03± 69.35 | 171.3± 54.34 | 188.62± 45.63 | 161.69± 58.27 | 218.9± 55.84 | 109.11± 37.42 |
| | Catechin gallate | C22H18O10 | 67.44± 43.69 | 7.13± 7.13 | 26.94± 17.56 | 51.12± 34.78 | 8.61± 8.61 | 25.02± 16.53 | 35.77± 24.66 |
| | Vanillin | C8H8O3 | 23.14± 3.07 | 22.62± 5.56 | 14.92± 2.68 | 14.11± 1.72 | 9.29± 1.7 | 15.51± 2.23 | 10.08± 1.32 |
| Polycyclic Hydrocarbons | hypericin | C30H16O8 | 0.51± 0.21 | 0.19± 0.17 | 0± 0 | 0.03± 0.02 | 0± 0 | 0.02± 0.01 | 0± 0 |
| Pyrimidine nucleosides | Cytidine | C9H13N3O5 | 29.19± 3.64 | 24.29± 3.31 | 25.05± 1.91 | 23.32± 3.5 | 21.4± 2.31 | 21.84± 2.7 | 18.27± 2.91 |

| | | | | | | | | | |
|---|---|---|---|---|---|---|---|---|---|
| Quinilines | 4-Hydroxyquinoline | C9H7NO | 255.71± 18.95 | 197.78± 19.16 | 178.56± 22.87 | 102.28± 13.85 | 114.91± 10.71 | 131.26± 11.84 | 141.9± 7.27 |
| Steroids and derivatives | andrastin A | C28H38O7 | 23.14± 4.71 | 16.18± 3.7 | 4.87± 0.97 | 3.56± 0.56 | 3.43± 0.52 | 3.08± 0.17 | 2.24± 0.38 |
| | Dehydroisoandrosterone sulfate | C19H28O5S | 0± 0 | 1.29± 0.89 | 0.75± 0.62 | 1.42± 0.58 | 0.07± 0.42 | 1.5± 0.73 | 1.96± 0.97 |
| | Edpetiline | C33H53NO8 | 10.27± 5.25 | 2.24± 0.4 | 3.83± 1.16 | 7.15± 2.44 | 2.49± 1.63 | 3.13± 1.3 | 0.88± 0.38 |
| | Eurycomalactone | C19H24O6 | 1.01± 0.25 | 3.08± 1.1 | 2.62± 0.84 | 1.22± 0.33 | 2.54± 1.17 | 1.46± 0.41 | 1.24± 0.35 |
| | Flutamide | C11H11F3N2O3 | 32.06± 2.3 | 23.06± 1.91 | 18.35± 1.3 | 17.36± 1.73 | 9.01± 0.17 | 11.15± 1.08 | 8.46± 0.65 |
| | Hydrocortisone | C21H30O5 | 112.94± 31.74 | 134.94± 38.31 | 93.91± 25.46 | 160.53± 93.95 | 86.68± 25.18 | 145.91± 70.19 | 91.71± 46.49 |
| | Hydroxyprogesterone | C21H30O3 | 614.14± 163.71 | 845.38± 300.62 | 546.15± 95.95 | 836.21± 302.64 | 547.56± 257.67 | 878.33± 355.51 | 644.4± 319.15 |
| | Mestranol | C21H26O2 | 59.73± 9.68 | 44.33± 6.12 | 16.95± 2.41 | 14.02± 2.95 | 8.38± 1.3 | 11.21± 1.49 | 7.27± 1.38 |
| | Progesterone | C21H30O2 | 20.62± 5.31 | 1197.48± 556.46 | 1061.14± 124.51 | 1425.78± 310.9 | 812.72± 318.37 | 1085.33± 319.51 | 1034.32± 285.97 |
| | senegenin | C30H45ClO6 | 2251.66± 1115.74 | 1205.1± 277.99 | 2035.04± 695.11 | 1278.66± 311.53 | 1537.48± 570.31 | 1170.96± 253.81 | 1082.51± 304.38 |
| Terpenes and Terpenoids | 13-alpha-(21)-Epoxyeurycomanone | C20H24O10 | 113.7± 15.3 | 20.12± 3.07 | 40.16± 12.4 | 42.34± 11.25 | 17.98± 2.28 | 32.66± 12.41 | 25.46± 6.41 |
| | 2-(4-oxoquinazolin-3(4H)-yl)ethyl isobutyrate | C14H16N2O3 | 35.42± 6.34 | 32.08± 2.77 | 34.05± 2.72 | 31.51± 3.25 | 28.03± 3.15 | 27.9± 2.83 | 25.99± 3.18 |

| | | | | | | | | | |
|---|---|---|---|---|---|---|---|---|---|
| | 2-(furan-3-yl)-7,8-dihydroxy-6a,7,10b-trimethyl-octahydro-1H-naphtho[2,1-c]pyran-4-one | C20H28O5 | 41.78± 11.88 | 29.5± 7.01 | 25.42± 4.61 | 48± 26.56 | 20.67± 4.87 | 33.44± 14.9 | 20.83± 9.9 |
| | 20-deoxyingenol | C20H28O4 | 1.34± 0.36 | 1.3± 0.3 | 1.2± 0.29 | 1.71± 0.39 | 0.61± 0.06 | 1.1± 0.3 | 1.02± 0.16 |
| | Asiatic Acid | C30H48O5 | 3.84± 2.28 | 1.08± 0.33 | 0.16± 0.11 | 0± 0 | 0.04± 0.04 | 0.19± 0.12 | 0.06± 0.06 |
| | Asperulosidic acid | C18H24O12 | 0.17± 0.02 | 0.34± 0.14 | 0.22± 0.05 | 0.11± 0.03 | 0.18± 0.05 | 0.15± 0.03 | 0.13± 0.04 |
| | bilobalide | C15H18O8 | 21.83± 2.66 | 16.8± 3.58 | 15.63± 2.02 | 12.87± 2.57 | 18.01± 1.34 | 11.61± 2.09 | 14.2± 2.1 |
| | Daphylloside | C21H36O10 | 1.32± 0.91 | 6.43± 3.16 | 2.68± 1.71 | 3.01± 1.25 | 1.48± 0.94 | 3.76± 2.04 | 5.59± 2.29 |
| | gamma,gamma-Dimethyallyl pyrophosphate ammonium salt | C5H12O7P2 | 2.69± 1.16 | 1.35± 0.75 | 0.78± 0.54 | 0.95± 0.28 | 0.43± 0.36 | 0.67± 0.16 | 0.4± 0.13 |
| | Ganoderic acid D2 | C30H42O8 | 1.01± 0.23 | 1.28± 0.34 | 0.64± 0.13 | 0.93± 0.21 | 0.8± 0.12 | 0.61± 0.13 | 0.66± 0.23 |
| | gardenoside | C17H24O11 | 0.06± 0.03 | 0.08± 0.04 | 0.04± 0.04 | 0.05± 0.02 | 0.07± 0.05 | 0.12± 0.02 | 0.06± 0.03 |
| | geniposidic acid | C16H22O10 | 4.12± 1.49 | 1.7± 1.02 | 1.1± 0.36 | 1.32± 0.4 | 0.7± 0.45 | 1.47± 0.55 | 1.39± 0.44 |
| | Ginsenoside Rf | C42H72O14 | 9.45± 5.89 | 1.81± 0.67 | 3.84± 2.39 | 3.35± 1.5 | 4.93± 1.83 | 1.48± 0.73 | 1.55± 0.44 |
| | Ginsenoside Rh1 | C36H62O9 | 2.03± 0.17 | 1.83± 0.26 | 0.86± 0.06 | 0.78± 0.05 | 0.38± 0.08 | 0.62± 0.05 | 0.59± 0.04 |
| | Gossypol | C30H30O8 | 2.31± 0.43 | 4.49± 1.97 | 1.23± 0.33 | 1.95± 0.52 | 0.6± 0.24 | 0.32± 0.02 | 0.53± 0.19 |

| | | | | | | | | | |
|---|---|---|---|---|---|---|---|---|---|
| | Hederagenin base + O-AcetylHex | C38H60O10 | 20.26± 5.39 | 10.7± 0.99 | 4.91± 0.24 | 3.19± 0.3 | 2.38± 0.27 | 3.32± 0.11 | 2.69± 0.15 |
| | Hetisine | C20H27NO3 | 0± 0 | 0± 0 | 0± 0 | 0± 0 | 0± 0 | 0± 0 | 0± 0 |
| | Hydroxyvalerenic Acid | C15H22O3 | 0.67± 0.23 | 0.44± 0.06 | 0.18± 0.08 | 1.27± 1.21 | 0.65± 1.71 | 2.99± 1.82 | 2.23± 1.39 |
| | Lagochilin | C20H36O5 | 3.78± 1.25 | 6.02± 1.29 | 3.36± 1.04 | 4.46± 1.19 | 3.66± 1.57 | 4.29± 1.02 | 2.63± 0.96 |
| | Lucidenic acid D | C29H38O8 | 5.06± 2.56 | 2.37± 1.15 | 3.94± 1.93 | 3.67± 1.55 | 2.33± 1.32 | 2.97± 1.51 | 3.29± 1.33 |
| | Miltirone | C19H22O2 | 33.03± 4.08 | 31.83± 5.25 | 28.48± 3.76 | 25.7± 4.63 | 26.4± 3.43 | 22.06± 2.78 | 23.07± 4.11 |
| | Muurolladie-3-One | C15H22O | 12.89± 2.23 | 40.15± 16.92 | 260.66± 171.73 | 21.24± 5.08 | 20.97± 5.51 | 292.26± 132.59 | 182.5± 98.81 |
| | Ophiopogonoside A | C21H38O8 | 199.19± 31.15 | 227.33± 39.92 | 187.43± 29.2 | 133.67± 26.34 | 179.44± 20.86 | 181.7± 32.8 | 158.02± 19.51 |
| | Quillaic acid | C30H46O5 | 0.41± 0.16 | 0.77± 0.13 | 0.6± 0.21 | 2.17± 0.88 | 1.71± 1.06 | 0.18± 0.1 | 3.56± 1.32 |
| | Sylvestroside I | C33H48O19 | 1.9± 0.54 | 2.18± 0.23 | 1.4± 0.42 | 1.54± 0.3 | 0.45± 0.11 | 0.83± 0.14 | 1.03± 0.11 |
| Vitamins | D-(+)-Pantothenic acid | C9H17NO5 | 183.32± 26.2 | 206.66± 33.82 | 286.2± 38.84 | 177.27± 30.43 | 190.72± 22.2 | 208.96± 21.73 | 159.69± 15.43 |
| | Niacinamide | C6H6N2O | 41.64± 4.65 | 42.58± 2.84 | 39.55± 2.46 | 41.61± 1.49 | 38.01± 2.1 | 33.52± 3.5 | 27.07± 3.94 |
| | Nicotinamide | C6H6N2O | 73.61± 11.93 | 68.04± 5.36 | 48± 9.28 | 68.51± 8.95 | 50.68± 6.79 | 51.45± 7.23 | 38.21± 7.65 |

**Table S3 |** List of identified and annotated metabolites in eggplant leaves and their correlations with larval occurrence, mass, and mortality.

| Class | Compound Name | RT | Adduct / Charge | Molecular formula | Precursor ion mass (Da) | Spearman's ($r_s$) correlation of metabolite concentration among seven eggplant varieties w/ larval | | |
|---|---|---|---|---|---|---|---|---|
| | | | | | | Occurrence | Mass | Mortality |
| Aldehyde and ketones | 1-(4-methoxyphenyl) ethanone | 2.29 | [M+H]+ | C9H10O2 | 151.075 | $R^2$= -0.75 P= 0.04 | $R^2$= -0.75 P= 0.06 | $R^2$= 0.78 P= 0.06 |
| | Dihydrojasmone | 8.38 | [M+H]+ | C11H18O | 167.143 | $R^2$= -0.32 P= 0.44 | $R^2$= -0.32 P= 0.44 | $R^2$= 0.35 P= 0.44 |
| | Phenylacetaldehyde | 1.31 | [M+H]+ | C8H8O | 121.065 | $R^2$= -0.67 P= 0.1 | $R^2$= -0.67 P= 0.08 | $R^2$= 0.64 P= 0.08 |
| | 3-FORMYLINDOLE | 9.13 | [M-H]- | C9H7NO | 144.045 | $R^2$= 0.07 P= 0.83 | $R^2$= 0.07 P= 0.83 | $R^2$= 0 P= 1 |
| | 4-Hydroxybenzaldehyde | 8.28 | [M-H]- | C7H6O2 | 121.03 | $R^2$= 0.17 P= 0.71 | $R^2$= 0.17 P= 0.71 | $R^2$= -0.07 P= 0.83 |
| | Protocatechuic aldehyde | 7.69 | [M-H]- | C7H6O3 | 137.024 | $R^2$= 0.64 P= 0.1 | $R^2$= 0.64 P= 0.1 | $R^2$= -0.57 P= 0.16 |
| Alkaloids and derivatives | Brucine | 7.18 | [M+H]+ | C23H26N2O4 | 395.197 | $R^2$= -0.21 P= 0.44 | $R^2$= -0.21 P= 0.59 | $R^2$= 0.32 P= 0.59 |
| | Calycanthine | 10.33 | [M+H]+ | C22H26N4 | 347.223 | $R^2$= -0.07 P= 0.59 | $R^2$= -0.07 P= 0.83 | $R^2$= 0.21 P= 0.83 |
| | Cinchonine | 0.1 | [M-H]- | C19H22N2O | 293.166 | $R^2$= -0.17 P= 0.71 | $R^2$= -0.17 P= 0.71 | $R^2$= 0.21 P= 0.59 |
| | Hirsuteine | 9.97 | [M-H]- | C22H26N2O3 | 365.187 | $R^2$= 0.07 P= 0.83 | $R^2$= 0.07 P= 0.83 | $R^2$= 0 P= 1 |

| | | | | | | | | |
|---|---|---|---|---|---|---|---|---|
| | HYDROQUINIDINE | 12.16 | [M-H]- | $C_{20}H_{26}N_2O_2$ | 325.192 | $R^2$= -0.6<br>P= 0.13 | $R^2$= -0.6<br>P= 0.13 | $R^2$= 0.53<br>P= 0.23 |
| | Imperialine | 6.53 | [M+H]+ | $C_{27}H_{43}NO_3$ | 430.332 | $R^2$= -0.03<br>P= 0.71 | $R^2$= -0.03<br>P= 0.9 | $R^2$= 0.14<br>P= 0.9 |
| | Khasianine | 6.96 | [M+H]+ | $C_{39}H_{63}NO_{11}$ | 722.447 | $R^2$= 0.12<br>P= 0.85 | $R^2$= 0.12<br>P= 0.8 | $R^2$= -0.09<br>P= 0.8 |
| | quinidine | 7.01 | [M+H]+ | $C_{20}H_{24}N_2O_2$ | 325.191 | $R^2$= -0.05<br>P= 0.79 | $R^2$= -0.05<br>P= 0.91 | $R^2$= 0.12<br>P= 0.91 |
| | Ranaconitine | 8.68 | [M+H]+ | $C_{32}H_{44}N_2O_9$ | 601.312 | $R^2$= -0.14<br>P= 0.59 | $R^2$= -0.14<br>P= 0.71 | $R^2$= 0.21<br>P= 0.71 |
| | Solamargine | 6.96 | [M+H]+ | $C_{45}H_{73}NO_{15}$ | 868.505 | $R^2$= -0.32<br>P= 0.49 | $R^2$= -0.32<br>P= 0.44 | $R^2$= 0.28<br>P= 0.44 |
| | Solanidine base -2H + 1O, O-Hex-dHex-dHex | 6.5 | [M+H]+ | $C_{45}H_{71}NO_{15}$ | 866.49 | $R^2$= -0.21<br>P= 0.49 | $R^2$= -0.21<br>P= 0.59 | $R^2$= 0.28<br>P= 0.59 |
| | Solasodine | 7.27 | [M+H]+ | $C_{27}H_{43}NO_2$ | 414.337 | $R^2$= -0.35<br>P= 0.3 | $R^2$= -0.35<br>P= 0.44 | $R^2$= 0.46<br>P= 0.44 |
| | Solasonine | 6.92 | [M+H]+ | $C_{45}H_{73}NO_{16}$ | 884.5 | $R^2$= 0.64<br>P= 0.13 | $R^2$= 0.64<br>P= 0.1 | $R^2$= -0.6<br>P= 0.1 |
| | Splendoline | 7.42 | [M+H]+ | $C_{21}H_{26}N_2O_4$ | 371.197 | $R^2$= 0.35<br>P= 0.3 | $R^2$= 0.35<br>P= 0.44 | $R^2$= -0.46<br>P= 0.44 |
| | Strictosamide | 10.84 | [M-H]- | $C_{26}H_{30}N_2O_8$ | 497.193 | $R^2$= -0.14<br>P= 0.71 | $R^2$= -0.14<br>P= 0.71 | $R^2$= 0.07<br>P= 0.83 |
| | Subsessiline | 9.81 | [M+H]+ | $C_{43}H_{48}N_4O_6$ | 717.365 | $R^2$= 0.03<br>P= 0.71 | $R^2$= 0.03<br>P= 0.9 | $R^2$= 0.14<br>P= 0.9 |
| | Trigonelline | 1.24 | [M+H]+ | $C_7H_7NO_2$ | 138.055 | $R^2$= -0.35<br>P= 0.44 | $R^2$= -0.35<br>P= 0.44 | $R^2$= 0.32<br>P= 0.44 |
| | Vinpocetine | 6.95 | [M+H]+ | $C_{22}H_{26}N_2O_2$ | 351.207 | $R^2$= -0.53<br>P= 0.1 | $R^2$= -0.53<br>P= 0.23 | $R^2$= 0.64<br>P= 0.23 |

| | | | | | | | | |
|---|---|---|---|---|---|---|---|---|
| | Xanthine | 3.43 | [M+H]+ | C5H4N4O2 | 153.041 | $R^2$= -0.75 P= 0.03 | $R^2$= -0.75 P= 0.06 | $R^2$= 0.82 P= 0.06 |
| Amines | Spermidine | 0.91 | [M+H]+ | C7H19N3 | 146.165 | $R^2$= 0.32 P= 0.49 | $R^2$= 0.32 P= 0.44 | $R^2$= -0.28 P= 0.44 |
| | Spermine | 6.26 | [M+H]+ | C10H26N4 | 203.223 | $R^2$= -0.14 P= 0.78 | $R^2$= -0.14 P= 0.71 | $R^2$= 0.1 P= 0.71 |
| | Tebuconazole | 10.05 | [M+H]+ | C16H22ClN3O | 308.152 | $R^2$= -0.42 P= 0.3 | $R^2$= -0.42 P= 0.3 | $R^2$= 0.46 P= 0.3 |
| | Tyramine | 1.32 | [M+H]+ | C8H11NO | 138.091 | $R^2$= -0.67 P= 0.1 | $R^2$= -0.67 P= 0.08 | $R^2$= 0.64 P= 0.08 |
| Amino acids and derivatives | 1-Aminocyclopropane-1-carboxylate | 1.17 | [M+H]+ | C4H7NO2 | 102.055 | $R^2$= -0.25 P= 0.49 | $R^2$= -0.25 P= 0.59 | $R^2$= 0.28 P= 0.59 |
| | 2-(4-aminotetrahydro-2H-pyran-4-yl) acetic acid | 3.28 | [M+H]+ | C7H13NO3 | 160.097 | $R^2$= -0.67 P= 0.08 | $R^2$= -0.67 P= 0.08 | $R^2$= 0.71 P= 0.08 |
| | Aphyllic Acid | 11.46 | [M-H]- | C15H26N2O2 | 265.192 | $R^2$= -0.25 P= 0.59 | $R^2$= -0.25 P= 0.59 | $R^2$= 0.35 P= 0.44 |
| | Arginine | 1.03 | [M+H]+ | C6H14N4O2 | 175.119 | $R^2$= 0.42 P= 0.49 | $R^2$= 0.42 P= 0.3 | $R^2$= -0.28 P= 0.3 |
| | Asparagine | 1.11 | [M-H]- | C4H8N2O3 | 131.046 | $R^2$= 0.53 P= 0.23 | $R^2$= 0.53 P= 0.23 | $R^2$= -0.6 P= 0.13 |
| | Aspartic acid | 1.16 | [M-H]- | C4H7NO4 | 132.03 | $R^2$= 0.28 P= 0.49 | $R^2$= 0.28 P= 0.49 | $R^2$= -0.32 P= 0.44 |
| | Betaine | 1.2 | [M+H]+ | C5H11NO2 | 118.086 | $R^2$= -0.03 P= 0.83 | $R^2$= -0.03 P= 0.9 | $R^2$= 0.07 P= 0.9 |
| | Citrulline | 1.12 | [M+H]+ | C6H13N3O3 | 176.103 | $R^2$= 0 P= 0.71 | $R^2$= 0 P= 1 | $R^2$= 0.17 P= 1 |

| | | | | | | | | |
|---|---|---|---|---|---|---|---|---|
| | CocamidopropylBetaine | 9.94 | [M+H]+ | C19H38N2O3 | 343.296 | $R^2$= 0.5 P= 0.3 | $R^2$= 0.5 P= 0.26 | $R^2$= -0.42 P= 0.26 |
| | Diphenylamine | 9.64 | [M+H]+ | C12H11N | 170.096 | $R^2$= 0 P= 0.9 | $R^2$= 0 P= 1 | $R^2$= -0.03 P= 1 |
| | DL-3-Aminoisobutyric acid | 1.14 | [M-H]- | C4H9NO2 | 102.056 | $R^2$= 0.21 P= 0.59 | $R^2$= 0.21 P= 0.59 | $R^2$= -0.14 P= 0.71 |
| | feruloyltyramine | 9.47 | [M-H]- | C18H19NO4 | 312.124 | $R^2$= 0.64 P= 0.1 | $R^2$= 0.64 P= 0.1 | $R^2$= -0.6 P= 0.13 |
| | gamma-Glutamyltyrosine | 6.15 | [M+H]+ | C14H18N2O6 | 311.124 | $R^2$= 0.28 P= 0.78 | $R^2$= 0.28 P= 0.49 | $R^2$= -0.1 P= 0.49 |
| | Glutamic acid | 1.18 | [M+H]+ | C5H9NO4 | 148.06 | $R^2$= -0.46 P= 0.23 | $R^2$= -0.46 P= 0.3 | $R^2$= 0.53 P= 0.3 |
| | Glutamine | 1.14 | [M+H]+ | C5H10N2O3 | 147.076 | $R^2$= 0.28 P= 0.44 | $R^2$= 0.28 P= 0.49 | $R^2$= -0.32 P= 0.49 |
| | Glutathione (oxidized) | 3.21 | [M+H]+ | C20H32N6O12S2 | 613.159 | $R^2$= 0.03 P= 0.9 | $R^2$= 0.03 P= 0.9 | $R^2$= 0.03 P= 0.9 |
| | Histamine | 1.01 | [M+H]+ | C5H9N3 | 112.087 | $R^2$= 0.28 P= 0.59 | $R^2$= 0.28 P= 0.49 | $R^2$= -0.21 P= 0.49 |
| | Isoleucine | 1.98 | [M+H]+ | C6H13NO2 | 132.102 | $R^2$= 0.21 P= 0.71 | $R^2$= 0.21 P= 0.59 | $R^2$= -0.17 P= 0.59 |
| | L-5-Oxoproline | 1.14 | [M-H]- | C5H7NO3 | 128.035 | $R^2$= 0.32 P= 0.44 | $R^2$= 0.32 P= 0.44 | $R^2$= -0.28 P= 0.49 |
| | L-beta-Homoisoleucine | 4.46 | [M+H]+ | C7H15NO2 | 146.118 | $R^2$= -0.71 P= 0.03 | $R^2$= -0.71 P= 0.08 | $R^2$= 0.82 P= 0.08 |
| | L-glutamic acid-L-glutamine | 1.47 | [M+H]+ | C10H17N3O6 | 276.119 | $R^2$= 0.1 P= 0.83 | $R^2$= 0.1 P= 0.78 | $R^2$= -0.07 P= 0.78 |

| | | | | | | | | |
|---|---|---|---|---|---|---|---|---|
| | L-Glutathione (oxidized form) | 1.4 | [M-H]- | C20H32N6O12S2 | 611.145 | $R^2$= -0.03 P= 0.9 | $R^2$= -0.03 P= 0.9 | $R^2$= 0 P= 1 |
| | L-Phenylalanine | 4.6 | [M+H]+ | C9H11NO2 | 166.086 | $R^2$= -0.21 P= 0.39 | $R^2$= -0.21 P= 0.59 | $R^2$= 0.39 P= 0.59 |
| | N, N-Dimethylarginine | 1.12 | [M+H]+ | C8H18N4O2 | 203.15 | $R^2$= -0.14 P= 0.59 | $R^2$= -0.14 P= 0.71 | $R^2$= 0.21 P= 0.71 |
| | N-Acetylarginine | 1.21 | [M+H]+ | C8H16N4O3 | 217.13 | $R^2$= 0.28 P= 0.59 | $R^2$= 0.28 P= 0.49 | $R^2$= -0.21 P= 0.49 |
| | N-Acetylhistidine | 1.17 | [M+H]+ | C8H11N3O3 | 198.087 | $R^2$= -0.03 P= 0.83 | $R^2$= -0.03 P= 0.9 | $R^2$= 0.07 P= 0.9 |
| | N-ACETYL-L-LEUCINE | 8.66 | [M-H]- | C8H15NO3 | 172.098 | $R^2$= 0.35 P= 0.44 | $R^2$= 0.35 P= 0.44 | $R^2$= -0.32 P= 0.44 |
| | N-acetyl phenylalanine | 9.06 | [M-H]- | C11H13NO3 | 206.082 | $R^2$= 0.32 P= 0.44 | $R^2$= 0.32 P= 0.44 | $R^2$= -0.28 P= 0.49 |
| | N-acetyltryptophan | 9.14 | [M-H]- | C13H14N2O3 | 245.093 | $R^2$= 0.28 P= 0.49 | $R^2$= 0.28 P= 0.49 | $R^2$= -0.32 P= 0.44 |
| | N-acetyltyramine | 6.37 | [M+H]+ | C10H13NO2 | 180.102 | $R^2$= -0.14 P= 0.59 | $R^2$= -0.14 P= 0.71 | $R^2$= 0.21 P= 0.71 |
| | N-Fructosyl tyrosine | 6.7 | [M+H]+ | C15H21NO8 | 344.134 | $R^2$= 0.28 P= 0.44 | $R^2$= 0.28 P= 0.49 | $R^2$= -0.32 P= 0.49 |
| | O-Phosphothreonine | 1.75 | [M-H]- | C4H10NO6P | 198.017 | $R^2$= -0.03 P= 0.9 | $R^2$= -0.03 P= 0.9 | $R^2$= 0 P= 1 |
| | Pantothenic acid | 6.22 | [M+H]+ | C9H17NO5 | 220.118 | $R^2$= 0.1 P= 1 | $R^2$= 0.1 P= 0.78 | $R^2$= 0 P= 0.78 |
| | Phenylalanine | 4.22 | [M-H]- | C9H11NO2 | 164.072 | $R^2$= 0.32 P= 0.44 | $R^2$= 0.32 P= 0.44 | $R^2$= -0.17 P= 0.71 |
| | Proline | 1.25 | [M+H]+ | C5H9NO2 | 116.071 | $R^2$= -0.42 P= 0.16 | $R^2$= -0.42 P= 0.3 | $R^2$= 0.57 P= 0.3 |

| | | | | | | | | |
|---|---|---|---|---|---|---|---|---|
| | Pyroglutamic acid | 2.77 | [M+H]+ | C5H7NO3 | 130.05 | $R^2$= -0.1<br>P= 0.71 | $R^2$= -0.1<br>P= 0.78 | $R^2$= 0.17<br>P= 0.78 |
| | Quercetin 3-O-malonylglucoside | 8.08 | [M+H]+ | C24H22O15 | 551.103 | $R^2$= 0.07<br>P= 0.9 | $R^2$= 0.07<br>P= 0.83 | $R^2$= -0.03<br>P= 0.83 |
| | Serine | 1.1 | [M-H]- | C3H7NO3 | 104.035 | $R^2$= 0.32<br>P= 0.44 | $R^2$= 0.32<br>P= 0.44 | $R^2$= -0.28<br>P= 0.49 |
| | Threonine | 1.11 | [M-H]- | C4H9NO3 | 118.051 | $R^2$= 0.28<br>P= 0.49 | $R^2$= 0.28<br>P= 0.49 | $R^2$= -0.39<br>P= 0.39 |
| | Triethanolamine | 6.02 | [M+H]+ | C6H15NO3 | 150.112 | $R^2$= -0.42<br>P= 0.44 | $R^2$= -0.42<br>P= 0.3 | $R^2$= 0.35<br>P= 0.3 |
| | Tyrosine | 2.39 | [M+H]+ | C9H11NO3 | 182.081 | $R^2$= -0.21<br>P= 0.44 | $R^2$= -0.21<br>P= 0.59 | $R^2$= 0.35<br>P= 0.59 |
| | Valine | 6.72 | [M-H]- | C5H11NO2 | 116.072 | $R^2$= 0.34<br>P= 0.45 | $R^2$= 0.34<br>P= 0.45 | $R^2$= -0.3<br>P= 0.51 |
| Anthocyans | Cyanidin-3,5-di-O-glucoside | 8.35 | [M+H]+ | C27H31O16 | 612.168 | $R^2$= 0.01<br>P= 0.98 | $R^2$= 0.01<br>P= 0.98 | $R^2$= -0.01<br>P= 0.98 |
| | Cyanidin-3-glucoside | 9.55 | [M+H]+ | C21H21O11 | 450.116 | $R^2$= 0.28<br>P= 0.49 | $R^2$= 0.28<br>P= 0.49 | $R^2$= -0.42<br>P= 0.3 |
| | Cyanidine-3-O-sambubioside | 9.37 | [M+H]+ | C26H29O15 | 582.158 | $R^2$= -0.11<br>P= 0.84 | $R^2$= -0.11<br>P= 0.84 | $R^2$= 0.03<br>P= 0.96 |
| | Malvidin-3-O-glucoside | 10.21 | [M+H]+ | C23H25O12 | 494.142 | $R^2$= -0.5<br>P= 0.25 | $R^2$= -0.5<br>P= 0.25 | $R^2$= 0.41<br>P= 0.35 |
| | Peonidin-3-O-glucoside | 7.81 | [M+H]+ | C22H23O11 | 464.131 | $R^2$= 0.21<br>P= 0.59 | $R^2$= 0.21<br>P= 0.59 | $R^2$= -0.14<br>P= 0.71 |
| Carbohydrates and derivatives | 1-Cinnamoylpyrrolidine | 6.31 | [M+H]+ | C13H15NO | 202.123 | $R^2$= -0.35<br>P= 0.59 | $R^2$= -0.35<br>P= 0.44 | $R^2$= 0.21<br>P= 0.44 |
| | 2-alpha-Mannobiose | 7.4 | [M-H]- | C12H22O11 | 341.109 | $R^2$= -0.39<br>P= 0.39 | $R^2$= -0.39<br>P= 0.39 | $R^2$= 0.42<br>P= 0.3 |

| | | | | | | | | |
|---|---|---|---|---|---|---|---|---|
| | 2'-Deoxyguanosine-5'-diphosphate | 3.04 | [M-H]- | C10H15N5O10P2 | 426.022 | $R^2$= 0.21 P= 0.59 | $R^2$= 0.21 P= 0.59 | $R^2$= -0.25 P= 0.59 |
| | 5'-S-Methylthioadenosine | 6.35 | [M+H]+ | C11H15N5O3S | 298.097 | $R^2$= 0.03 P= 0.83 | $R^2$= 0.03 P= 0.9 | $R^2$= -0.07 P= 0.9 |
| | Adenine | 1.11 | [M-H]- | C5H5N5 | 134.047 | $R^2$= 0.5 P= 0.26 | $R^2$= 0.5 P= 0.26 | $R^2$= -0.42 P= 0.3 |
| | Adenosine | 3.9 | [M+H]+ | C10H13N5O4 | 268.104 | $R^2$= 0.67 P= 0.1 | $R^2$= 0.67 P= 0.08 | $R^2$= -0.64 P= 0.08 |
| | Adenosine 3'5'-cyclic monophosphate | 5.13 | [M-H]- | C10H12N5O6P | 328.045 | $R^2$= 0.14 P= 0.71 | $R^2$= 0.14 P= 0.71 | $R^2$= -0.25 P= 0.59 |
| | Adenosine 3'-monophosphate | 5.92 | [M-H]- | C10H14N5O7P | 346.056 | $R^2$= 0.6 P= 0.13 | $R^2$= 0.6 P= 0.13 | $R^2$= -0.71 P= 0.08 |
| | Adenosine 5'-diphosphate | 2.98 | [M-H]- | C10H15N5O10P2 | 426.022 | $R^2$= 0 P= 1 | $R^2$= 0 P= 1 | $R^2$= -0.07 P= 0.83 |
| | ADENOSINE 5'-DIPHOSPHATE-2 | 2.81 | [M-H]- | C10H15N5O10P2 | 426.022 | $R^2$= -0.32 P= 0.44 | $R^2$= -0.32 P= 0.44 | $R^2$= 0.39 P= 0.39 |
| | Adenosine_Diphosphate | 4.15 | [M+H]+ | C10H15N5O10P2 | 428.037 | $R^2$= 0.14 P= 0.59 | $R^2$= 0.14 P= 0.71 | $R^2$= -0.21 P= 0.71 |
| | ADP | 3.52 | [M-H]- | C10H15N5O10P2 | 426.022 | $R^2$= 0.1 P= 0.78 | $R^2$= 0.1 P= 0.78 | $R^2$= -0.28 P= 0.49 |
| | AMP | 5.66 | [M-H]- | C10H14N5O7P | 346.056 | $R^2$= 0.64 P= 0.1 | $R^2$= 0.64 P= 0.1 | $R^2$= -0.57 P= 0.16 |
| | beta-D-glucopyranosiduronic acid | 7.58 | [M+H]+ | C21H26N2O4 | 371.197 | $R^2$= 0.14 P= 0.49 | $R^2$= 0.14 P= 0.71 | $R^2$= -0.28 P= 0.71 |
| | beta-Guanidinopropionic acid | 6.2 | [M+H]+ | C4H9N3O2 | 132.077 | $R^2$= -0.07 P= 0.59 | $R^2$= -0.07 P= 0.83 | $R^2$= 0.21 P= 0.83 |
| | beta-Nicotinamide adenine dinucleotide | 3.22 | [M+H]+ | C21H27N7O14P2 | 664.116 | $R^2$= -0.64 P= 0.13 | $R^2$= -0.64 P= 0.1 | $R^2$= 0.6 P= 0.1 |

| | | | | | | | | |
|---|---|---|---|---|---|---|---|---|
| | D-Arabinose-5-phosphate disodium salt | 1.97 | [M-H]- | C5H11O8P | 229.012 | $R^2$= 0.6 P= 0.13 | $R^2$= 0.6 P= 0.13 | $R^2$= -0.64 P= 0.1 |
| | Glucose-6-phosphate | 2.01 | [M+H]+ | C6H13O9P | 261.037 | $R^2$= -0.25 P= 0.71 | $R^2$= -0.25 P= 0.59 | $R^2$= 0.14 P= 0.59 |
| | Guanine | 5.03 | [M+H]+ | C5H5N5O | 152.057 | $R^2$= -0.14 P= 0.59 | $R^2$= -0.14 P= 0.71 | $R^2$= 0.25 P= 0.71 |
| | Guanosine | 5.03 | [M+H]+ | C10H13N5O5 | 284.099 | $R^2$= -0.14 P= 0.59 | $R^2$= -0.14 P= 0.71 | $R^2$= 0.25 P= 0.71 |
| | Guanosine 5'-diphosphate-D-mannose | 5.56 | [M-H]- | C16H25N5O16P2 | 604.07 | $R^2$= -0.42 P= 0.3 | $R^2$= -0.42 P= 0.3 | $R^2$= 0.39 P= 0.39 |
| | Hypoxanthine | 5.18 | [M+H]+ | C5H4N4O | 137.046 | $R^2$= 0 P= 0.9 | $R^2$= 0 P= 1 | $R^2$= 0.03 P= 1 |
| | Inosine | 4.91 | [M+H]+ | C10H12N4O5 | 269.088 | $R^2$= -0.28 P= 0.44 | $R^2$= -0.28 P= 0.49 | $R^2$= 0.32 P= 0.49 |
| | isomaltulose | 1.42 | [M+H]+ | C12H22O11 | 343.123 | $R^2$= -0.75 P= 0.04 | $R^2$= -0.75 P= 0.06 | $R^2$= 0.78 P= 0.06 |
| | Mannose 1-phosphate | 2.01 | [M-H]- | C6H13O9P | 259.022 | $R^2$= 0.53 P= 0.23 | $R^2$= 0.53 P= 0.23 | $R^2$= -0.6 P= 0.13 |
| | Melezitose | 1.33 | [M+H]+ | C18H32O16 | 505.176 | $R^2$= -0.17 P= 0.83 | $R^2$= -0.17 P= 0.71 | $R^2$= 0.07 P= 0.71 |
| | N2-Methylguanosine | 5.99 | [M+H]+ | C11H15N5O5 | 298.115 | $R^2$= -0.32 P= 0.3 | $R^2$= -0.32 P= 0.44 | $R^2$= 0.46 P= 0.44 |
| | roseoside | 6.91 | [M+H]+ | C19H30O8 | 387.201 | $R^2$= -0.07 P= 0.78 | $R^2$= -0.07 P= 0.83 | $R^2$= -0.1 P= 0.83 |
| | Sorbitol | 1.18 | [M-H]- | C6H14O6 | 181.072 | $R^2$= 0.32 P= 0.44 | $R^2$= 0.32 P= 0.44 | $R^2$= -0.28 P= 0.49 |
| | Swertiamarin | 9.88 | [M-H]- | C16H22O10 | 373.114 | $R^2$= -0.4 P= 0.42 | $R^2$= -0.4 P= 0.42 | $R^2$= 0.4 P= 0.42 |

| | | | | | | | | |
|---|---|---|---|---|---|---|---|---|
| | trans-piceid | 7.51 | [M+FA-H]- | C20H22O8 | 435.13 | $R^2$= 0.5 P= 0.26 | $R^2$= 0.5 P= 0.26 | $R^2$= -0.39 P= 0.39 |
| | Trehalose | 1.22 | [M-H]- | C12H22O11 | 341.109 | $R^2$= 0.28 P= 0.49 | $R^2$= 0.28 P= 0.49 | $R^2$= -0.32 P= 0.44 |
| | UDP-D-glucose | 6.84 | [M-H]- | C15H24N2O17P2 | 565.048 | $R^2$= -0.46 P= 0.3 | $R^2$= -0.46 P= 0.3 | $R^2$= 0.57 P= 0.16 |
| | Uracil | 3.58 | [M+H]+ | C4H4N2O2 | 113.035 | $R^2$= -0.03 P= 1 | $R^2$= -0.03 P= 0.9 | $R^2$= 0 P= 0.9 |
| | Uridine | 1.77 | [M-H]- | C9H12N2O6 | 243.062 | $R^2$= 0.64 P= 0.1 | $R^2$= 0.64 P= 0.1 | $R^2$= -0.6 P= 0.13 |
| | Uridine 5'-diphospho-D-glucose | 6.67 | [M-H]- | C15H24N2O17P2 | 565.048 | $R^2$= -0.25 P= 0.59 | $R^2$= -0.25 P= 0.59 | $R^2$= 0.39 P= 0.39 |
| | Xanthosine | 5.91 | [M-H]- | C10H12N4O6 | 283.068 | $R^2$= 0.07 P= 0.83 | $R^2$= 0.07 P= 0.83 | $R^2$= -0.1 P= 0.78 |
| Carboxylic acids and derivatives | 2,8-Quinolinediol | 9.56 | [M+H]+ | C9H7NO2 | 162.055 | $R^2$= -0.6 P= 0.08 | $R^2$= -0.6 P= 0.13 | $R^2$= 0.67 P= 0.13 |
| | 2-acetoxy-4-pentadecylbenzoic acid | 11.36 | [M+H]+ | C24H38O4 | 391.284 | $R^2$= -0.32 P= 0.49 | $R^2$= -0.32 P= 0.44 | $R^2$= 0.28 P= 0.44 |
| | 2-Hydroxy-4-methylpentanoic acid | 8.56 | [M-H]- | C6H12O3 | 131.071 | $R^2$= -0.17 P= 0.71 | $R^2$= -0.17 P= 0.71 | $R^2$= 0.21 P= 0.59 |
| | 2-Isopropylmalic acid | 7.95 | [M-H]- | C7H12O5 | 175.061 | $R^2$= 0.64 P= 0.1 | $R^2$= 0.64 P= 0.1 | $R^2$= -0.57 P= 0.16 |
| | 2-Oxobutyric acid | 6.17 | [M-H]- | C4H6O3 | 101.024 | $R^2$= 0 P= 1 | $R^2$= 0 P= 1 | $R^2$= 0.07 P= 0.83 |
| | 3-(4-HYDROXY-3,5-DIMETHOXYPHENYL)-2-PROPENOIC ACID | 6.88 | [M+H]+ | C11H12O5 | 225.076 | $R^2$= 0.21 P= 0.83 | $R^2$= 0.21 P= 0.59 | $R^2$= -0.07 P= 0.59 |

| | | | | | | | | |
|---|---|---|---|---|---|---|---|---|
| | 3-(4-HYDROXYPHENYL)PROP-2-ENOIC ACID | 6.7 | [M+H]+ | C9H8O3 | 165.055 | $R^2$= 0.03 P= 0.83 | $R^2$= 0.03 P= 0.9 | $R^2$= -0.07 P= 0.9 |
| | 3-(Benzoyloxy)-2-hydroxypropyl beta-D-glucopyranosiduronic acid | 7.41 | [M+H]+ | C16H20O10 | 373.113 | $R^2$= -0.75 P= 0.03 | $R^2$= -0.75 P= 0.06 | $R^2$= 0.82 P= 0.06 |
| | 3-Indolepropionic acid | 7.27 | [M+H]+ | C11H11NO2 | 190.086 | $R^2$= -0.42 P= 0.16 | $R^2$= -0.42 P= 0.3 | $R^2$= 0.57 P= 0.3 |
| | 3-phenyl lactic acid | 9.93 | [M-H]- | C9H10O3 | 165.056 | $R^2$= 0 P= 1 | $R^2$= 0 P= 1 | $R^2$= -0.14 P= 0.71 |
| | 4-Hydroxyquinoline-2-carboxylic acid | 10.31 | [M-H]- | C10H7NO3 | 188.035 | $R^2$= 0.07 P= 0.83 | $R^2$= 0.07 P= 0.83 | $R^2$= -0.1 P= 0.78 |
| | Benzoic acid | 5.37 | [M+H]+ | C7H6O2 | 123.044 | $R^2$= -0.85 P= 0.48 | $R^2$= -0.85 P= 0.12 | $R^2$= 0.78 P= 0.12 |
| | cis-Aconitate | 2.13 | [M-H]- | C6H6O6 | 173.009 | $R^2$= 0.5 P= 0.26 | $R^2$= 0.5 P= 0.26 | $R^2$= -0.42 P= 0.3 |
| | Citric acid | 2.35 | [M-H]- | C6H8O7 | 191.02 | $R^2$= 0.53 P= 0.23 | $R^2$= 0.53 P= 0.23 | $R^2$= -0.46 P= 0.3 |
| | Diethyl phthalate | 8.69 | [M+H]+ | C12H14O4 | 223.096 | $R^2$= -0.07 P= 0.78 | $R^2$= -0.07 P= 0.83 | $R^2$= 0.1 P= 0.83 |
| | Dioctyl Phthalate | 11.31 | [M+H]+ | C24H38O4 | 391.284 | $R^2$= 0 P= 0.9 | $R^2$= 0 P= 1 | $R^2$= -0.03 P= 1 |
| | Ethylenediaminetetraacetic acid EDTA | 1.37 | [M+H]+ | C10H16N2O8 | 293.098 | $R^2$= -0.07 P= 0.71 | $R^2$= -0.07 P= 0.83 | $R^2$= 0.14 P= 0.83 |
| | Fumaric acid | 1.56 | [M-H]- | C4H4O4 | 115.004 | $R^2$= 0.32 P= 0.44 | $R^2$= 0.32 P= 0.44 | $R^2$= -0.28 P= 0.49 |
| | Kynurenic acid | 10.21 | [M-H]- | C10H7NO3 | 188.035 | $R^2$= 0.07 P= 0.83 | $R^2$= 0.07 P= 0.83 | $R^2$= -0.1 P= 0.78 |
| | L-Glutamic acid | 1.14 | [M-H]- | C5H9NO4 | 146.046 | $R^2$= 0.32 P= 0.44 | $R^2$= 0.32 P= 0.44 | $R^2$= -0.28 P= 0.49 |

| | | | | | | | | |
|---|---|---|---|---|---|---|---|---|
| | L-kynurenine | 4.06 | [M+H]+ | C10H12N2O3 | 209.092 | $R^2$= 0.21 P= 0.71 | $R^2$= 0.21 P= 0.59 | $R^2$= -0.14 P= 0.59 |
| | L-Saccharopine | 1.17 | [M+H]+ | C11H20N2O6 | 277.139 | $R^2$= -0.71 P= 0.03 | $R^2$= -0.71 P= 0.08 | $R^2$= 0.82 P= 0.08 |
| | Methyl nicotinic acid | 6.73 | [M+H]+ | C7H7NO2 | 138.055 | $R^2$= 0.25 P= 0.59 | $R^2$= 0.25 P= 0.59 | $R^2$= -0.21 P= 0.59 |
| | MUCIC ACID | 1.79 | [M-H]- | C6H10O8 | 209.03 | $R^2$= 0.5 P= 0.26 | $R^2$= 0.5 P= 0.26 | $R^2$= -0.42 P= 0.3 |
| | Nicotinic acid | 1.83 | [M+H]+ | C6H5NO2 | 124.039 | $R^2$= -0.03 P= 0.71 | $R^2$= -0.03 P= 0.9 | $R^2$= 0.14 P= 0.9 |
| | Phthalic anhydride | 10.4 | [M+H]+ | C8H4O3 | 149.023 | $R^2$= -0.46 P= 0.44 | $R^2$= -0.46 P= 0.3 | $R^2$= 0.35 P= 0.3 |
| | Quercetin-4'-glucoside | 7.45 | [M+H]+ | C21H20O12 | 465.103 | $R^2$= 0.32 P= 0.3 | $R^2$= 0.32 P= 0.44 | $R^2$= -0.42 P= 0.44 |
| | Salicylic acid | 7.58 | [M-H]- | C7H6O3 | 137.024 | $R^2$= 0.53 P= 0.23 | $R^2$= 0.53 P= 0.23 | $R^2$= -0.46 P= 0.3 |
| | Trifloxystrobin | 10.39 | [M+H]+ | C20H19F3N2O4 | 409.137 | $R^2$= -0.14 P= 0.83 | $R^2$= -0.14 P= 0.71 | $R^2$= 0.07 P= 0.71 |
| | Tryptophan | 6.71 | [M-H]- | C11H12N2O2 | 203.083 | $R^2$= 0.17 P= 0.71 | $R^2$= 0.17 P= 0.71 | $R^2$= -0.07 P= 0.83 |
| | Vanillic acid | 6.96 | [M-H]- | C8H8O4 | 167.035 | $R^2$= 0.21 P= 0.59 | $R^2$= 0.21 P= 0.59 | $R^2$= -0.14 P= 0.71 |
| | Xanthurenic Acid | 7.96 | [M+H]+ | C10H7NO4 | 206.045 | $R^2$= -0.67 P= 0.16 | $R^2$= -0.67 P= 0.08 | $R^2$= 0.57 P= 0.08 |
| Cholines | Acetylcholine | 1.12 | [M+H]+ | C7H16NO2 | 147.125 | $R^2$= 0.1 P= 0.71 | $R^2$= 0.1 P= 0.78 | $R^2$= -0.14 P= 0.78 |
| Fatty acid and derivatives | 9,12,15-octadecatrienoic acid | 11.7 | [M+H]+ | C18H30O2 | 279.232 | $R^2$= 0.17 P= 0.59 | $R^2$= 0.17 P= 0.71 | $R^2$= -0.21 P= 0.71 |

| | | | | | | | | |
|---|---|---|---|---|---|---|---|---|
| | Acaranoic acid | 12.51 | [M-H]- | C17H30O4 | 297.207 | $R^2$= -0.53 P= 0.28 | $R^2$= -0.53 P= 0.28 | $R^2$= 0.53 P= 0.28 |
| | Avocadyne Acetate | 10.56 | [M+H]+ | C19H34O4 | 327.253 | $R^2$= 0.03 P= 1 | $R^2$= 0.03 P= 0.9 | $R^2$= 0 P= 0.9 |
| | Azelaic acid | 9.52 | [M-H]- | C9H16O4 | 187.098 | $R^2$= 0.42 P= 0.3 | $R^2$= 0.42 P= 0.3 | $R^2$= -0.32 P= 0.44 |
| | beta-D-Glucopyranoside | 8.48 | [M+H]+ | C21H36O10 | 449.238 | $R^2$= -0.17 P= 0.44 | $R^2$= -0.17 P= 0.71 | $R^2$= 0.32 P= 0.71 |
| | Heptadecanoic acid | 13.9 | [M-H]- | C17H34O2 | 269.249 | $R^2$= 0.5 P= 0.26 | $R^2$= 0.5 P= 0.26 | $R^2$= -0.42 P= 0.3 |
| | hexadecanedioic acid | 10.79 | [M+H]+ | C16H30O4 | 287.222 | $R^2$= 0.5 P= 0.3 | $R^2$= 0.5 P= 0.26 | $R^2$= -0.42 P= 0.26 |
| | Linoelaidic acid | 12.82 | [M-H]- | C18H32O2 | 279.233 | $R^2$= 0.32 P= 0.44 | $R^2$= 0.32 P= 0.44 | $R^2$= -0.28 P= 0.49 |
| | Linoleic acid | 13.43 | [M-H]- | C18H32O2 | 279.233 | $R^2$= 0.17 P= 0.71 | $R^2$= 0.17 P= 0.71 | $R^2$= -0.1 P= 0.78 |
| | linolenic acid | 11.64 | [M+H]+ | C18H30O2 | 279.232 | $R^2$= -0.6 P= 0.23 | $R^2$= -0.6 P= 0.13 | $R^2$= 0.53 P= 0.13 |
| | Mesaconic acid | 2.3 | [M-H]- | C5H6O4 | 129.019 | $R^2$= 0.32 P= 0.44 | $R^2$= 0.32 P= 0.44 | $R^2$= -0.28 P= 0.49 |
| | methyl palmitate | 10.88 | [M+H]+ | C17H34O2 | 271.263 | $R^2$= 0 P= 0.9 | $R^2$= 0 P= 1 | $R^2$= 0.03 P= 1 |
| | Palmitic acid (NMR) | 13.72 | [M-H]- | C16H32O2 | 255.233 | $R^2$= 0.64 P= 0.1 | $R^2$= 0.64 P= 0.1 | $R^2$= -0.6 P= 0.13 |
| | Palmitoleic acid | 13.3 | [M-H]- | C16H30O2 | 253.217 | $R^2$= 0.32 P= 0.44 | $R^2$= 0.32 P= 0.44 | $R^2$= -0.28 P= 0.49 |
| | Stearic acid | 14.36 | [M-H]- | C18H36O2 | 283.264 | $R^2$= 0.5 P= 0.26 | $R^2$= 0.5 P= 0.26 | $R^2$= -0.42 P= 0.3 |

| | | | | | | | | |
|---|---|---|---|---|---|---|---|---|
| | Trans-Vaccenic acid | 13.81 | [M-H]- | C18H34O2 | 281.249 | $R^2$= 0.42 P= 0.3 | $R^2$= 0.42 P= 0.3 | $R^2$= -0.5 P= 0.26 |
| | y-Linolenic acid | 13.14 | [M-H]- | C18H30O2 | 277.217 | $R^2$= -0.1 P= 0.78 | $R^2$= -0.1 P= 0.78 | $R^2$= 0.21 P= 0.59 |
| Flavins | (-)-RIBOFLAVIN | 6.9 | [M+H]+ | C17H20N4O6 | 377.146 | $R^2$= 0.35 P= 0.44 | $R^2$= 0.35 P= 0.44 | $R^2$= -0.32 P= 0.44 |
| Furanochromones | 5-O-methylvisammioside | 7.46 | [M+H]+ | C22H28O10 | 453.176 | $R^2$= -0.6 P= 0.16 | $R^2$= -0.6 P= 0.13 | $R^2$= 0.57 P= 0.13 |
| Indoles | Indole-3-carbinol | 7.94 | [M+H]+ | C9H9NO | 148.076 | $R^2$= 0.12 P= 0.85 | $R^2$= 0.12 P= 0.8 | $R^2$= -0.09 P= 0.8 |
| Indolizidines | Corynoxine | 6.91 | [M+H]+ | C22H28N2O4 | 385.212 | $R^2$= 0.1 P= 0.49 | $R^2$= 0.1 P= 0.78 | $R^2$= -0.28 P= 0.78 |
| Lignans and derivatives | Arctigenin | 8.46 | [M-H]- | C21H24O6 | 371.15 | $R^2$= 0.35 P= 0.44 | $R^2$= 0.35 P= 0.44 | $R^2$= -0.32 P= 0.44 |
| | Eleutheroside E | 8.32 | [M+FA-H]- | C34H46O18 | 787.267 | $R^2$= 0.1 P= 0.78 | $R^2$= 0.1 P= 0.78 | $R^2$= -0.07 P= 0.83 |
| | Secoisolariciresinol | 9.76 | [M-H]- | C20H26O6 | 361.166 | $R^2$= -0.01 P= 0.98 | $R^2$= -0.01 P= 0.98 | $R^2$= 0.1 P= 0.81 |
| Lipids | Dehydrophytosphingosine | 8.4 | [M+H]+ | C18H37NO3 | 316.285 | $R^2$= -0.53 P= 0.08 | $R^2$= -0.53 P= 0.23 | $R^2$= 0.67 P= 0.23 |
| | Phosphocholine | 1.16 | [M+H]+ | C5H15NO4P | 185.081 | $R^2$= -0.75 P= 0.03 | $R^2$= -0.75 P= 0.06 | $R^2$= 0.82 P= 0.06 |
| | quercetin-3-O-glc-1-3-rham-1-6-glucoside | 6.92 | [M+H]+ | C33H40O21 | 773.213 | $R^2$= 0.35 P= 0.44 | $R^2$= 0.35 P= 0.44 | $R^2$= -0.32 P= 0.44 |
| | sn-Glycero-3-phosphocholine | 1.18 | [M+H]+ | C8H21NO6P | 259.118 | $R^2$= -0.14 P= 0.59 | $R^2$= -0.14 P= 0.71 | $R^2$= 0.25 P= 0.71 |
| Macrolides and derivatives | curvularin | 11.97 | [M-H]- | C16H20O5 | 291.124 | $R^2$= -0.4 P= 0.42 | $R^2$= -0.4 P= 0.42 | $R^2$= 0.4 P= 0.42 |

| | | | | | | | | |
|---|---|---|---|---|---|---|---|---|
| | dihydroalbocycline | 9.81 | [M+H]+ | C18H30O4 | 311.222 | $R^2$= 0.07 P= 0.9 | $R^2$= 0.07 P= 0.83 | $R^2$= 0.03 P= 0.83 |
| | Gardnutine | 6.83 | [M+FA-H]- | C20H22N2O2 | 367.166 | $R^2$= 0.28 P= 0.49 | $R^2$= 0.28 P= 0.49 | $R^2$= -0.21 P= 0.59 |
| Oxanes | Cochlioquinone A | 11.91 | [M-H]- | C30H44O8 | 531.296 | $R^2$= -0.17 P= 0.71 | $R^2$= -0.17 P= 0.71 | $R^2$= 0.21 P= 0.59 |
| Phenolics and flavonoids | 1-O-beta-D-Glucopyranosyl sinapate | 7.97 | [M-H]- | C17H22O10 | 385.114 | $R^2$= 0.07 P= 0.83 | $R^2$= 0.07 P= 0.83 | $R^2$= 0 P= 1 |
| | 2-Glucosyloxy-4-methoxy cinnamic acid | 7.36 | [M+H]+ | C16H20O9 | 357.118 | $R^2$= 0.03 P= 0.71 | $R^2$= 0.03 P= 0.9 | $R^2$= -0.14 P= 0.9 |
| | 3,4-di-O-caffeoylquinic acid | 7.51 | [M+H]+ | C25H24O12 | 517.134 | $R^2$= 0.35 P= 0.49 | $R^2$= 0.35 P= 0.44 | $R^2$= -0.28 P= 0.44 |
| | 3-Deoxycaryoptinol | 9.22 | [M-H]- | C24H34O7 | 433.223 | $R^2$= 0.21 P= 0.59 | $R^2$= 0.21 P= 0.59 | $R^2$= -0.14 P= 0.71 |
| | 3-Hydroxy-4-methoxy cinnamic acid (isoferulic acid) | 9.38 | [M-H]- | C10H10O4 | 193.051 | $R^2$= 0.42 P= 0.3 | $R^2$= 0.42 P= 0.3 | $R^2$= -0.28 P= 0.49 |
| | 3-Hydroxycinnamic acid | 7.85 | [M-H]- | C9H8O3 | 163.04 | $R^2$= 0.67 P= 0.08 | $R^2$= 0.67 P= 0.08 | $R^2$= -0.64 P= 0.1 |
| | 3-O-Feruloylquinic acid | 8.62 | [M-H]- | C17H20O9 | 367.103 | $R^2$= -0.03 P= 0.9 | $R^2$= -0.03 P= 0.9 | $R^2$= 0 P= 1 |
| | 4',5,7-Trihydroxy-6,8-diprenylisoflavone | 7.49 | [M+H]+ | C25H26O5 | 407.185 | $R^2$= -0.53 P= 0.13 | $R^2$= -0.53 P= 0.23 | $R^2$= 0.6 P= 0.23 |
| | 4-Caffeoylquinic acid | 8.05 | [M-H]- | C16H18O9 | 353.088 | $R^2$= 0.53 P= 0.23 | $R^2$= 0.53 P= 0.23 | $R^2$= -0.5 P= 0.26 |
| | 6,7-Dihydroxycoumarin | 8.21 | [M-H]- | C9H6O4 | 177.019 | $R^2$= 0.32 P= 0.44 | $R^2$= 0.32 P= 0.44 | $R^2$= -0.35 P= 0.44 |
| | Apigenin-7-O-glucoside | 9.04 | [M-H]- | C21H20O10 | 431.098 | $R^2$= 0 P= 1 | $R^2$= 0 P= 1 | $R^2$= 0.03 P= 0.9 |

| | | | | | | | | |
|---|---|---|---|---|---|---|---|---|
| | Benzoic acid + 1O, 2MeO, O-Hex | 7.37 | [M-H]- | C15H20O10 | 359.098 | $R^2$= 0.46 P= 0.3 | $R^2$= 0.46 P= 0.3 | $R^2$= -0.35 P= 0.44 |
| | Benzoic acid + 2O, O-Hex | 7.3 | [M-H]- | C13H16O9 | 315.072 | $R^2$= 0.32 P= 0.44 | $R^2$= 0.32 P= 0.44 | $R^2$= -0.35 P= 0.44 |
| | Bergenin | 8.46 | [M-H]- | C14H16O9 | 327.072 | $R^2$= 0 P= 1 | $R^2$= 0 P= 1 | $R^2$= -0.03 P= 0.9 |
| | Biochanin-7-O-glucoside | 8.25 | [M+FA-H]- | C22H22O10 | 491.119 | $R^2$= 0.03 P= 0.9 | $R^2$= 0.03 P= 0.9 | $R^2$= 0 P= 1 |
| | Caffeoyl putrescin | 5.69 | [M-H]- | C13H18N2O3 | 249.124 | $R^2$= -0.78 P= 0.04 | $R^2$= -0.78 P= 0.04 | $R^2$= 0.71 P= 0.08 |
| | Caffeic acid hexoside | 7.83 | [M-H]- | C15H18O9 | 341.088 | $R^2$= 0.67 P= 0.08 | $R^2$= 0.67 P= 0.08 | $R^2$= -0.64 P= 0.1 |
| | CAFFEIC ACID | 8.21 | [M-H]- | C9H8O4 | 179.035 | $R^2$= 0.67 P= 0.08 | $R^2$= 0.67 P= 0.08 | $R^2$= -0.64 P= 0.1 |
| | 3-*O*-Caffeoyl quinic acid | 7.41 | [M-H]- | C16H18O9 | 353.088 | $R^2$= -0.21 P= 0.59 | $R^2$= -0.21 P= 0.59 | $R^2$= 0.28 P= 0.49 |
| | Caffeoylcholine | 5.25 | [M+H]+ | C14H20NO4 | 267.147 | $R^2$= 0.53 P= 0.3 | $R^2$= 0.53 P= 0.23 | $R^2$= -0.46 P= 0.23 |
| | **CHLOROGENIC ACID** | **8.14** | **[M-H]-** | **C16H18O9** | **353.088** | **$R^2$= -0.88 P= 0.008** | **$R^2$= -0.82 P= 0.03** | **$R^2$= -0.78 P= 0.01** |
| | Citrusin | 7.62 | [M+H]+ | C16H22O7 | 327.144 | $R^2$= 0.6 P= 0.06 | $R^2$= 0.6 P= 0.13 | $R^2$= -0.75 P= 0.13 |
| | Coumarin + 1O + 1MeO, O-Hex-Hex | 7.76 | [M-H]- | C22H28O14 | 515.141 | $R^2$= -0.14 P= 0.71 | $R^2$= -0.14 P= 0.71 | $R^2$= 0.21 P= 0.59 |
| | Coumaroyl Hexoside | 7.85 | [M-H]- | C15H18O8 | 325.093 | $R^2$= 0.64 P= 0.1 | $R^2$= 0.64 P= 0.1 | $R^2$= -0.57 P= 0.16 |
| | Coumaroyl quinic acid | 8.55 | [M-H]- | C16H18O8 | 337.093 | $R^2$= 0.64 P= 0.1 | $R^2$= 0.64 P= 0.1 | $R^2$= -0.6 P= 0.13 |

| | | | | | | | | |
|---|---|---|---|---|---|---|---|---|
| | Coumaroyl tyramine | 7.63 | [M+H]+ | C17H17NO3 | 284.128 | $R^2$= 0.5 P= 0.23 | $R^2$= 0.5 P= 0.26 | $R^2$= -0.53 P= 0.26 |
| | Cupressuflavone | 7.46 | [M-H]- | C30H18O10 | 537.083 | $R^2$= 0.14 P= 0.71 | $R^2$= 0.14 P= 0.71 | $R^2$= -0.35 P= 0.44 |
| | Cyanidin | 8 | [M+H]+ | C15H11O6 | 288.063 | $R^2$= -0.28 P= 0.53 | $R^2$= -0.28 P= 0.53 | $R^2$= 0.25 P= 0.59 |
| | Cyanidin 3-(2G-glucosylrutinoside) | 7.06 | [M+H]+ | C33H41O20 | 758.226 | $R^2$= 0.03 P= 1 | $R^2$= 0.03 P= 0.9 | $R^2$= 0 P= 0.9 |
| | Cyanidin-3-O-(2"-O-beta-xylopyranosyl-beta-glucopyranoside) | 7.48 | [M+H]+ | C26H29O15 | 582.158 | $R^2$= -0.46 P= 0.49 | $R^2$= -0.46 P= 0.3 | $R^2$= 0.28 P= 0.3 |
| | Cyclohexanecarboxylic acid | 7.07 | [M+H]+ | C16H18O8 | 339.107 | $R^2$= 0 P= 0.83 | $R^2$= 0 P= 1 | $R^2$= -0.07 P= 1 |
| | Cynarin | 7.25 | [M-H]- | C25H24O12 | 515.119 | $R^2$= 0.6 P= 0.13 | $R^2$= 0.6 P= 0.13 | $R^2$= -0.53 P= 0.23 |
| | D-(-)-Quinic acid | 1.39 | [M-H]- | C7H12O6 | 191.056 | $R^2$= 0.67 P= 0.08 | $R^2$= 0.67 P= 0.08 | $R^2$= -0.75 P= 0.06 |
| | Daphnetin | 7.48 | [M-H]- | C9H6O4 | 177.019 | $R^2$= 0.57 P= 0.16 | $R^2$= 0.57 P= 0.16 | $R^2$= -0.5 P= 0.26 |
| | Di-dihydro caffeoyl spermidine | 6.21 | [M+H]+ | C25H35N3O6 | 474.26 | $R^2$= -0.39 P= 0.44 | $R^2$= -0.39 P= 0.39 | $R^2$= 0.35 P= 0.39 |
| | Epigallocatechin | 3.89 | [M+H]+ | C15H14O7 | 307.081 | $R^2$= -0.14 P= 0.59 | $R^2$= -0.14 P= 0.71 | $R^2$= 0.21 P= 0.71 |
| | epsilon-Viniferin | 6.65 | [M+H]+ | C28H22O6 | 455.149 | $R^2$= 0.42 P= 0.23 | $R^2$= 0.42 P= 0.3 | $R^2$= -0.53 P= 0.3 |
| | Eriodictyol-7-O-neohesperidoside | 7.95 | [M-H]- | C27H32O15 | 595.167 | $R^2$= 0.21 P= 0.59 | $R^2$= 0.21 P= 0.59 | $R^2$= -0.14 P= 0.71 |
| | Eriodictyol-7-O-rutinoside | 6.66 | [M+H]+ | C27H32O15 | 597.181 | $R^2$= 0.03 P= 0.83 | $R^2$= 0.03 P= 0.9 | $R^2$= -0.07 P= 0.9 |

| | | | | | | | | |
|---|---|---|---|---|---|---|---|---|
| | Esculin | 6.54 | [M+H]+ | C15H16O9 | 341.087 | $R^2$= -0.1 P= 0.83 | $R^2$= -0.1 P= 0.78 | $R^2$= 0.07 P= 0.78 |
| | Eupatilin | 7.75 | [M-H]- | C18H16O7 | 343.082 | $R^2$= 0.64 P= 0.1 | $R^2$= 0.64 P= 0.1 | $R^2$= -0.6 P= 0.13 |
| | Ferulic acid | 6.73 | [M+H]+ | C10H10O4 | 195.065 | $R^2$= -0.57 P= 0.08 | $R^2$= -0.57 P= 0.16 | $R^2$= 0.67 P= 0.16 |
| | Feruloyl tyramine | 7.66 | [M+H]+ | C18H19NO4 | 314.139 | $R^2$= 0.14 P= 0.78 | $R^2$= 0.14 P= 0.71 | $R^2$= -0.1 P= 0.71 |
| | Flavone base + 4O, O-MalonylHex | 9.96 | [M-H]- | C24H22O14 | 533.094 | $R^2$= -0.14 P= 0.71 | $R^2$= -0.14 P= 0.71 | $R^2$= 0.1 P= 0.78 |
| | Flavonol base + 4O, O-Hex-dHex, O-Hex | 8.64 | [M-H]- | C33H40O21 | 771.199 | $R^2$= 0.64 P= 0.1 | $R^2$= 0.64 P= 0.1 | $R^2$= -0.6 P= 0.13 |
| | Glabrol | 6.75 | [M+H]+ | C25H28O4 | 393.206 | $R^2$= -0.1 P= 0.71 | $R^2$= -0.1 P= 0.78 | $R^2$= 0.14 P= 0.78 |
| | Hyperoside | 9.32 | [M-H]- | C21H20O12 | 463.088 | $R^2$= 0.28 P= 0.49 | $R^2$= 0.28 P= 0.49 | $R^2$= -0.32 P= 0.44 |
| | irigenin | 7.26 | [M+H]+ | C18H16O8 | 361.092 | $R^2$= 0.64 P= 0.06 | $R^2$= 0.64 P= 0.1 | $R^2$= -0.75 P= 0.1 |
| | Isorhamnetin | 7.73 | [M+H]+ | C16H12O7 | 317.066 | $R^2$= 0.42 P= 0.3 | $R^2$= 0.42 P= 0.3 | $R^2$= -0.46 P= 0.3 |
| | Isorhamnetin 3-galactoside | 7.08 | [M-H]- | C22H22O12 | 477.104 | $R^2$= 0.28 P= 0.49 | $R^2$= 0.28 P= 0.49 | $R^2$= -0.32 P= 0.44 |
| | isorhamnetin-3-glucoside-4'-glucoside (Isorhamnetin 3,4'-diglucoside) | 8.98 | [M-H]- | C28H32O17 | 639.157 | $R^2$= 0.32 P= 0.44 | $R^2$= 0.32 P= 0.44 | $R^2$= -0.28 P= 0.49 |
| | isorhamnetin-3-O-glucoside | 9.57 | [M-H]- | C22H22O12 | 477.104 | $R^2$= 0.32 P= 0.44 | $R^2$= 0.32 P= 0.44 | $R^2$= -0.28 P= 0.49 |
| | Isorhamnetin-3-O-rutinoside | 9.59 | [M-H]- | C28H32O16 | 623.162 | $R^2$= 0.64 P= 0.1 | $R^2$= 0.64 P= 0.1 | $R^2$= -0.6 P= 0.13 |

| | | | | | | | | |
|---|---|---|---|---|---|---|---|---|
| | Kaempferol | 7.15 | [M+H]+ | C15H10O6 | 287.055 | $R^2$= -0.32 P= 0.71 | $R^2$= -0.32 P= 0.44 | $R^2$= 0.17 P= 0.44 |
| | Kaempferol 3-O-gentiobioside | 10.61 | [M+H]+ | C27H30O16 | 611.161 | $R^2$= -0.1 P= 0.83 | $R^2$= -0.1 P= 0.78 | $R^2$= 0.07 P= 0.78 |
| | Kaempferol 3-O-sophoroside | 8.99 | [M-H]- | C27H30O16 | 609.146 | $R^2$= 0.42 P= 0.3 | $R^2$= 0.42 P= 0.3 | $R^2$= -0.5 P= 0.26 |
| | Kaempferol-3-Glucoside-3"-Rhamnoside | 7.65 | [M+H]+ | C27H30O15 | 595.166 | $R^2$= -0.07 P= 0.78 | $R^2$= -0.07 P= 0.83 | $R^2$= 0.1 P= 0.83 |
| | Kaempferol-3-O-glucoside | 9.57 | [2M-H]- | C21H20O11 | 895.194 | $R^2$= -0.39 P= 0.39 | $R^2$= -0.39 P= 0.39 | $R^2$= 0.21 P= 0.59 |
| | kaempferol-3-O-rutinoside | 13.97 | [M-H]- | C27H30O15 | 593.151 | $R^2$= 0 P= 1 | $R^2$= 0 P= 1 | $R^2$= 0 P= 1 |
| | Kaempferol-4'-glucoside | 7.71 | [M+H]+ | C21H20O11 | 449.108 | $R^2$= -0.39 P= 0.59 | $R^2$= -0.39 P= 0.39 | $R^2$= 0.21 P= 0.39 |
| | Kaempferol-7-neohesperidoside | 9.53 | [M-H]- | C27H30O15 | 593.151 | $R^2$= 0.14 P= 0.71 | $R^2$= 0.14 P= 0.71 | $R^2$= -0.17 P= 0.71 |
| | Kukoamine B | 9.79 | [M+H]+ | C28H42N4O6 | 531.318 | $R^2$= -0.46 P= 0.1 | $R^2$= -0.46 P= 0.3 | $R^2$= 0.64 P= 0.3 |
| | Luteolin-3',7-di-O-glucoside | 9.73 | [M-H]- | C27H30O16 | 609.146 | $R^2$= 0.28 P= 0.49 | $R^2$= 0.28 P= 0.49 | $R^2$= -0.17 P= 0.71 |
| | Luteolin-7-O-glucoside | 1.26 | [M-H]- | C21H20O11 | 447.093 | $R^2$= -0.39 P= 0.38 | $R^2$= -0.39 P= 0.38 | $R^2$= 0.36 P= 0.42 |
| | methoxy-myricetin-O-hexosyl-deoxyhexoside | 7.16 | [M+H]+ | C28H32O17 | 641.171 | $R^2$= 0.32 P= 0.59 | $R^2$= 0.32 P= 0.44 | $R^2$= -0.25 P= 0.44 |
| | methyl chlorogenate | 7.1 | [M+H]+ | C17H20O9 | 369.118 | $R^2$= -0.78 P= 0.1 | $R^2$= -0.78 P= 0.04 | $R^2$= 0.64 P= 0.04 |
| | Methylophiopogonanone A | 1.43 | [M+H]+ | C19H18O6 | 343.118 | $R^2$= -0.1 P= 0.83 | $R^2$= -0.1 P= 0.78 | $R^2$= 0.07 P= 0.78 |

| | | | | | | | | |
|---|---|---|---|---|---|---|---|---|
| | N-Caffeoylputrescine | 5.28 | [M+H]+ | C13H18N2O3 | 251.139 | $R^2$= -0.21 P= 0.71 | $R^2$= -0.21 P= 0.59 | $R^2$= 0.14 P= 0.59 |
| | Panasenoside | 6.77 | [M+H]+ | C27H30O16 | 611.161 | $R^2$= -0.21 P= 0.83 | $R^2$= -0.21 P= 0.59 | $R^2$= 0.07 P= 0.59 |
| | Prunin | 7.31 | [M+H]+ | C21H22O10 | 435.129 | $R^2$= -0.07 P= 1 | $R^2$= -0.07 P= 0.83 | $R^2$= 0 P= 0.83 |
| | Quercetin | 7 | [M+H]+ | C15H10O7 | 303.05 | $R^2$= 0.71 P= 0.06 | $R^2$= 0.71 P= 0.08 | $R^2$= -0.75 P= 0.08 |
| | Quercetin 3-gentiobioside | 7 | [M+H]+ | C27H30O17 | 627.156 | $R^2$= 0.71 P= 0.06 | $R^2$= 0.71 P= 0.08 | $R^2$= -0.75 P= 0.08 |
| | Quercetin-3,4'-O-di-beta-glucopyranoside | 8.74 | [M-H]- | C27H30O17 | 625.141 | $R^2$= 0.53 P= 0.23 | $R^2$= 0.53 P= 0.23 | $R^2$= -0.46 P= 0.3 |
| | RUTOSIDE (rutin) | 9.29 | [M-H]- | C27H30O16 | 609.146 | $R^2$= -0.46 P= 0.3 | $R^2$= -0.46 P= 0.3 | $R^2$= 0.32 P= 0.44 |
| | Scoparone | 7.45 | [M+H]+ | C11H10O4 | 207.065 | $R^2$= -0.67 P= 0.06 | $R^2$= -0.67 P= 0.08 | $R^2$= 0.75 P= 0.08 |
| | Scopoletin | 6.61 | [M+H]+ | C10H8O4 | 193.05 | $R^2$= -0.92 P= 0.01 | $R^2$= -0.92 P= 0 | $R^2$= 0.85 P= 0 |
| | Tiliroside | 9.5 | [M-H]- | C30H26O13 | 593.13 | $R^2$= 0.35 P= 0.44 | $R^2$= 0.35 P= 0.44 | $R^2$= -0.39 P= 0.39 |
| | trans-5-O-Caffeoylquinic acid | 6.6 | [M-H]- | C16H18O9 | 353.088 | $R^2$= -0.4 P= 0.57 | $R^2$= -0.4 P= 0.57 | $R^2$= 0.2 P= 0.85 |
| | trans-Caffeic acid | 7.84 | [M-H]- | C9H8O4 | 179.035 | $R^2$= 0.67 P= 0.08 | $R^2$= 0.67 P= 0.08 | $R^2$= -0.64 P= 0.1 |
| | 8-Gingerol | 10.35 | [M+H]+ | C19H30O4 | 323.222 | $R^2$= -0.28 P= 0.3 | $R^2$= -0.28 P= 0.49 | $R^2$= 0.42 P= 0.49 |
| | Catechin gallate | 9.11 | [M+H]+ | C22H18O10 | 443.097 | $R^2$= -0.42 P= 0.44 | $R^2$= -0.42 P= 0.3 | $R^2$= 0.35 P= 0.3 |

| | | | | | | | | |
|---|---|---|---|---|---|---|---|---|
| | Vanillin | 7.06 | [M+H]+ | C8H8O3 | 153.055 | $R^2$= -0.32 P= 0.44 | $R^2$= -0.32 P= 0.44 | $R^2$= 0.35 P= 0.44 |
| Polycyclic Hydrocarbons | hypericin | 7.77 | [M-H]- | C30H16O8 | 503.077 | $R^2$= -0.4 P= 0.37 | $R^2$= -0.4 P= 0.37 | $R^2$= 0.48 P= 0.28 |
| Pyrimidine nucleosides | Cytidine | 1.15 | [M+H]+ | C9H13N3O5 | 244.093 | $R^2$= -0.21 P= 0.49 | $R^2$= -0.21 P= 0.59 | $R^2$= 0.28 P= 0.59 |
| Quinilines | 4-Hydroxyquinoline | 10.12 | [M-H]- | C9H7NO | 144.045 | $R^2$= 0.07 P= 0.83 | $R^2$= 0.07 P= 0.83 | $R^2$= -0.1 P= 0.78 |
| Steroids and derivatives | andrastin A | 12.09 | [M-H]- | C28H38O7 | 485.254 | $R^2$= 0.14 P= 0.71 | $R^2$= 0.14 P= 0.71 | $R^2$= -0.1 P= 0.78 |
| | Dehydroisoandrosterone sulfate | 5.69 | [M-H]- | C19H28O5S | 367.158 | $R^2$= -0.14 P= 0.71 | $R^2$= -0.14 P= 0.71 | $R^2$= 0.1 P= 0.78 |
| | Edpetiline | 6.54 | [M+H]+ | C33H53NO8 | 592.384 | $R^2$= -0.03 P= 0.71 | $R^2$= -0.03 P= 0.9 | $R^2$= 0.14 P= 0.9 |
| | Eurycomalactone | 7.91 | [M+H]+ | C19H24O6 | 349.165 | $R^2$= 0.42 P= 0.39 | $R^2$= 0.42 P= 0.3 | $R^2$= -0.39 P= 0.3 |
| | Flutamide | 7.18 | [M-H]- | C11H11F3N2O3 | 275.065 | $R^2$= 0.5 P= 0.26 | $R^2$= 0.5 P= 0.26 | $R^2$= -0.42 P= 0.3 |
| | Hydrocortisone | 9.56 | [M+H]+ | C21H30O5 | 363.217 | $R^2$= -0.07 P= 0.59 | $R^2$= -0.07 P= 0.83 | $R^2$= 0.21 P= 0.83 |
| | Hydroxyprogesterone | 10.85 | [M+H]+ | C21H30O3 | 331.227 | $R^2$= -0.25 P= 0.44 | $R^2$= -0.25 P= 0.59 | $R^2$= 0.35 P= 0.59 |
| | Mestranol | 11.37 | [M-H]- | C21H26O2 | 309.186 | $R^2$= 0.25 P= 0.59 | $R^2$= 0.25 P= 0.59 | $R^2$= -0.14 P= 0.71 |
| | Progesterone | 10.93 | [M+H]+ | C21H30O2 | 315.232 | $R^2$= 0 P= 0.9 | $R^2$= 0 P= 1 | $R^2$= -0.03 P= 1 |
| | senegenin | 10.91 | [M+H]+ | C30H45ClO6 | 537.298 | $R^2$= 0.14 P= 0.78 | $R^2$= 0.14 P= 0.71 | $R^2$= -0.1 P= 0.71 |

| | | | | | | | | |
|---|---|---|---|---|---|---|---|---|
| Terpenes and Terpenoids | 13-alpha-(21)-Epoxyeurycomanone | 6.77 | [M+H]+ | C20H24O10 | 425.144 | $R^2$= -0.39 P= 0.3 | $R^2$= -0.39 P= 0.39 | $R^2$= 0.42 P= 0.39 |
| | 2-(4-oxoquinazolin-3(4H)-yl)ethyl isobutyrate | 6.43 | [M+H]+ | C14H16N2O3 | 261.123 | $R^2$= -0.07 P= 0.78 | $R^2$= -0.07 P= 0.83 | $R^2$= 0.1 P= 0.83 |
| | 2-(furan-3-yl)-7,8-dihydroxy-6a,7,10b-trimethyl-octahydro-1H-naphtho[2,1-c]pyran-4-one | 8.77 | [M+H]+ | C20H28O5 | 349.201 | $R^2$= -0.25 P= 0.44 | $R^2$= -0.25 P= 0.59 | $R^2$= 0.35 P= 0.59 |
| | 20-deoxyingenol | 7.67 | [M+H]+ | C20H28O4 | 333.206 | $R^2$= -0.14 P= 0.71 | $R^2$= -0.14 P= 0.71 | $R^2$= 0.17 P= 0.71 |
| | Asiatic Acid | 12.92 | [M-H]- | C30H48O5 | 487.343 | $R^2$= -0.71 P= 0.08 | $R^2$= -0.71 P= 0.08 | $R^2$= 0.78 P= 0.04 |
| | Asperulosidic acid | 1.26 | [M+H]+ | C18H24O12 | 433.134 | $R^2$= 0.14 P= 0.78 | $R^2$= 0.14 P= 0.71 | $R^2$= -0.1 P= 0.71 |
| | bilobalide | 6.7 | [M+H]+ | C15H18O8 | 327.107 | $R^2$= -0.07 P= 1 | $R^2$= -0.07 P= 0.83 | $R^2$= 0 P= 0.83 |
| | Daphylloside | 6.85 | [M+H]+ | C21H36O10 | 449.238 | $R^2$= -0.1 P= 0.83 | $R^2$= -0.1 P= 0.78 | $R^2$= 0.07 P= 0.78 |
| | gamma,gamma-Dimethyallyl pyrophosphate ammonium salt | 2.49 | [M-H]- | C5H12O7P2 | 244.999 | $R^2$= -0.1 P= 0.78 | $R^2$= -0.1 P= 0.78 | $R^2$= 0.17 P= 0.71 |
| | Ganoderic acid D2 | 10.9 | [M+H]+ | C30H42O8 | 531.295 | $R^2$= -0.03 P= 0.9 | $R^2$= -0.03 P= 0.9 | $R^2$= -0.03 P= 0.9 |
| | gardenoside | 9.08 | [M+H]+ | C17H24O11 | 405.139 | $R^2$= -0.14 P= 0.59 | $R^2$= -0.14 P= 0.71 | $R^2$= 0.25 P= 0.71 |
| | geniposidic acid | 9.33 | [M-H]- | C16H22O10 | 373.114 | $R^2$= -0.71 P= 0.08 | $R^2$= -0.71 P= 0.08 | $R^2$= 0.75 P= 0.06 |
| | Ginsenoside Rf | 11.39 | [M+H]+ | C42H72O14 | 801.499 | $R^2$= -0.6 P= 0.08 | $R^2$= -0.6 P= 0.13 | $R^2$= 0.71 P= 0.13 |

| | | | | | | | | |
|---|---|---|---|---|---|---|---|---|
| | Ginsenoside Rh1 | 0.1 | [M+FA-H]- | C36H62O9 | 683.438 | $R^2$= 0.32 P= 0.44 | $R^2$= 0.32 P= 0.44 | $R^2$= -0.28 P= 0.49 |
| | Gossypol | 7.9 | [M-H]- | C30H30O8 | 517.187 | $R^2$= 0.57 P= 0.16 | $R^2$= 0.57 P= 0.16 | $R^2$= -0.64 P= 0.1 |
| | Hederagenin base + O-AcetylHex | 12.29 | [M+FA-H]- | C38H60O10 | 721.417 | $R^2$= -0.42 P= 0.3 | $R^2$= -0.42 P= 0.3 | $R^2$= 0.5 P= 0.26 |
| | Hetisine | 10.62 | [M+FA-H]- | C20H27NO3 | 374.197 | $R^2$= 0 P= 1 | $R^2$= 0 P= 1 | $R^2$= 0 P= 1 |
| | Hydroxyvalerenic Acid | 12.68 | [M-H]- | C15H22O3 | 249.15 | $R^2$= -0.07 P= 0.83 | $R^2$= -0.07 P= 0.83 | $R^2$= 0.1 P= 0.78 |
| | Lagochilin | 9.85 | [M+H]+ | C20H36O5 | 357.264 | $R^2$= 0.14 P= 1 | $R^2$= 0.14 P= 0.71 | $R^2$= 0 P= 0.71 |
| | Lucidenic acid D | 7.24 | [M+H]+ | C29H38O8 | 515.264 | $R^2$= -0.46 P= 0.3 | $R^2$= -0.46 P= 0.3 | $R^2$= 0.42 P= 0.3 |
| | Miltirone | 6.56 | [M+H]+ | C19H22O2 | 283.169 | $R^2$= -0.1 P= 0.83 | $R^2$= -0.1 P= 0.78 | $R^2$= 0.07 P= 0.78 |
| | Muurolladie-3-One | 10.11 | [M+H]+ | C15H22O | 219.174 | $R^2$= -0.14 P= 0.59 | $R^2$= -0.14 P= 0.71 | $R^2$= 0.21 P= 0.71 |
| | Ophiopogonoside A | 8.19 | [M+H]+ | C21H38O8 | 419.264 | $R^2$= -0.35 P= 0.3 | $R^2$= -0.35 P= 0.44 | $R^2$= 0.42 P= 0.44 |
| | Quillaic acid | 8.42 | [M+H]+ | C30H46O5 | 487.342 | $R^2$= 0.32 P= 0.23 | $R^2$= 0.32 P= 0.44 | $R^2$= -0.53 P= 0.44 |
| | Sylvestroside I | 6.07 | [M+H]+ | C33H48O19 | 749.286 | $R^2$= -0.32 P= 0.49 | $R^2$= -0.32 P= 0.44 | $R^2$= 0.28 P= 0.44 |
| Vitamins | D-(+)-Pantothenic acid | 6.18 | [M+H]+ | C9H17NO5 | 220.118 | $R^2$= 0.21 P= 0.9 | $R^2$= 0.21 P= 0.59 | $R^2$= -0.03 P= 0.59 |
| | Niacinamide | 1.09 | [M+H]+ | C6H6N2O | 123.055 | $R^2$= 0.03 P= 1 | $R^2$= 0.03 P= 0.9 | $R^2$= 0 P= 0.9 |

| | | | | | | | | |
|---|---|---|---|---|---|---|---|---|
| | Nicotinamide | 1.8 | [M+H]+ | C6H6N2O | 123.055 | $R^2$= -0.14<br>P= 0.59 | $R^2$= -0.14<br>P= 0.71 | $R^2$= 0.25<br>P= 0.71 |
